# Extracellular Matrix Proteomic Signatures Associate with Disease-Free Survival in Later Events of Ductal Carcinoma In Situ or Invasive Breast Cancer

**DOI:** 10.64898/2026.07.16.738889

**Authors:** Taylor S Hulahan, Laura Spruill, Bryn Gerding, Jade K Macdonald, Harrison B. Taylor, Mengjun Wang, Elizabeth Wallace, Siri H. Strand, Anand S. Mehta, Marvella E. Ford, Harikrishna Nakshatri, Jeffrey R. Marks, Robert M. Angelo, Graham A. Colditz, E. Shelley Hwang, Richard R. Drake, Robert West, Peggi M. Angel

## Abstract

**Background:** Ductal carcinoma in situ (DCIS) is a noninvasive breast lesion with variable risk of progression to invasive breast cancer (IBC). Current transcription and cell marker investigations suggest ECM decreases in later events but are limited in details of ECM proteomic composition, including post-translational modifications. We investigated whether the extracellular matrix (ECM) proteome alters with later breast events of DCIS or IBC.

**Methods:** ECM-targeted mass spectrometry imaging and liquid chromatography–tandem mass spectrometry (LC–MS/MS) were applied to ten tissue microarrays from the Resource of Archival Human Breast Tissue cohort (RAHBT). Primary DCIS specimens (n=136) were analyzed in relation to later events of DCIS (n=40) or IBC(n=30), with a mean follow-up of 192.1 months 95% CI [179.1,205.1]. Statistical modeling, survival analyses, and exploratory machine learning approaches were used to identify ECM peptide signatures associated with later events.

**Results:** Distinct ECM peptide profiles were associated with later events of DCIS or IBC. Fifteen peptides derived from fibrillar collagens (COL1A1, COL1A2, COL3A1) and elastin, showed significantly reduced abundance in patients who developed IBC. Lower expression of specific collagen peptides associated with overall 19.9% 95% CI [17.92, 21.81] decreased disease-free survival for IBC. Lower expression of these peptides was significantly associated with reduced disease-free survival (age-adjusted hazard ratio [HR] = 2.45, 95% CI: 2.33–2.57; P < 0.05). Patient-matched samples of primary DCIS, later DCIS, and later invasive breast cancer further demonstrated reduction in ECM peptide detection. Exploratory predictive modeling from patient-matched samples achieved high performance (AUROC >0.98, accuracy >93%) in distinguishing primary from later events. Following prior work in the RAHBT cohort, reduction of certain collagen peptides was also observed in primary DCIS samples from higher risk patient groups.

**Conclusions:** ECM proteomic remodeling, particularly decreases of specific collagen domains, is strongly associated with later events of DCIS and IBC. These findings highlight ECM proteome as a critical regulator of breast cancer emergence with potential as a prognosticator of risk stratification to guide clinical management of DCIS.

## Introduction

Benign epithelial neoplasms and ductal carcinoma in situ (DCIS) are preinvasive breast pathologies that are thought to increase a patient’s risk for subsequent development of invasive breast cancer (IBC). However, only a fraction of these patients with preinvasive breast pathologies will develop IBC ^1–3^. Pathological features such as nuclear grade and architectural patterns of comedo necrosis in DCIS have been associated with progression^4–7^. Consistent pathological evaluation of these features has been challenging to clinically achieve^8,9^. Therefore, genetic assays such as Oncotype DX DCIS are used to produce a score for prediction of recurrence^10^. In an independent study of 718 women using Oncotype DX DCIS with a ten-year follow up, approximately 8% of patients in the low risk for invasive disease recurrence risk group experienced IBC, whereas 15% of women in the high recurrence risk group reported local invasive disease^10^. Taken in the context of contemporary literature, preinvasive pathologies share many genetic aberrations with invasive breast cancer yet have distinct phenotypes^11,12^. Thus, the transcriptome and proteome likely drive breast cancer progression, supported by recent work showing that transcriptional and proteomic differences in stroma-associated genes particularly in the myoepithelial compartment are linked to progression^13–16^.

The stroma microenvironment undergoes discrete alterations with breast cancer progression including increases in stiffness, changes in alignment, and modifications in composition^17–26^. Notably, structural and compositional differences have been linked to clinical outcomes^27–29^, yet the regulation of the extracellular stroma proteome, particularly post-translational modifications that facilitate cell-matrix interactions, is largely unmapped across risk and recurrent breast pathologies. In the current study, the prognostic potential of extracellular matrix (ECM) protein domains is evaluated within the well-studied Resource of Archival Human Breast Tissue (RAHBT) cohort^13^, including a matched patient comparison of primary DCIS patients followed for 15.3±8.1 years for DCIS or invasive breast cancer later event. Analysis shows that specific ECM peptides are associated with poor disease-free survival with significant decreases relative to later IBC. This study represents a novel proteomic evaluation of the prognostic value of the ECM proteome in DCIS.

## Methods

### Materials

Collagenase type III (COLase3) was through collaboration with the Angel Lab and Worthington Biochemical (Lakewood, NJ, USA). PNGase F PRIME™ was purchased from N-Zyme Scientific (Doylestown, PA, USA), and Sigma Aldrich (St. Louis, MO, USA) respectively. HPLC-grade water and xylenes were obtained from Fisher Scientifics (Hampton, NH, USA). Ethanol, α-cyano-4-hydroxycinnamic acid, trifluoroacetic acid, and acetonitrile were acquired from Sigma Aldrich (St. Louis, MO, USA).

### Patient Cohort

Ten tissue microarrays were acquired from the Human Tissue Atlas Network (HTAN) as part of the Resource of Archival Human Breast Tissue, **Supplemental Table 1**). Median age of diagnosis was 54±11.2 years old. Patients identified as White (76.1%), Black (23.9%), or Pacific Islander (0.4%). Patients underwent a mastectomy (34.4%), lumpectomy with radiotherapy (47.6%), lumpectomy without radiotherapy (21.8%), or lumpectomy with unknown radiotherapy (1.4%). Median follow-up time for the cohort was 15.3±8.1 years. DCIS histotypes included cribriform, solid, comedo necrosis, and papillary. Later event was defined as having DCIS recurrence or invasive breast cancer recurrence after primary DCIS treatment. Mass spectrometry experiments were performed on tissue obtained from the RAHBT cohort in accordance with Medical University of South Carolina (MUSC) approval under IRB protocol (Pro00131060).

### Tissue Preparation for Mass Spectrometry Imaging

Tissue microarrays were deparaffinized and stained with hematoxylin (Gill 2) and eosin-y (Fisher Scientific, Hampton, NH, USA) using the manufacturer’s instructions after deglycosylation and prior to collagenase digest. Stained images were acquired using a high-resolution scanner (Nanozoomer, Hamamatsu, Japan).

Collagen proteomic imaging was completed on the same H&E-stained TMA sections after removing the coverslip^30^. Prior to peptide digest, specimens were de-glycosylated using PNGase F PRIME™ enzyme as established in previously published methods^31–33^. Specimens were processed as established in the previously published protocol^34^. A vegetable steamer was used to perform antigen retrieval for 30 minutes in 10 mM Tris HCL at pH 9. Collagenase characterized by activity assay was applied by a M5 TM Sprayer (HTX Technologies, LLC, Chapel Hill, NC, USA) in 15 passes with the following settings: 25 µL/min, 40⁰C, 10 psi, and 1200 velocity. Following enzymatic spray, tissues were incubated for five hours at 37⁰C at ≥80% humidity. Specimens were sprayed with a Matrix-Assisted Laser Desorption/Ionization (MALDI) matrix of 7 mg/mL α-cyano-4-hydroxycinnamic acid in 50% acetonitrile/1% trifluoracetic acid with 0.15 picomoles of internal standard, Glu-1-Fibrinopeptide-1. Matrix was sprayed in 14 passes with the following settings: 70 µL/min, 79⁰C, 10 psi, and 1300 velocity. After matrix application and prior to imaging, slides were dipped in cold 5 mM ammonium phosphate, monobasic and then dried in a desiccator.

### Mass Spectrometry Imaging

Tissue specimens were analyzed by a timsTOF fleX imaging mass spectrometer (Bruker, Bremen, Germany) in positive ion mode across an m/z range of 600-2500. Settings included 300 laser shots per pixel with 40 µm stepsize between pixels for tissue microarrays. Transfer time was 80.0 µs and pre-pulse storage was set to 20.0 µs. Data visualization and analysis was performed in FlexImaging v.7.5 and SCiLS Lab software 2026b Pro (Bruker Scientific, LLC, Bremen, Germany) and normalized to the internal standard. Monoisotopic peaks were exported from SCiLs Lab using the mean spectrum statistics of maximum peak intensity with the internal processing mode of peak maximum with the peak interval width set to ± 20 ppm. The natural log of peak intensities integrated over a single core was used for statistical analysis.

### Sample Preparation for LC-MS/MS

DCIS reference libraries were created from the RAHBT TMAs after imaging analysis, and augmented by proteomic analysis of whole tissue samples. Formalin-fixed paraffin-embedded slides of whole DCIS surgical specimens (n=20) were heated for one hour at 65°C. Samples underwent a series of xylenes, Carnoy, and ethanol washes. Antigen retrieval was performed in 10 mM Tris HCL at pH 9 for 30 minutes in a vegetable steamer. Tissue was scraped from slide using a razor and placed in Eppendorf tubes. Samples were sonicated to fragment tissue and increase enzymatic access. Specimen were deglycosylated using PNGase F Prime™ in overnight digest at 38°C and 450 rpm. Following deglycosylation, samples were pelleted, washed, and then underwent an overnight digest at 38°C and 450 rpm^35^ or a tryptic digest for five hours ^36^. Following the initial digest, a second digest was performed adding an additional 3 micrograms of collagenase for five hours at 38°C and 450 rpm. A C18 StageTip (Thermo Fisher Scientific, Waltham, MA, USA) followed by a ZipTip (Millipore Sigma, Burlington, MA, USA) was utilized to remove enzymes, undigested proteins and salts. TMA sections were prepared for targeted mass spectrometry using the same procedure, scraping all cores into an Eppendorf, homogenizing, and digesting prior to targeted LC-MS/MS.

### LC-MS/MS

A NanoElute 2 coupled to a PepSep XTREME column (150 µm inner diameter, 25 cm length, 1.5 µm particle size, 100 Å pore size) (Bruker Daltonics) was used for analysis. Specimens were run in triplicate (200 ng) and in a randomized order. A 36.5-minute reverse-phase gradient from 3% acetonitrile to 32% acetonitrile (0.1% formic acid) was used for peptide separation. Eluent was analyzed in a timsTOF FleX (Bruker Daltonics) in positive ion mode over a mass range of 150-1700 m/z, 0.5-1.85 1/K0 trapped ion mobility range, 0.96 s cycle time with 8 PASEF ramps per cycle, 60 s transfer time, and 12 s pre pulse storage. A calibrant filter standard at m/z = 622, 922, 1221 was for ion mobility and mass to charge ratio calibration. Every six samples, Peirce™ HeLa Protein Digest Standard was injected to evaluate system integrity and Biognosys iRT peptides were used an internal standard.

### Proteomic Data Analysis

Data was searched against a SwissProt-reviewed database of 2,366 entries from search terms “homo sapiens” AND “extracellular matrix” OR “GO:0005615” using Fragpipe v20.0 and MSFragger v3.8. Data was searched as a nonspecific digest and for the following variable post-translational modifications: M oxidation, P hydroxylation, and NQ deamidation. To assist in false discovery rate assessment, reverse decoys and contaminants were appended to the database. Peptide false discovery rate was set at 0.01. Peptide identifications investigated by the current study were combined with previous breast peptide imaging reports^37–40^ to produce a reference list of DCIS/IBC breast peptides for putative identifications by accurate mass matching. The mass spectrometry proteomics data have been deposited to the ProteomeXchange Consortium via the PRIDE^41^ partner repository with the dataset identifier PXD081119 and will be released upon publication.

### Data Analysis

Natural log transformed data was imported into Metaboanalyst v. 6.0 and the auto-scaled normalization method was performed ^42^. Heatmaps were generated in MetaboAnalyst 6.0 using the Euclidean distance and Ward clustering method. Principal component analysis used Sparse Partial Least Squares Discriminant Analysis (sPLS-DA)^43^. Log_2_fold change values and false discovery rate calculations were exported from MetaboAnalyst 6.0^44^. GraphPad Prism 10.4.1 was used to generate box plots and receiver-operator curves (ROC). ANOVA or Mann-Whitney tests (p<0.05) were used to test significance of box plots. Wilson/Brown t-tests (p<0.05) were utilized to test significance for ROC analysis. On a curated list of 479 peaks (removing isotopes and matrix peaks), linear (sp. Ordinary least squares) regression was on peptide intensities controlling for patient age using python package statsmodels^45^. Age-adjusted p-values<0.05 were considered significant. Disease free survival analysis defined samples as either high expressors if they had intensities higher than the median and low expressors if they had intensities lower than the median. Kaplan-Meier (KM) curves were used to assess disease free survival and calculated using python-based lifelines v 0.30.0^46^. Cox proportional hazard ratios (HR) were used to measure the association of disease state against all other disease states including non-progressors controlling for age as a covariate. HR plots were created with Low/High expression and visualized in forestplot 0.4.1^47^.

### Exploratory Machine Learning

Mass spectrometry imaging data from the 27 patient-matched (primary DCIS and recurrent disease) samples was normalized with z-score normalization for the benefit of computation and to avoid undue influence of predictors in a different range or scale. To identify peptide subsets with good predictive performance, wrapped feature selection techniques with support vector machine algorithm (linear kernel) were used through R package e1071 version 0.17.0. Main searching strategies to find optimal subsets of peptides used wrapper methods, sequential search, random search, and recursive feature elimination search to optimized area under the receiver operating curve (AUROC, sensitivity, specificity, accuracy>80%). Peptide predictive models were challenged with Leave-One-Out Cross-Validation (LOOCV) and Repeated-3-Fold Cross Validation (200 iterations) to explore the robustness of each predictive model. Corresponding receiver-operator curves (ROCs), derived area under curve of ROC (AUROC), and standard error (SE) were based on predictions built for each model. A predictive probability of greater or equal to 0.5 was set as the cutoff to classify primary DCIS event from recurrence event as predictive outcome. This information was summarized as a confusion matrix. Accuracy, sensitivity, specificity, positive predictive value (PPV), and negative predictive value (NPV) were calculated from the confusion matrix.

## Results

### Study Overview

This study was designed to test the hypotheses that primary DCIS shows distinguishing collagen peptide signatures from later DCIS or invasive breast cancer (IBC) events and has prognostic potential in DCIS risk stratification. Samples in the study were from 10 tissue microarrays (TMAs) in the Resource for Archival Breast Tissue (RAHBT) cohort composed of primary DCIS specimens from patients who did not experience a later breast event (n=136) and from DCIS patients who experienced either a later ipsilateral (n=38) or contralateral event (n=32) (**Table 1**, **Figure 1A**). Later breast events were defined as either IBC (n=30) or DCIS (n=40). For a subset of DCIS patients (n=27), both the primary DCIS specimens and later matched subsequent breast events were available and used in patient-matched comparisons between the primary DCIS and later breast events. TMA core assignments on the same sections were further validated for DCIS or IBC pathology by a third pathologist (MUSC after H&E staining, **Figure 1B**). Mass spectrometry imaging targeting the extracellular microenvironment was used to detect ECM (**Figure 1C**). Individual cores showed distribution patterns within and around lesions (**Figure 1D**).

**Figure 1.**
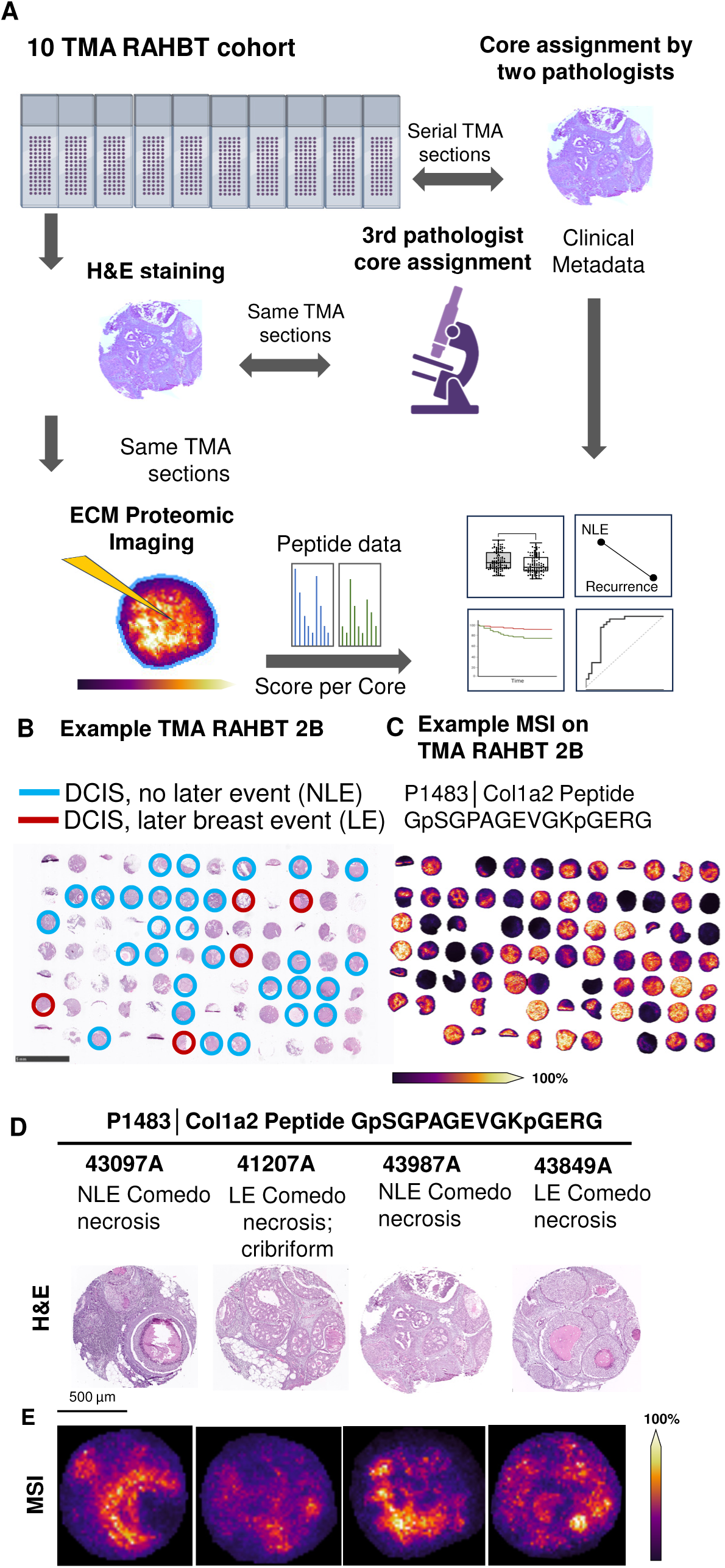
Tissue Microarrays from Resource for Archival Breast Tissue (RAHBT) on DCIS and recurrent DCIS and IBC. (A) Ten tissue microarrays (TMAs) previously evaluated for pathologies of interest were histologically stained by and subjected to pathologist evaluation for breast pathologies. After staining, microarrays were analyzed by targeted extracellular matrix proteomic imaging and peptide data from all TMAs normalized together and evaluated for peptide changes. B) example hematoxylin and eosin-stained image of TMA2B with cores marked by pathologist for inclusion in the study. C) Same TMA showing detection of a collagen peptide from the cores. D) DCIS pathology and image detection of collagen peptide from Col1a2 distributed across example cores.

**Table 1.** Characteristics of the ten tissue microarrays from the Resource for Archival Breast Tissue cohort.

| **Figure 4.**<br>Architecture Pattern |  |  |  |  |
| --- | --- | --- | --- | --- |
| Solid | 27.9% (38) | 36.4% (11) | 38.5% (15) | 25.0% (3) |
| Cribiform | 32.4% (44) | 30.3% (8) | 20.5% (8) | 25.0% (3) |
| Comedo Necrosis | 8.8% (12) | 3.0% (1) | 12.8% (5) | 0% (0) |
| Papillary | 2.2% (3) | 0% (0) | 2.6% (1) | 8.3% (1) |
| Mixed | 16.9% (23) | 12.1% (4) | 12.8% (5) | 25.0% (3) |
| Unspecified | 11.8% (16) | 18.2% (6) | 12.8% (5) | 16.7% (2) |
| **Overall Tumor Grade at Excision** |  |  |  |  |
| Grade 1 | 25.0% (34) | 21.2% (7) | 28.6% (14) | 25.0% (3) |
| Grade 2 | 33.8% (46) | 45.5% (14) | 35.7% (12) | 41.7% (5) |
| Grade 3 | 36.0% (49) | 27.3% (7) | 28.6% (15) | 25.0% (3) |
| Unknown | 5.2% (7) | 6.1% (2) | 0% (0) | 8.3% (1) |
| **Age of Primary DCIS Diagnosis** |  |  |  |  |
| Median (years) | 54 | 51 | 56 | 57 |
| Mean (years ± standard deviation) | 56 ± 11.1 | 52 ± 11.7 | 57 ± 11.5 | 56 ± 10.4 |
| **Time to Recurrence** |  |  |  |  |
| Median (months ± standard deviation) | - | 48.3 | 66.2 | 1 |
| Mean (months ± standard deviation) | - | 74.4 ± 63.7 | 82.22 ± 65.7 | 0.8 ± 0.6 |
| **Race** |  |  |  |  |
| Black | 23.5% (32) | 27.3% (9) | 19.0% (6) | 33.3% (4) |
| White | 76.4% (104) | 72.7% (21) | 78.6% (32) | 66.7% (8) |
| Pacific Islander | 0% (0) | 0% (0) | 2.4% (1) | 0% (0) |
| **Treatment** |  |  |  |  |
| Lumpectomy and radiation therapy | 42.6% (58) | 63.6% (19) | 45.2% (19) | 16.7% (2) |
| Lumpectomy without radiation therapy | 2.0% (27) | 15.2% (5) | 26.2% (10) | 25.0% (3) |
| Lumpectomy with unknown radiation status | 1.5% (2) | 0% (0) | 4.8% (1) | 0% (0) |
| Mastectomy | 36.0% (49) | 21.2% (6) | 23.8% (9) | 58.3% (7) |

### Discrete ECM Peptides Associate with IBC Later Events

To evaluate for potential prognostic peptides, all primary DCIS events were compared between all patients who experienced either DCIS or IBC recurrence (n=70) and those who did not (non-progressors; n=136) (**Figure 2A**). Samples were defined as no later event or by diagnosis with a later DCIS event (n=40) or with a later IBC event (n=30) (**Figure 2B**). Follow up time spanned 420 months with a mean 16.0 years 95% CI [14.9, 17.1] (**Figure 2C**). Mean time to event was 72.6 months 95% CI [57.7, 87.4] (**Figure 2D**). Peptides from collagen proteins were enriched in the dataset with changes in expression per category (**Figure 2E, Supplemental Table 2**). IBC later event had significantly lower levels of 15 peptides comparison to DCIS with no later events (**Figure 3A**). Peptide regions from primary fibrillar collagen Col1a2, Col1a1, and Col3a1 were the majority of the changes that also included elastin (ELN) (**Figure 3B**). For the same peptides, lower peptide expression was significantly associated with event free survival compared in IBC recurrence versus all later events (DCIS+IBC; **Figure 3C**). Disease free survival analysis linked these peptides to a 19.9% 95% CI [17.92, 21.81] decrease in survival (**Supplemental Figure 1, Supplemental Table 3 and 4**). Decreased expression associated with increased risk of invasive breast cancer, identifying 12 of the 15 peptides as conferring a significantly elevated increased risk (Cox Hazard Ratio > 1.2; age-adjusted p-values <0.05, **Figure 3D**).

**Figure 2.**
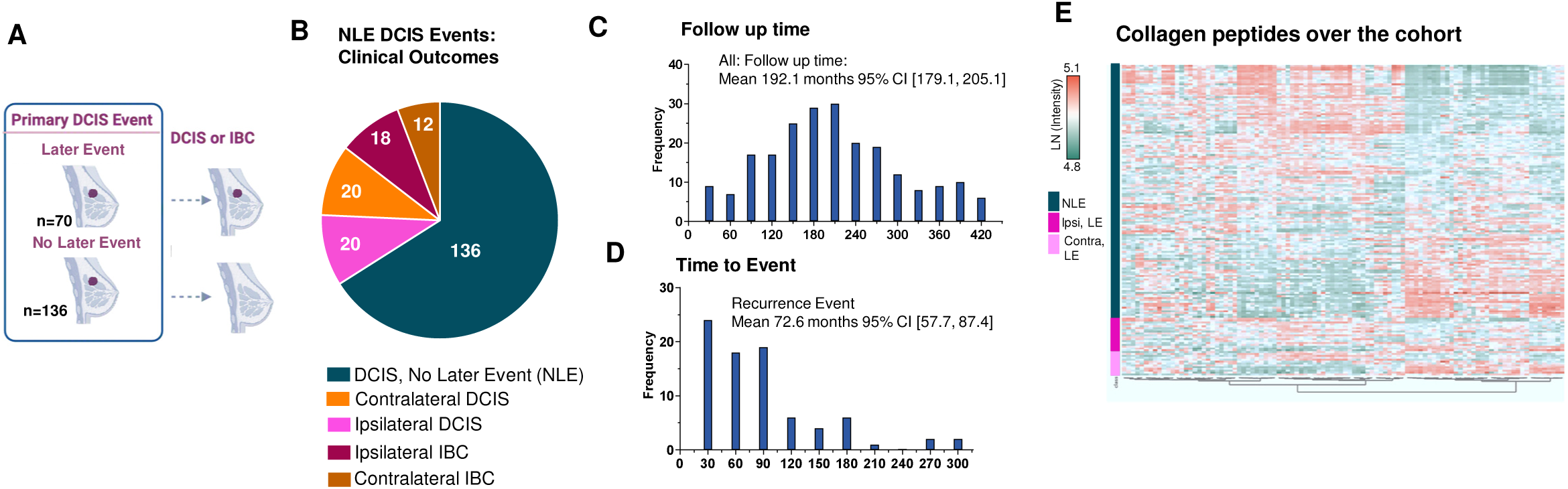
Follow-up time, and time to event, and composition. A) Samples were a cohort of DCIS NLE tissues collected from a cohort of patients who subsequently had no later event (non progressors) or had a later event either DCIS or invasive breast cancer (IBC). B) Samples with later event were distributed among ipsilateral and contralateral. C) Median follow-up for the cohort was 183.3 months [95% confidence interval [170.3, 195.5]. D) Median time to later event for the cohort was 57.8 months, 95% confidence interval [43.2, 72.4]. E) Summary heatmap of collagen peptide intensities in the study.

**Figure 3.**
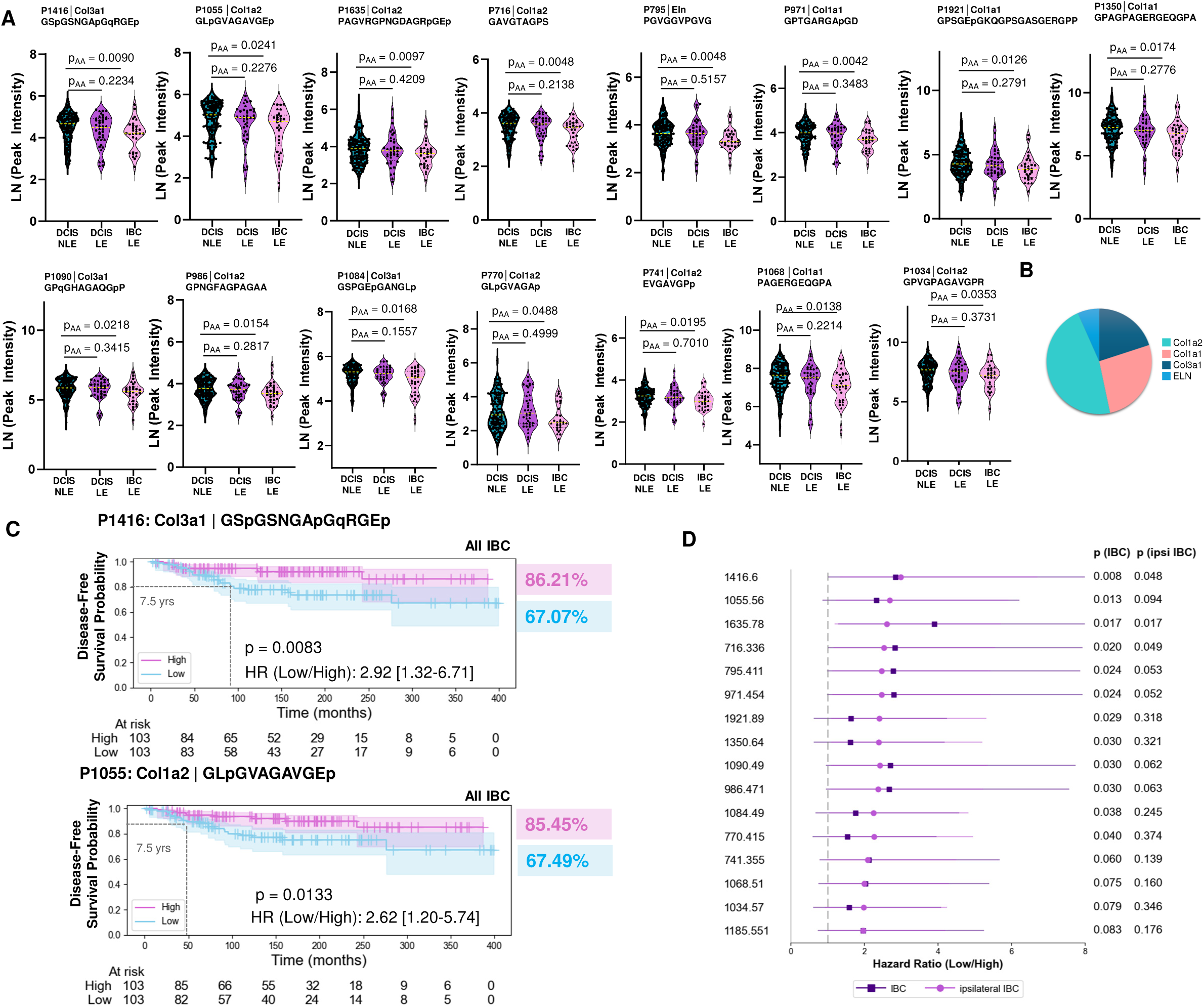
Specific extracellular matrix peptides decrease in later events of IBC and associate with decreased event free survival in a cohort from Resource for Archival Breast tissue. A) significantly altered peptides based comparing DCIS no later event, DCIS later event, and IBC later event using age-adjusted Mann-Whitney U p-value<0.05. p=hydroxylated proline; m=oxidized M; dq=deamidation NQ. Site modifications >0.95. peptides ±20 ppm. For later events of DCIS and IBC, triangle annotates contralateral event and circle indicates ipsilateral event. B) Summary of ECM proteins altered in later events. C) Peptides significantly associated with poor disease free survival, age-adjusted data, example data. D) Cox proportional hazard ratio, age-adjusted p-value <0.09 for all IBC events, hazard ratio >1.0. NLE- no later event, LE- later event, p=hydroxylated proline; m=oxidized M; nq=deamidation NQ.

### Primary DCIS is Distinguished from Later Events in Patient-Matched Samples

For a subset of the DCIS patients, patient-matched primary samples and subsequent DCIS or IBC events were evaluated, allow ECM proteomic expression from the same genetic background (**Figure 4A**). Multivariate analysis of all samples based on all peptides showed primary DCIS was distinguished from later events independent of laterality (**Figure 4B**). Primary DCIS specimens largely clustered together with higher intensity irrespective of contralateral or ipsilateral origin; lower expression was found in nearly all later events (**Figure 4C**). Fibrillar collagens Col1a1, Col1a2, Col3a1 showed reduced proportions compared to primary DCIS event with greater reduction in later events of IBC (**Figure 4D**). Paired analysis further demonstrated decreases from primary DCIS to later events of DCIS or IBC (**Figure 4E**). Peptides with decreased intensities were from Col1a1, Col1a2, Cool4a2, Col12a1, Col6a1, Col6a3, and a single HLA-DR, suggestive of board compositional changes within the microenvironment. Peptides could sensitively and specifically distinguish between primary event and later events irrespective of laterality (**Figure 4F**). Since single peptides could distinguish between recurrence and the primary DCIS event, exploratory peptide classification modeling was evaluated. Using support vector machine algorithms, models were identified consisting of six, ten, eleven, fourteen or twenty-one peptides (**Figure 4G, Supplemental Figure 2**). Models showed area under the receiver-operator curve (AUROC) greater than 98%, accuracy above 93%, specificity greater than 96%, and sensitivity over 96% in leave-out-one cross validation (LOOCV).

**Figure 4.**
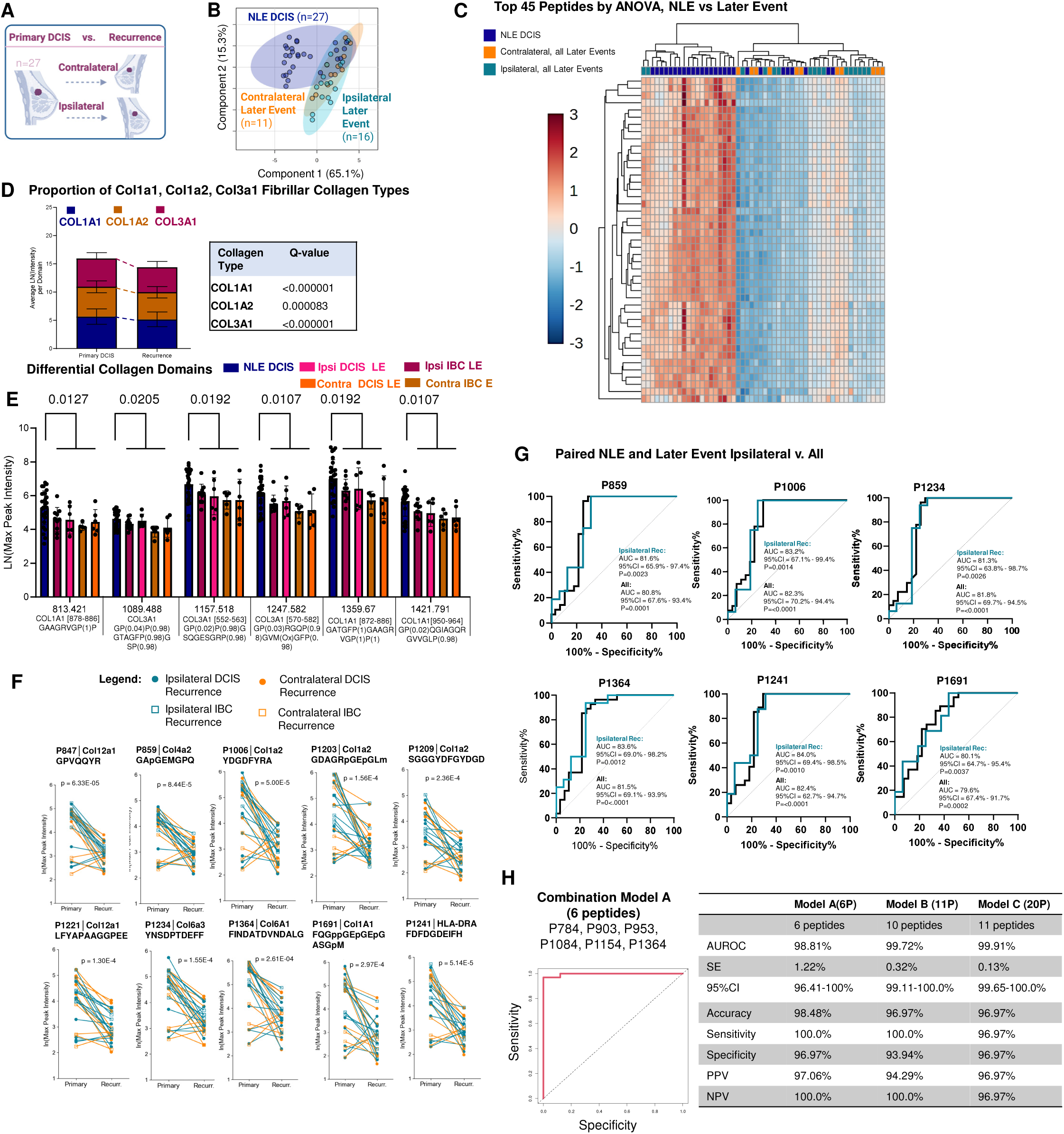
In patient-matched samples, ECM peptides differentiate primary DCIS from later events by decreases in collagen peptide intensity. (A) NLE DCIS specimens were compared to later event (DCIS or IBC) for patients with matched later event specimens (n=27). (B) Principal Components Analysis of 274 identified ECM peptide intensities demonstrates NLE DCIS event clusters apart from later event case irrespective of laterality. (C) Top ECM peptides decreases from NLE DCIS and later event cases by FDR-controlled ANOVA p-value<0.05. (D) Significant differences between NLE DCIS and later event per fibrillar collagen type proteins (q<0.05 by paired t-test with two-stage Benjamini, Krieger, and Yekutieli correction). (E) Specific peptides regions highlighted detected across all conditions and decreasing in later events of IBC. (F) Paired event comparisons in NLE and later events by FDR-adjusted paired t-test (p<3.0E-4). (G) Receiver-operator curve (ROC) analyses demonstrate discrete peptides discriminate between NLE DCIS and recurrence (AUC>80%; p<0.0001 by Wilson-Brown test). (H) Exploratory modeling in patient-matched samples demonstrates high sensitivity and specificity in Model 1, 6 peptides, Model B, 11 peptides, Model C, 20 peptides. PPP- positive predictive value; NPV- negative predictive value. p=Hydroxylated proline; m=oxidized M; nq=deamidation NQ. Site modifications >0.95. NLE= no later event; LE- later event. NLE- no later event, LE- later event. p=hydroxylated proline; m=oxidized M; nq=deamidation NQ.

### ECM Peptide Profile Differs in Populations with High Risk for Later Events

Higher risk for later events have been described as associated with African ancestry^48,49^ and changes with breast collagen fibers differ by African ancestry. Previous work in the RAHBT cohort has shown pathway differences in ipsilateral breast outcome (DCIS or IBC) after DCIS treatment by self-identified race (Black versus White)^50^, a social and demographic variable collected as part of the RAHBT cohort. A comparative analysis of ECM peptide expression was performed stratified by self-identified race as Black women (BW, n=52) and White women (WW, n=165) were defined as DCIS No later event, DCIS later events, IBC later events. Follow-up time was significantly shorter for Black patients compared to White patients (mean 16.9 years [15.68,18.2]; 13.1 years [11.2,14.9], p-value 0.0025) (**Figure 5A**). Age at diagnosis was similar between the two groups (Black, mean 56.4 years [53.6,59.3]; White 55.0 years [53.3,56.8] p-value 0.3969 (**Figure 5B**). Time to event trended as shorter for Black patients compared to White patients (Black, mean 53.9 months [28.3,76.0]; White, 78.9 [44.2,77.0],p-value 0.0787 (**Figure 5C**). Clinical outcomes of later events were similarly distributed between groups (**Figure 5D**). ECM peptide profiles revealed contrasting signatures; later events had different overall signatures compared to no later events in both groups (**Figure 5E**). Comparative analyses of ECM peptides reported significant decreases with fewer changes in primary DCIS than when comparing later events between self-identified groups (**Figure 5F**). CM collagen peptide abundance was further reduced in lesions from Black patients who experienced later breast events, and these peptides distinguished later-event status between groups (**Figure 5G, H**). Investigation of the same peptides that showed decreases in disease free survival in the overall cohort revealed that Black samples showed a much lower expression of the same ECM peptides in later events of DCIS, with an apparent rebound when compared to later events of IBC (**Figure 5I**). However, for these peptides, primary DCIS and later IBC were not significantly altered.

**Figure 5.**
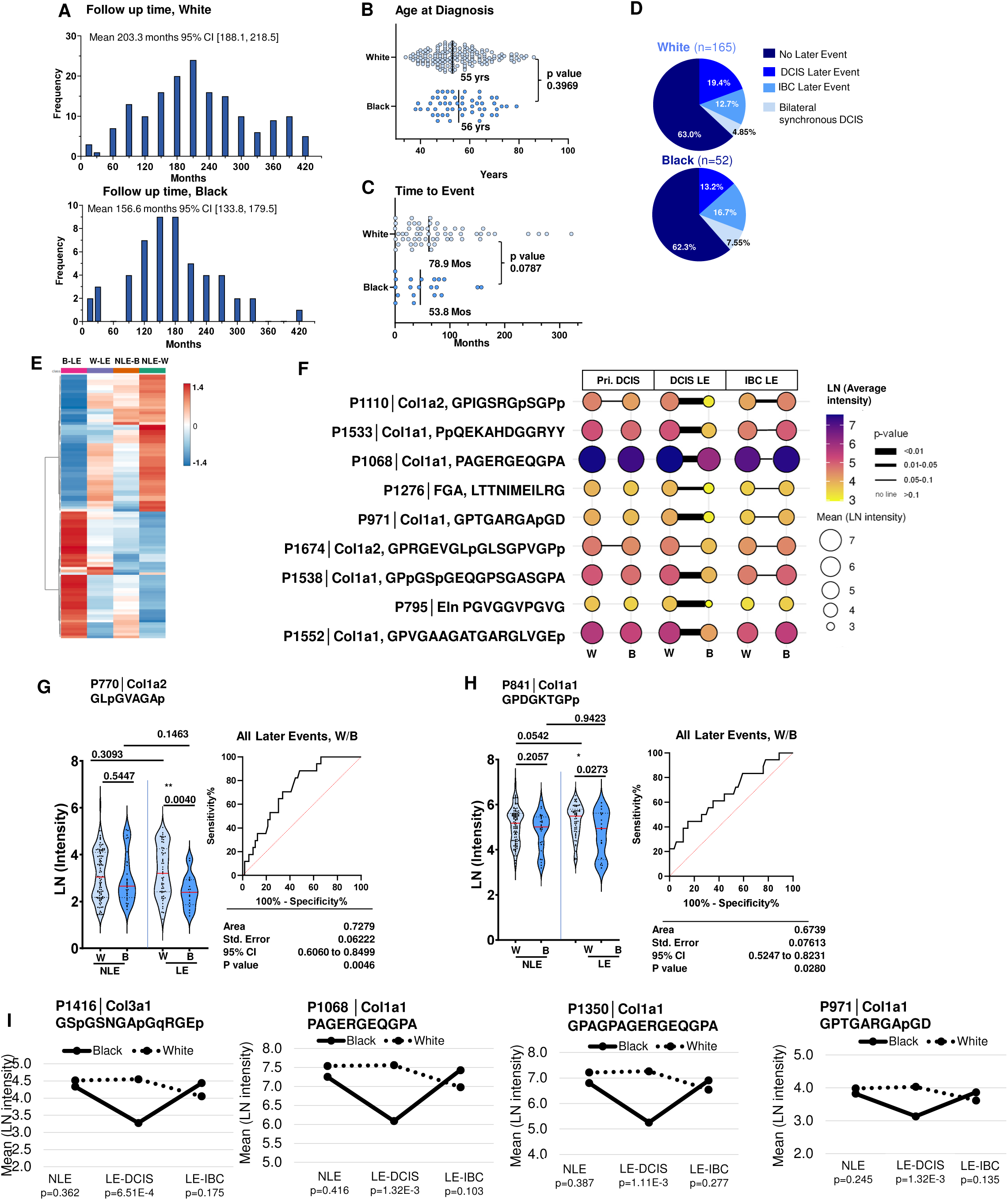
ECM proteome alters between self-identified black patients or white patients. (A) Mean follow up time shorter for black patients compared to white patients. (B) Age at diagnosis was similar for both groups. C) Time to event trended lower in black patients versus white patients. (D) Proportion of clinical outcomes was similar. (E) Average signatures demonstrate changes in no later event compared to later event. (F) Significant changes largely associate with decreases in ECM per later event of DCIS. Data is represented as natural log (LN) of detected intensity, age adjusted Mann-Whitney U test with multiple comparison-correction performed (q<0.05). (G, H) Evaluation of differential timepoints shows decreases in later events as a distinguishing features for black patients in peptides previously associated with disease free survival. (I) In collagen peptides previously associated with disease free survival in the whole cohort, samples from black patients frequently show lower expression in DCIS, Mann-Whitney U age-adjusted p-value. B- samples from black patients; W, samples from white patients; NLE-no later event; LE- later event; Pri. DCIS- primary DCIS; p=Hydroxylated proline; m=oxidized M; nq=deamidation NQ. Site modifications >0.95.

## Discussion

The data overall provides foundational information that the ECM proteome may be a source for distinguishing later events in DCIS. While DCIS itself is not considered lethal, progression to IBC increases mortality risk^51,52^. DCIS represents a crucial window for therapeutic de-escalation in low-risk patients and aggressive management in high-risk patients. Within this study, the previously published RAHBT cohort^13,14^ identified ECM peptides in primary DCIS are associated with an increased likelihood of IBC or DCIS later events. We found that low expression of discrete collagen peptide domains was associated with subsequent breast events (DCIS and IBC). This is consistent with prior proteomic and transcriptomic studies on this same cohort that showed decreased stromal desmoplasia and decreases with stromal growth and proliferation in IBC progressors, contrasting that patients with associating with signatures of ECM remodeling to have better outcomes. Further, gene expression studies on over 2,000 lesions from 145 patients have identified that loss of basal layer integrity is an early event in transitions to IBC, including COL1A2 and COL3A1^15^, which the current study confirms at the protein level.

The current study includes site specific post-translational modification of collagen hydroxylated prolines (HYPs) as markers of risk, which to our knowledge, is the first such report foundational the proteomic level. We found ECM peptides that had at least one proline that in the study showed a specific modification status (unmodified or modified); these peptides associated with decreased survival and increased risk of later event. An important concept is that collagen proteins have several hundred prolines within the triple helical region that may be dynamically and variably modified to influence biology of the tissue microenvironment yet remain largely unexplored. Discoidin domain receptor binding^53^ and integrin αβ binding^54^ domains are two well characterized collagen domains where HYP sites increase cellular affinity binding to alter functions of immune recruitment, signaling, migration, and protease interactions^53,55–60^. These domains did not change in this study, yet we have shown widespread gradients across the breast cancer^37,38^ and hepatocellular carcinoma tissue microenvironment^61^. In different cancers, discoidin domain receptors associate with increased immune exclusion, poor prognosis and increased invasion^60,62–64^ and thus are targets of therapeutic intervention^65–67^. Prolyl hydroxylases, the enzymes that catalyze proline hydroxylation, promote proliferation, invasion and metastasis in breast cancer^68–70^ yet there are limited reports evaluating site-specific modification with therapeutic potential in cancer^71,72^. Although we have recently shown that collagen peptide HYP modifications may differentially change transcriptional expression of breast epithelial tumor cell behavior^73^, it remains unknown how site-specific modifications to collagen contribute to initiating later breast events. Our findings suggest that certain HYP collagen domains are reduced in compared to the primary DCIS event. This coincides with literature showing increased desmoplasia and ECM remodeling in non-progressors^14^, and later events showing genetic decreases in ECM genes COL1A2, COL3A1, and Lumican^15^ with subsequent increases in prolyl hydroxylases P4HA1 and LEPRE1 in DCIS more closely associated with invasive breast cancer. Given that decreased hydroxylation of known cell-binding domains reduces binding affinity, changes in HYP collagen domains likely impact cell receptor interaction with collagen fibers in other ways, contributing to the variability in disease progression including potential to alter immune cell composition and recruitment. Col3a1 by antibody staining has been associated with a tumor-dormant phenotype^74^ and we found a significant association with disease free survival and higher risk to IBC in defined motifs from Col3a1 in later events. This study suggests a unique interaction with the immune response, Prior RAHBT cohort studies have shown that shifts to breast cancer are accompanied by increased myoepithelial proliferation, stromal mast cells, and CD4 T cells^13,14^.and this study support that these changes are accompanied by post-translational regulation of collagen type proteins. Further, collagen HYP sites may have a potential use in predicting later events. Collagen peptides sampled directly from tissue in performed similarly to the commercially available 12-gene Oncotype DX DCIS Score® in the assessment of ipsilateral IBC recurrence^75^. In the current study with patient-matched samples, as few as 6 peptides were found to have predictive potential, results that merit further studies on large cohorts.

Contemporary work has shown that only up to 75% of ipsilateral IBC later events are clonally related to the primary DCIS suggesting that a high proportion of IBC could be attributed to a de novo event^76^. Since ECM is produced by specific combinations of cell types during lesion evolvement, this suggests that early alterations to the collagen proteome could change the risk of subsequent breast cancers, including those induced by therapeutic interventions (i.e. radiation or endocrine therapy) or systemic changes (i.e. inflammation). It is plausible that collagen proteomes associated with higher risk develop an impaired capacity to mount an appropriate adaptive change in stromal composition and organization that maintains healthy localized immune ecologies around lesions. New studies are required to understand breast collagen proteome variations with progression from non-invasive to invasive breast cancer, but also in early-stage epithelial neoplasms and in dense normal breast tissue.

Previous work has investigated populations at high risk of IBC and mortality, focusing on sel-reported black patients as a population with high mortality risk^77,78^. This study expanded on previous work within the RAHBT cohort has found that Black women with IBC recurrence were enriched with resting fibroblasts and showed less diverse immune cell composition than Black women who do not experience later events^50^, potentially reflecting ancestry-dependent stromal variations^79^. In the current study, samples from Black women showed an ECM more similar to that of later events when compared to samples from White patients, with further decreases in certain ECM expression patterns. This suggests earlier stages of ECM proteomic variations may predispose Black women to subsequent IBC later event. Taken together, these studies underscore the need for investigating high risk populations for stromal variation that might impact early stages of disease progression and contribute to increased breast cancer aggressiveness.

### Limitations of the study

Within this study, we characterized ECM peptide signatures linked to recurrence using ten tissue microarrays from the RAHBT cohort. While tissue microarrays allow for high-throughput analysis of many patients, they capture a small fraction of the tumor microenvironment, and proteomic signatures are known be heterogeneous across breast tissue ^80^. This study included measurement of the total collagen microenvironment captured in each core due to available spatial resolution and later studies could further subdivide the stroma into compartments based on cellular composition. Genetic ancestry was not assessed, but has previously been found to be highly concordant with self-reported race in the RAHBT cohort ^50^. Our study suggested a predisposition to signatures of later events in Black women, supporting that larger cohorts need to be evaluated with molecular pathology. Clonality studies report that 75% of later breast cancer events are related to primary DCIS^81^ and this study does not consider clonality in later events. Although we report laterality, samples sizes were too small for statistical evaluation when separating ipsilateral versus contralateral. In spite of these study limitations, we found collagen proteomic signatures have potential contributions in later events and could be useful in determining patients at higher risk of later event.

## Conclusion

Based on the evidence in this study and past literature investigations^13–15^, we propose that dynamic regulation, e.g., synthesis and turnover, of the DCIS extracellular matrix allows healthy cellular trafficking around the DCIS lesion and that target ECM decreases result in later events of DCIS recurrence and invasive breast cancer. It is likely that localized inability to generate an appropriate stromal adaptive response disrupts cellular trafficking, thereby creating a permissive niche that facilitates invasive breast cancer. Exploratory machine learning on combinations of peptides from patient matched primary DCIS and later recurrence of IBC shows the potential to distinguish recurrent peptide signatures, requiring continued study in larger cohorts. Currently, two collagen domains, discoidin domain binding and integrin binding, have been characterized in the tumor microenvironment as cell-interactive post translational modifications, with significant data showing modulation of cellular function and immune distribution ^64,82–84^; this study shows potential for additional bioactive regions within the collagen structure. Further studies on larger cohorts characterizing the combined cellular and extracellular proteomic landscape of the DCIS and IBC tissue microenvironment are needed to truly understand programming in DCIS risk of recurrence and progression within the breast microenvironment. The data shown here demonstrates that the breast extracellular microenvironment holds significant variation in translational and post-translational changes with a promise of leverage for clinical management of DCIS patients.

## Acknowledgements

TSH was supported by the National Center for Advancing Translational Sciences of the National Institutes of Health under Grant Numbers TL1 TR001451 & UL1 TR001450 and NIH/NCI R01CA253460. PMA was supported by NIH/NCI R21CA263464, R21CA286287, R01CA253460; and in part by R01AG078702, P30CA138313, 5P20GM130457, the Biorepository & Tissue Analysis Shared Resource, and the Translational Science Laboratory, Hollings Cancer Center, MUSC. Tissues provided through NIH/NCI 1U2CCA233254 to RBW and ESH through the Molecular, Cellular and Tissue Characterization Unit JRM. HN was supported by DOD WH1X-WH-2010577, Susan G Komen for the Cure DRS20645418 and 1IK6BX005244 and I01BX007249 from the Department of Veterans Affairs. JKM was supported through NIH/NCI R01CA253460 and MUSC Team Science Award PR002126 funded through philanthropy and NIH/NCI P30CA138313. The Mass Spectrometry Facility and Proteomics Core are supported by NIH/NIGMS P20GM103542 with shared instrumentation NIH/OD S10OD030212 to PMA and S10OD025126 & S10OD028692 to Lauren E. Ball. The contents are solely the responsibility of the authors and do not necessarily represent the official views of the NIH.

