## Supplemental Tables and Figures for "Extracellular Matrix Proteomic Signatures Associate with Disease-Free Survival in Later Events of Ductal Carcinoma In Situ or Invasive Breast Cancer"

**Supplemental Table 1: Tissue microarrays from the Resource for Archival Breast Tissue [illegible]**

| TMA Label | Section number |
|---|---|
| TA 535 RAHBT 1B | 21 |
| TA 536 RAHBT 2B | 24 |
| TA 534 RAHBT 3B | 21 |
| TA 538 RAHBT 4B | 22 |
| TA 539 RAHBT 5B | 23 |
| TA 583 RAHBT 6B | 23 |
| TA 584 RAHBT 7B | 22 |
| TA 585 RAHBT 8B | 22 |
| TA 586 RAHBT 9B | 28 |
| TA 587 RAHBT 10B | 23 |

**Supplemental Table 2. Reference library from DCIS tissues used in the study. MUSC_DCIS refers to data generated by the current study.**

2022 DCIS Lumpectomies refers to data previously generated and publicly available reference Hulahan et al, Int J Mol Sci. 2024 Jun 19;25(12):6748, PMC11203487.
In the case of peptides in both databases, both scoring strategies from historical study (MaxQuant)and current study (MSFragger) are given. Site probabilities given in parenthesis.
All masses within a target +/-20 ppm peak interval are reported.

| Database | Gene.Names | Hyperscore or Matchscore if MaxQuant Search | Amino Acid Sequence | Peptide Length | Site Modification Probability | PPM | m/z | m/z | 43032A Contralater CP | 46145A Contralater CP | 44367A Contralater CP | 45018A Contralater CP | 44353A Contralater CP | 44007J Contralater CP | 43669A Contralater CP | 43918A Contralater CP | 45047A Contralater CP | 43674A Contralater CP | 43864A Contralater CP | 43668A Contralater CP | 43032A Contralater CP | 44253A Ipsilateral F IP | 44080D Ipsilateral F IP | 45006A Ipsilateral F IP | 4862A Ipsilateral F IP | 45043A Ipsilateral F IP | 41207A Ipsilateral F IP | 43113A Ipsilateral F IP | 43064A Ipsilateral F IP | 44334D Ipsilateral F IP | 43050A Ipsilateral F IP | 43797A Ipsilateral F IP | 41212A Ipsilateral F IP | 43124A Ipsilateral F IP | 43917A Ipsilateral F IP | 43141A Ipsilateral F IP | 42275J Ipsilateral F IP | 43162A Ipsilateral F IP | 43047A Ipsilateral F IP | 42310J Non-progre NP | 43192A Non-progre NP | 43190A Non-progre NP | 43075A Non-progre NP | 43724D Non-progre NP | 42301J Non-progre NP |
|---|---|---|---|---|---|---|---|---|---|---|---|---|---|---|---|---|---|---|---|---|---|---|---|---|---|---|---|---|---|---|---|---|---|---|---|---|---|---|---|---|---|---|---|---|---|
| MUSC_DCIS | COL3A1 | 22.5 | GPEGGKGAA | 9 | Unmodified | -2.42 | 743.368 | 743.370 | 2.933 | 1.919 | 2.292 | 2.399 | 2.498 | 2.743 | 3.279 | 3.571 | 3.582 | 3.756 | 3.560 | 3.906 | 2.933 | 2.282 | 2.618 | 2.466 | 3.672 | 3.189 | 3.662 | 3.652 | 3.131 | 3.391 | 3.287 | 4.039 | 3.440 | 3.542 | 3.776 | 3.925 | 3.519 | 3.818 | 3.782 | 2.095 | 2.308 | 2.262 | 2.300 | 2.130 | 2.177 |
| 2022 DCIS Lumpectomies | COL3A1 | 143.0 | IAGITGAR | 8 | Unmodified | -4.09 | 758.452 | 758.455 | 3.985 | 2.381 | 3.322 | 3.399 | 3.771 | 3.916 | 4.773 | 4.977 | 4.662 | 5.453 | 5.566 | 5.722 | 3.985 | 2.972 | 3.795 | 3.866 | 4.777 | 4.420 | 5.053 | 5.290 | 4.709 | 4.581 | 4.958 | 5.830 | 5.073 | 5.378 | 5.011 | 5.399 | 5.168 | 5.455 | 5.373 | 3.053 | 3.294 | 3.223 | 4.185 | 3.030 | 3.118 |
| MUSC_DCIS | COL1A2 | 20.9 | GPSGISGPP | 9 | GP(0.0070)SGISGP(0.0080)P(0.9850) | -0.64 | 784.384 | 784.384 | 3.768 | 2.588 | 3.226 | 3.501 | 3.489 | 3.745 | 4.203 | 4.395 | 4.430 | 4.558 | 4.330 | 4.493 | 3.768 | 3.321 | 3.638 | 3.587 | 4.059 | 3.959 | 4.589 | 4.504 | 4.145 | 4.199 | 4.449 | 4.466 | 4.461 | 4.237 | 4.566 | 4.384 | 4.639 | 4.701 | 4.464 | 2.872 | 3.270 | 3.462 | 3.411 | 3.353 | 3.315 |
| 2022 DCIS Lumpectomies | COL1A2 | 110.6 | GPSGISGPP | 9 | GPSGISGP(0.005)P(0.995) | -0.64 | 784.384 | | | | | | | | | | | | | | | | | | | | | | | | | | | | | | | | | | | | | | |
| MUSC_DCIS | COL1A1 | 20.2 | GAAGRVGPP | 9 | GAAGRVGP(0.5000)P(0.5000) | -4.51 | 797.426 | 797.430 | 5.961 | 3.669 | 5.029 | 5.319 | 5.245 | 6.008 | 6.561 | 6.483 | 7.019 | 7.782 | 7.829 | 7.487 | 5.961 | 4.456 | 5.107 | 5.570 | 6.765 | 6.472 | 7.249 | 6.914 | 6.875 | 6.553 | 6.743 | 7.142 | 6.990 | 7.028 | 7.017 | 7.497 | 7.394 | 7.380 | 7.639 | 4.048 | 4.500 | 5.109 | 4.983 | 5.130 | 4.716 |
| 2022 DCIS Lumpectomies | COL2A1;COL1A1 | 100.1 | GAAGRVGPP | 9 | GAAGRVGP(0.5)P(0.5) | -4.51 | 797.426 | | | | | | | | | | | | | | | | | | | | | | | | | | | | | | | | | | | | | | |
| 2022 DCIS Lumpectomies | COL1A1 | 114.2 | GPVGPVGAR | 9 | Unmodified | 2.22 | 809.463 | 809.461 | 5.309 | 1.866 | 2.995 | 3.573 | 3.482 | 4.629 | 5.805 | 6.402 | 6.786 | 7.072 | 7.680 | 7.323 | 5.309 | 2.538 | 3.664 | 3.687 | 5.513 | 6.143 | 6.145 | 6.676 | 6.524 | 6.509 | 6.547 | 6.748 | 6.690 | 6.790 | 7.111 | 7.192 | 7.127 | 7.520 | 8.154 | 2.270 | 3.108 | 3.339 | 4.356 | 3.353 | 3.562 |
| 2022 DCIS Lumpectomies | COL2A1;COL1A1 | 86.9 | GAAGRVGPP | 9 | GAAGRVGP(1)P(1) | 0.37 | 813.421 | 813.421 | 3.705 | 2.420 | 3.254 | 3.151 | 4.085 | 4.829 | 4.931 | 5.513 | 4.802 | 5.425 | 5.907 | 6.768 | 3.705 | 3.132 | 3.984 | 3.939 | 5.412 | 5.091 | 5.651 | 6.119 | 5.522 | 4.422 | 4.700 | 6.550 | 5.314 | 6.459 | 5.295 | 6.098 | 5.483 | 6.210 | 6.323 | 3.215 | 3.289 | 3.199 | 3.735 | 2.907 | 3.122 |
| MUSC_DCIS | COL1A1 | 21.4 | GPAGERGSP | 9 | GP(0.0128)AGERGSP(0.9872) | 1.78 | 843.396 | 843.394 | 7.109 | 4.551 | 5.901 | 6.109 | 6.149 | 6.794 | 7.635 | 8.304 | 8.413 | 8.287 | 8.521 | 8.495 | 7.109 | 5.413 | 6.200 | 6.142 | 7.507 | 7.794 | 7.707 | 8.062 | 7.733 | 8.114 | 7.828 | 8.035 | 7.844 | 8.073 | 8.581 | 8.405 | 8.307 | 8.340 | 8.691 | 5.075 | 5.554 | 5.718 | 5.907 | 5.749 | 5.645 |
| MUSC_DCIS | COL1A2 | 20.1 | GADGRAGVM | 9 | GADGRAGVM(1.0000) | 2.71 | 849.388 | 849.386 | 4.095 | 2.098 | 2.901 | 2.795 | 3.100 | 3.573 | 4.381 | 4.922 | 5.069 | 5.185 | 5.739 | 5.026 | 4.095 | 2.377 | 3.140 | 3.145 | 4.719 | 4.573 | 4.526 | 4.937 | 4.271 | 4.978 | 4.302 | 5.392 | 5.027 | 4.909 | 5.068 | 5.088 | 5.227 | 4.727 | 4.965 | 2.229 | 2.483 | 2.547 | 2.913 | 2.611 | 2.690 |
| 2022 DCIS Lumpectomies | COL3A1 | 110.6 | GLPGIAGPR | 9 | GPVGLQGLP(1) | -1.17 | 853.489 | 853.490 | 4.007 | 2.097 | 2.565 | 2.865 | 2.985 | 3.844 | 4.885 | 5.385 | 4.814 | 5.976 | 6.186 | 6.303 | 4.007 | 2.385 | 3.468 | 3.190 | 5.080 | 4.795 | 5.306 | 5.776 | 5.077 | 5.445 | 5.199 | 6.218 | 5.321 | 6.022 | 5.532 | 6.146 | 5.563 | 5.971 | 6.296 | 2.390 | 2.696 | 2.760 | 3.702 | 2.752 | 2.543 |
| 2022 DCIS Lumpectomies | COL1A2 | 84.2 | GPRGDQGPV | 9 | Unmodified | -0.23 | 882.443 | 882.443 | 3.298 | 1.877 | 2.279 | 2.308 | 2.347 | 2.818 | 2.990 | 4.108 | 4.555 | 4.126 | 4.693 | 4.376 | 3.298 | 2.432 | 2.343 | 2.477 | 3.446 | 3.844 | 3.401 | 3.459 | 3.332 | 4.058 | 3.624 | 3.834 | 4.115 | 3.620 | 4.078 | 4.001 | 4.614 | 4.054 | 4.509 | 1.971 | 2.166 | 2.180 | 2.029 | 2.312 | 2.230 |
| MUSC_DCIS | COL1A2 | 20.1 | GPAGKEGLR | 9 | Unmodified | -2.49 | 884.495 | 884.497 | 3.936 | 2.244 | 2.939 | 3.415 | 3.253 | 4.221 | 4.931 | 5.783 | 5.341 | 5.683 | 5.337 | 6.030 | 3.936 | 3.023 | 3.249 | 3.456 | 4.209 | 4.823 | 5.368 | 5.336 | 5.262 | 5.066 | 5.528 | 5.252 | 5.254 | 5.446 | 6.289 | 5.923 | 5.641 | 6.399 | 6.482 | 2.376 | 2.880 | 3.141 | 3.769 | 3.461 | 3.204 |
| 2022 DCIS Lumpectomies | COL1A1 | 113.1 | GVRGEPGPP | 9 | GVRGEP(1)GP(0.275)P(0.725) | 0.56 | 897.443 | 897.442 | 6.934 | 4.137 | 5.287 | 5.548 | 5.592 | 6.505 | 6.444 | 7.322 | 8.140 | 8.016 | 8.372 | 8.250 | 6.934 | 4.938 | 5.894 | 5.794 | 7.141 | 7.476 | 7.435 | 7.478 | 7.278 | 7.497 | 7.345 | 7.853 | 8.076 | 7.648 | 7.549 | 7.808 | 8.409 | 7.413 | 7.934 | 4.477 | 5.151 | 5.388 | 5.559 | 5.301 | 5.209 |
| MUSC_DCIS | COL5A1 | 20.3 | GPDGLRGLP | 9 | GP(0.0121)DGLRGLP(0.9879) | 0.89 | 897.479 | 897.478 | 3.759 | 1.926 | 2.474 | 2.831 | 2.837 | 3.430 | 4.894 | 5.833 | 4.884 | 5.765 | 6.231 | 5.769 | 3.759 | 2.387 | 3.575 | 2.929 | 4.080 | 4.412 | 5.026 | 5.656 | 5.039 | 4.713 | 5.178 | 5.891 | 5.069 | 5.514 | 5.437 | 5.505 | 5.451 | 5.893 | 6.238 | 2.341 | 2.528 | 2.778 | 4.274 | 2.734 | 2.633 |
| 2022 DCIS Lumpectomies | COL1A2 | 102.8 | GPRGEVGLP | 9 | GPRGEVGLP(1) | 0.89 | 897.479 | | | | | | | | | | | | | | | | | | | | | | | | | | | | | | | | | | | | | | |
| MUSC_DCIS | COL1A1 | 21.6 | GQRGVVGLP | 9 | GQRGVVGLP(1.0000) | -0.67 | 898.510 | 898.511 | 3.752 | 2.052 | 2.808 | 3.148 | 2.920 | 3.740 | 4.501 | 6.000 | 5.058 | 5.363 | 5.136 | 4.869 | 3.752 | 2.659 | 3.354 | 3.145 | 4.190 | 4.719 | 4.894 | 5.202 | 4.955 | 5.025 | 4.808 | 5.151 | 4.689 | 5.012 | 5.076 | 5.316 | 5.098 | 5.791 | 5.952 | 2.333 | 2.949 | 3.020 | 3.440 | 3.125 | 2.754 |
| 2022 DCIS Lumpectomies | COL1A1 | 97.1 | GQRGVVGLP | 9 | GQRGVVGLP(1) | -0.56 | 898.511 | | | | | | | | | | | | | | | | | | | | | | | | | | | | | | | | | | | | | | |
| 2022 DCIS Lumpectomies | COL3A1 | 93.2 | AGAPGERGPP | 10 | AGAP(1)GERGP(0.31)P(0.69) | -1.81 | 940.448 | 940.450 | 3.475 | 2.383 | 2.878 | 2.980 | 2.987 | 3.560 | 3.845 | 4.486 | 4.251 | 4.787 | 4.813 | 5.316 | 3.475 | 2.737 | 2.950 | 3.167 | 4.175 | 3.966 | 4.240 | 4.197 | 4.022 | 4.324 | 3.985 | 4.733 | 4.352 | 4.532 | 4.684 | 4.748 | 4.676 | 5.123 | 5.123 | 2.457 | 2.593 | 2.554 | 2.665 | 2.684 | 2.617 |
| 2022 DCIS Lumpectomies | COL1A2 | 89.0 | GPRGEVGLPG | 10 | GPRGEVGLP(1)G | -0.73 | 954.500 | 954.501 | 5.447 | 2.673 | 3.787 | 4.381 | 4.504 | 4.801 | 5.937 | 5.806 | 7.209 | 6.402 | 6.831 | 7.500 | 5.447 | 4.185 | 4.667 | 4.492 | 5.569 | 6.397 | 5.563 | 5.674 | 5.602 | 6.498 | 6.398 | 6.032 | 6.488 | 6.290 | 6.917 | 6.490 | 7.236 | 6.796 | 6.932 | 3.400 | 3.906 | 3.981 | 4.356 | 4.141 | 4.261 |
| MUSC_DCIS | COL3A1 | 22.2 | IAGITGARGLA | 11 | Unmodified | 1.50 | 999.595 | 999.593 | 3.038 | 2.875 | 2.764 | 2.959 | 3.242 | 3.317 | 4.590 | 4.928 | 3.970 | 4.897 | 4.608 | 4.834 | 3.038 | 2.926 | 3.436 | 3.404 | 3.974 | 4.045 | 4.786 | 4.972 | 4.404 | 4.193 | 4.474 | 5.030 | 4.277 | 5.047 | 4.308 | 4.866 | 4.622 | 5.518 | 5.281 | 2.644 | 3.062 | 3.037 | 3.961 | 2.532 | 2.713 |
| 2022 DCIS Lumpectomies | COL3A1 | 126.4 | IAGITGARGLA | 11 | Unmodified | 1.50 | 999.595 | | | | | | | | | | | | | | | | | | | | | | | | | | | | | | | | | | | | | | |
| MUSC_DCIS | COL1A1 | 22.3 | GARGNDGATGAA | 12 | Unmodified | -3.15 | 1017.471 | 1017.474 | 4.673 | 2.606 | 3.390 | 3.449 | 3.563 | 4.172 | 4.608 | 6.086 | 6.061 | 5.399 | 5.880 | 5.630 | 4.673 | 2.967 | 4.042 | 3.458 | 4.890 | 5.468 | 5.143 | 5.030 | 4.717 | 5.644 | 5.041 | 5.367 | 5.519 | 5.088 | 5.595 | 5.417 | 5.826 | 5.215 | 5.402 | 2.883 | 3.344 | 3.406 | 3.639 | 3.181 | 3.242 |
| MUSC_DCIS | COL1A1 | 21.4 | GQRGERGFP | 9 | GQRGERGFP(1.0000) | -4.32 | 1019.502 | 1019.506 | 5.331 | 2.291 | 2.719 | 2.869 | 2.735 | 4.019 | 4.118 | 4.908 | 6.829 | 6.522 | 7.387 | 6.931 | 5.331 | 2.314 | 2.610 | 3.267 | 5.566 | 5.989 | 5.294 | 5.587 | 5.755 | 6.334 | 6.265 | 6.250 | 6.552 | 6.107 | 5.881 | 6.247 | 7.094 | 6.267 | 7.349 | 2.308 | 4.104 | 2.593 | 2.992 | 3.079 | 3.465 |
| MUSC_DCIS | COL6A3 | 22.4 | LAFALDTSEG | 10 | Unmodified | -0.68 | 1023.499 | 1023.500 | 3.608 | 1.991 | 2.612 | 2.729 | 2.887 | 3.166 | 4.454 | 3.994 | 5.198 | 4.164 | 4.459 | 4.998 | 3.608 | 2.396 | 2.615 | 3.011 | 3.630 | 4.332 | 3.332 | 3.569 | 3.691 | 4.561 | 4.054 | 3.784 | 4.311 | 4.371 | 5.081 | 4.298 | 5.035 | 4.791 | 4.804 | 2.185 | 2.674 | 2.383 | 2.325 | 2.678 | 3.034 |
| MUSC_DCIS | COL1A2 | 21.0 | GATGDRGEAGAA | 12 | Unmodified | 1.36 | 1032.470 | 1032.469 | 4.421 | 2.991 | 3.622 | 3.681 | 3.895 | 3.988 | 4.897 | 6.055 | 5.723 | 5.149 | 5.564 | 5.403 | 4.421 | 3.695 | 3.739 | 3.897 | 4.724 | 5.061 | 4.760 | 4.818 | 4.388 | 5.230 | 4.672 | 5.024 | 5.188 | 4.863 | 5.271 | 5.118 | 5.580 | 5.426 | 5.324 | 2.919 | 3.248 | 3.323 | 3.375 | 3.261 | 3.508 |
| MUSC_DCIS | COL1A1 | 21.7 | GPVGPVGARGPA | 12 | Unmodified | 2.03 | 1034.574 | 1034.572 | 6.298 | 4.424 | 5.570 | 6.220 | 5.849 | 6.481 | 7.536 | 7.893 | 7.361 | 7.795 | 7.032 | 8.382 | 6.298 | 5.711 | 5.847 | 6.088 | 6.623 | 7.023 | 7.412 | 7.012 | 7.203 | 7.642 | 7.651 | 7.089 | 7.382 | 7.251 | 8.413 | 7.807 | 8.041 | 8.955 | 8.940 | 4.850 | 5.296 | 5.800 | 6.181 | 5.938 | 6.071 |
| MUSC_DCIS | COL1A2 | 25.8 | GPAGSRGDGGPP | 12 | GP(0.0068)AGSRGDGGP(0.0075)P(0.9857) | -0.58 | 1040.475 | 1040.476 | 6.384 | 3.530 | 4.897 | 5.090 | 5.085 | 6.145 | 6.445 | 6.999 | 7.748 | 7.847 | 8.057 | 8.093 | 6.384 | 4.527 | 5.131 | 5.255 | 6.617 | 7.001 | 7.211 | 7.200 | 7.175 | 7.177 | 7.251 | 7.524 | 7.651 | 7.401 | 7.586 | 7.722 | 8.050 | 7.725 | 8.132 | 3.816 | 4.571 | 4.892 | 5.268 | 5.025 | 4.540 |
| 2022 DCIS Lumpectomies | COL5A1;COL11A1 | 97.1 | GPPGTMLMLP | 10 | GP(1)P(1)GTM(1)LM(1)LP | 1.39 | 1077.496 | 1077.494 | 3.462 | 2.462 | 3.332 | 3.336 | 3.735 | 3.784 | 4.594 | 5.125 | 4.208 | 4.568 | 4.353 | 4.801 | 3.462 | 3.122 | 3.291 | 3.745 | 4.176 | 4.065 | 4.395 | 4.392 | 4.262 | 4.489 | 4.365 | 4.364 | 4.205 | 4.711 | 4.714 | 5.157 | 4.477 | 5.679 | 5.427 | 2.611 | 2.895 | 2.892 | 2.929 | 3.062 | 3.203 |
| MUSC_DCIS | COL3A1 | 26.9 | GPSGSPGKDGPP | 12 | GP(0.0189)SGSP(0.9717)GKDGP(0.0385)P(0 | -2.49 | 1084.490 | 1084.493 | 4.406 | 3.180 | 3.986 | 4.007 | 4.280 | 4.597 | 4.914 | 5.278 | 5.131 | 5.392 | 5.138 | 5.556 | 4.406 | 4.103 | 4.213 | 4.212 | 5.219 | 4.800 | 5.337 | 5.319 | 4.767 | 4.863 | 4.854 | 5.607 | 5.059 | 5.078 | 5.474 | 5.502 | 5.186 | 5.353 | 5.518 | 3.514 | 3.848 | 3.772 | 3.739 | 3.664 | 3.710 |
| MUSC_DCIS | COL3A1 | 23.9 | GPPGTAGFPGSP | 12 | GP(0.0460)P(0.9847)GTAGFP(0.9847)GSP(0 | -3.12 | 1089.485 | 1089.488 | 3.805 | 3.278 | 3.943 | 3.978 | 4.224 | 3.978 | 4.776 | 4.755 | 4.616 | 5.124 | 5.318 | 5.102 | 3.805 | 4.345 | 3.795 | 4.050 | 4.483 | 4.075 | 4.313 | 4.647 | 4.437 | 4.474 | 4.158 | 5.170 | 4.677 | 4.763 | 4.612 | 5.022 | 4.865 | 5.130 | 4.959 | 2.991 | 3.260 | 3.795 | 2.896 | 3.586 | 3.411 |
| 2022 DCIS Lumpectomies | COL3A1 | 187.6 | GPPGTAGFPGSP | 12 | GP(0.001)P(0.999)GTAGFP(1)GSP(1) | -3.12 | 1089.485 | | | | | | | | | | | | | | | | | | | | | | | | | | | | | | | | | | | | | | |
| MUSC_DCIS | COL3A1 | 20.9 | GAAGTPGLQGMP | 12 | GAAGTP(0.9851)GLQGM(0.0298)P(0.9851) | 0.00 | 1089.488 | | | | | | | | | | | | | | | | | | | | | | | | | | | | | | | | | | | | | | |
| MUSC_DCIS | COL3A1 | 21.0 | GPPGTSGHPGSP | 12 | GP(0.0375)P(0.9875)GTSGHP(0.9875)GSP(0 | -4.47 | 1095.470 | 1095.475 | 3.886 | 2.865 | 3.432 | 3.605 | 3.652 | 3.943 | 4.554 | 4.906 | 4.495 | 4.884 | 4.780 | 4.664 | 3.886 | 3.487 | 3.645 | 3.809 | 4.760 | 4.135 | 4.752 | 4.570 | 4.396 | 4.670 | 4.335 | 4.913 | 4.489 | 4.546 | 4.844 | 5.100 | 4.500 | 4.749 | 4.779 | 2.901 | 3.079 | 3.254 | 3.046 | 3.306 | 3.253 |
| MUSC_DCIS | COL1A1 | 21.6 | GIAGAPGFPGAR | 12 | GIAGAP(1.0000)GFP(1.0000)GAR | 0.73 | 1102.564 | 1102.563 | 3.339 | 1.809 | 2.009 | 2.265 | 2.024 | 2.619 | 2.960 | 3.678 | 5.017 | 4.412 | 5.434 | 4.825 | 3.339 | 2.013 | 2.448 | 2.374 | 3.412 | 4.260 | 3.771 | 4.053 | 3.673 | 4.486 | 3.724 | 4.303 | 4.657 | 4.063 | 4.201 | 4.243 | 5.480 | 4.068 | 4.703 | 1.849 | 2.165 | 2.062 | 2.722 | 2.283 | 2.059 |
| 2022 DCIS Lumpectomies | COL1A1 | 127.0 | GIAGAPGFPGAR | 12 | GIAGAP(1)GFP(1)GAR | 0.73 | 1102.564 | | | | | | | | | | | | | | | | | | | | | | | | | | | | | | | | | | | | | | |
| MUSC_DCIS | COL1A2 | 20.9 | LHGEFGLPGPA | 11 | LHGEFGLP(0.9863)GP(0.0137)A | 4.23 | 1110.558 | 1110.553 | 2.994 | 2.608 | 2.988 | 3.053 | 3.668 | 4.407 | 4.381 | 5.097 | 4.514 | 4.038 | 3.991 | 5.818 | 2.994 | 3.545 | 3.833 | 3.574 | 3.838 | 4.367 | 4.203 | 4.411 | 4.180 | 4.358 | 3.889 | 4.466 | 4.075 | 4.798 | 4.480 | 4.608 | 4.514 | 5.901 | 5.607 | 2.609 | 2.938 | 2.662 | 2.684 | 2.752 | 2.949 |
| MUSC_DCIS | COL1A1 | 20.6 | GPSGEPGKQGPS | 12 | GP(0.0066)SGEP(0.9792)GKQGP(0.0143)S | 0.81 | 1113.517 | 1113.516 | 4.353 | 2.587 | 3.202 | 3.469 | 3.493 | 4.139 | 4.728 | 5.328 | 5.834 | 5.640 | 5.726 | 5.928 | 4.353 | 2.984 | 3.786 | 3.502 | 4.881 | 5.068 | 5.013 | 5.077 | 4.833 | 5.319 | 4.945 | 5.536 | 5.303 | 5.213 | 5.802 | 5.426 | 5.811 | 5.168 | 5.236 | 2.814 | 3.140 | 3.081 | 3.562 | 3.254 | 3.177 |
| MUSC_DCIS | COL2A1 | 20.6 | GPSGEPGKQGAP | 12 | GP(0.0282)SGEP(0.9859)GKQGAP(0.9859) | 0.81 | 1113.517 | | | | | | | | | | | | | | | | | | | | | | | | | | | | | | | | | | | | | | |
| 2022 DCIS Lumpectomies | COL2A1 | 179.4 | GPSGEPGKQGAP | 12 | GPSGEP(1)GKQGAP(1) | 0.81 | 1113.517 | | | | | | | | | | | | | | | | | | | | | | | | | | | | | | | | | | | | | | |
| MUSC_DCIS | COL4A1 | 20.4 | GPPGIPGFDGAP | 12 | GP(0.0534)P(0.9661)GIP(0.9674)GFDGAP(0. | 4.49 | 1113.521 | | | | | | | | | | | | | | | | | | | | | | | | | | | | | | | | | | | | | | |
| MUSC_DCIS | COL1A1 | 20.8 | GAKGDRGETGPA | 12 | Unmodified | -0.99 | 1115.544 | 1115.545 | 6.169 | 3.383 | 4.613 | 4.708 | 4.742 | 5.663 | 6.263 | 6.849 | 7.444 | 7.495 | 7.584 | 7.614 | 6.169 | 3.943 | 4.919 | 4.763 | 6.351 | 6.719 | 6.893 | 6.967 | 6.529 | 6.790 | 6.797 | 7.351 | 7.344 | 7.110 | 7.323 | 7.254 | 7.686 | 7.089 | 7.167 | 3.877 | 4.493 | 4.640 | 5.031 | 4.474 | 4.267 |
| 2022 DCIS Lumpectomies | COL1A1 | 112.9 | GAKGDRGETGPA | 12 | Unmodified | -0.99 | 1115.544 | | | | | | | | | | | | | | | | | | | | | | | | | | | | | | | | | | | | | | |
| MUSC_DCIS | COL1A1 | 20.9 | GAEGSPGRDGSP | 12 | GAEGSP(1.0000)GRDGSP(1.0000) | -0.27 | 1118.471 | 1118.471 | 6.068 | 4.023 | 4.762 | 4.657 | 4.574 | 5.512 | 6.169 | 6.210 | 7.187 | 7.098 | 7.241 | 7.234 | 6.068 | 4.162 | 5.315 | 4.586 | 6.333 | 6.640 | 6.631 | 6.820 | 6.288 | 6.744 | 6.353 | 7.074 | 7.097 | 6.919 | 7.163 | 6.976 | 7.423 | 6.342 | 6.680 | 4.170 | 4.533 | 4.668 | 5.199 | 4.570 | 4.487 |
| MUSC_DCIS | COL1A1 | 21.0 | GPAGRPGEVGPP | 12 | GP(0.0105)AGRP(0.7554)GEVGP(0.6170)P(0 | -2.05 | 1122.554 | 1122.556 | 5.868 | 2.801 | 4.131 | 4.232 | 4.056 | 5.558 | 5.634 | 6.143 | 7.450 | 7.397 | 7.591 | 7.740 | 5.868 | 3.343 | 4.274 | 4.433 | 6.220 | 6.737 | 6.406 | 6.642 | 6.657 | 7.018 | 6.708 | 6.904 | 7.091 | 7.083 | 7.458 | 7.337 | 7.661 | 7.074 | 7.826 | 3.109 | 3.780 | 4.069 | 4.186 | 4.217 | 4.021 |
| 2022 DCIS Lumpectomies | COL1A1 | 87.3 | GPAGRPGEVGPP | 12 | GPAGRP(1)GEVGP(0.06)P(0.94) | -2.05 | 1122.554 | | | | | | | | | | | | | | | | | | | | | | | | | | | | | | | | | | | | | | |
| MUSC_DCIS | COL1A1 | 23.1 | GKSGDRGETGPA | 12 | Unmodified | 0.71 | 1131.539 | 1131.538 | 5.314 | 3.008 | 3.885 | 3.896 | 3.968 | 4.606 | 5.168 | 5.770 | 6.333 | 6.300 | 6.520 | 6.430 | 5.314 | 3.400 | 4.219 | 3.787 | 5.730 | 5.712 | 5.762 | 5.715 | 5.202 | 5.802 | 5.501 | 6.358 | 6.263 | 5.728 | 6.063 | 5.938 | 6.469 | 5.574 | 5.772 | 3.458 | 3.901 | 3.940 | 4.100 | 3.726 | 3.481 |
| MUSC_DCIS | COL1A1 | 21.7 | GPPGESGREGAP | 12 | GP(0.0283)P(0.9859)GESGREGAP(0.9859) | -2.54 | 1142.507 | 1142.510 | 6.138 | 3.901 | 4.927 | 4.990 | 4.794 | 5.923 | 6.144 | 7.359 | 7.529 | 7.546 | 8.045 | 7.886 | 6.138 | 4.241 | 4.959 | 5.084 | 6.515 | 6.901 | 6.702 | 7.019 | 6.935 | 7.215 | 6.825 | 7.235 | 7.246 | 7.337 | 7.664 | 7.594 | 7.877 | 7.250 | 7.864 | 4.283 | 4.536 | 4.649 | 4.944 | 4.906 | 4.427 |
| 2022 DCIS Lumpectomies | COL1A1 | 108.2 | GPPGESGREGAP | 12 | GP(0.003)P(0.997)GESGREGAP(1) | -2.54 | 1142.507 | | | | | | | | | | | | | | | | | | | | | | | | | | | | | | | | | | | | | | |
| MUSC_DCIS | COL1A1 | 20.8 | GAPGDRGEPGPP | 12 | GAP(0.9825)GDRGEP(0.9825)GP(0.0525)P(0 | -4.16 | 1154.507 | 1154.512 | 4.137 | 2.792 | 3.464 | 3.807 | 3.771 | 4.164 | 4.409 | 6.657 | 5.489 | 4.799 | 4.965 | 5.253 | 4.137 | 3.333 | 3.654 | 3.916 | 3.964 | 4.776 | 4.208 | 4.519 | 4.327 | 4.812 | 4.626 | 4.327 | 4.778 | 4.609 | 4.924 | 4.822 | 5.371 | 5.294 | 5.516 | 2.896 | 3.250 | 3.404 | 3.055 | 3.647 | 3.399 |
| 2022 DCIS Lumpectomies | COL1A1 | 98.0 | GAPGDRGEPGPP | 12 | GAP(0.333)GDRGEP(0.333)GP(0.026)P(0.30 | -4.16 | 1154.507 | | | | | | | | | | | | | | | | | | | | | | | | | | | | | | | | | | | | | | |
| MUSC_DCIS | COL3A1 | 21.1 | GPPGSQGESGRP | 12 | GP(0.0255)P(0.9872)GSQGESGRP(0.9872) | 0.00 | 1157.518 | 1157.518 | 6.004 | 3.258 | 4.456 | 4.557 | 4.822 | 5.804 | 5.693 | 6.303 | 6.695 | 7.528 | 7.384 | 7.792 | 6.004 | 3.948 | 4.796 | 4.920 | 6.723 | 6.328 | 6.736 | 6.581 | 6.317 | 6.626 | 6.416 | 7.522 | 6.976 | 7.003 | 6.942 | 7.422 | 7.041 | 6.921 | 7.431 | 3.648 | 3.919 | 4.283 | 4.389 | 4.407 | 3.875 |
| MUSC_DCIS | COL3A1 | 20.2 | GMPGERGGLGSP | 12 | GM(1.0000)P(1.0000)GERGGLGSP(1.0000) | -1.20 | 1162.516 | 1162.517 | 3.899 | 2.362 | 2.963 | 2.963 | 3.003 | 3.634 | 4.036 | 4.667 | 4.736 | 5.185 | 5.613 | 5.561 | 3.899 | 2.618 | 3.106 | 3.115 | 4.816 | 4.398 | 4.435 | 4.735 | 4.244 | 4.866 | 4.399 | 5.565 | 4.942 | 5.073 | 4.890 | 5.257 | 5.067 | 5.012 | 5.422 | 2.577 | 2.617 | 2.787 | 2.761 | 2.746 | 2.795 |
| MUSC_DCIS | COL3A1 | 22.3 | GSPGERGETGPP | 12 | GSP(0.9844)GERGETGP(0.0313)P(0.9843) | -4.52 | 1172.518 | 1172.523 | 6.370 | 5.027 | 5.760 | 5.873 | 6.163 | 6.632 | 7.322 | 8.004 | 7.525 | 8.172 | 7.967 | 8.176 | 6.370 | 5.696 | 6.069 | 6.308 | 7.187 | 7.021 | 7.710 | 7.167 | 7.242 | 7.614 | 7.092 | 7.989 | 7.609 | 7.295 | 8.176 | 7.964 | 7.589 | 7.982 | 8.165 | 4.882 | 5.132 | 5.531 | 5.830 | 5.483 | 5.463 |
| MUSC_DCIS | COL1A1 | 21.9 | GARGQAGVMGFP | 12 | GARGQAGVM(1.0000)GFP(1.0000) | -3.14 | 1179.557 | 1179.561 | 6.217 | 4.024 | 4.690 | 5.220 | 5.043 | 5.817 | 6.309 | 6.730 | 7.518 | 7.601 | 7.685 | 7.849 | 6.217 | 5.089 | 5.012 | 5.382 | 6.466 | 6.747 | 6.941 | 6.655 | 6.988 | 7.218 | 7.100 | 7.087 | 7.516 | 7.011 | 7.552 | 7.433 | 8.116 | 8.218 | 8.437 | 3.963 | 4.989 | 5.042 | 4.898 | 5.181 | 5.324 |
| 2022 DCIS Lumpectomies | COL1A1 | 92.4 | GARGQAGVMGFP | 12 | GARGQAGVM(1)GFP(1) | -2.97 | 1179.558 | | | | | | | | | | | | | | | | | | | | | | | | | | | | | | | | | | | | | | |
| MUSC_DCIS | COL1A1 | 21.8 | GADGSPGKDGVRG | 13 | GADGSP(1.0000)GKDGVRG | 0.25 | 1188.560 | 1188.560 | 4.218 | 2.446 | 3.068 | 3.009 | 2.745 | 3.122 | 4.392 | 3.825 | 5.880 | 4.715 | 5.182 | 4.565 | 4.218 | 2.527 | 3.187 | 2.842 | 4.182 | 5.083 | 3.625 | 3.277 | 3.427 | 5.130 | 4.625 | 3.378 | 4.917 | 4.022 | 4.858 | 4.078 | 5.523 | 4.003 | 4.172 | 2.605 | 3.079 | 3.055 | 2.834 | 2.914 | 2.790 |
| MUSC_DCIS | COL1A1 | 21.2 | FDFSFLPQPP | 10 | Unmodified | -0.92 | 1194.583 | 1194.584 | 3.359 | 1.948 | 2.250 | 2.436 | 2.306 | 2.979 | 3.041 | 3.559 | 4.170 | 4.525 | 4.927 | 4.885 | 3.359 | 2.194 | 2.426 | 2.625 | 3.550 | 3.960 | 3.632 | 3.674 | 3.627 | 4.263 | 3.757 | 4.216 | 4.336 | 4.134 | 4.179 | 4.386 | 4.783 | 4.383 | 5.058 | 2.088 | 2.440 | 2.201 | 2.297 | 2.507 | 2.384 |
| MUSC_DCIS | COL1A2 | 20.1 | GARGEPGNIGFP | 12 | GARGEP(1.0000)GNIGFP(1.0000) | -1.66 | 1203.575 | 1203.577 | 5.130 | 2.878 | 3.543 | 4.254 | 3.812 | 5.038 | 5.111 | 5.759 | 6.973 | 6.227 | 6.159 | 7.054 | 5.130 | 3.410 | 3.861 | 4.376 | 5.496 | 6.237 | 5.482 | 5.289 | 5.644 | 6.361 | 6.096 | 5.603 | 6.138 | 5.846 | 6.469 | 6.147 | 7.022 | 6.924 | 7.258 | 3.088 | 3.976 | 3.848 | 3.574 | 4.349 | 4.156 |
| 2022 DCIS Lumpectomies | COL1A2 | 98.0 | GARGEPGNIGFP | 12 | GARGEP(1)GNIGFP(1) | -1.66 | 1203.575 | | | | | | | | | | | | | | | | | | | | | | | | | | | | | | | | | | | | | | |
| MUSC_DCIS | COL1A2 | 21.7 | GPAGEVGKPGERG | 13 | GP(0.0130)AGEVGKP(0.9870)GERG | -1.39 | 1226.612 | 1226.614 | 4.084 | 2.061 | 2.460 | 2.721 | 2.754 | 3.232 | 4.079 | 4.519 | 5.703 | 4.468 | 5.068 | 5.578 | 4.084 | 2.455 | 2.633 | 2.718 | 4.037 | 4.964 | 3.438 | 3.423 | 3.311 | 4.973 | 4.159 | 4.176 | 4.970 | 4.214 | 4.999 | 4.185 | 5.576 | 4.453 | 4.593 | 2.138 | 2.658 | 2.553 | 2.226 | 2.621 | 2.801 |
| 2022 DCIS Lumpectomies | COL1A2 | 171.9 | GPAGEVGKPGERG | 13 | GPAGEVGKP(1)GERG | -1.39 | 1226.612 | | | | | | | | | | | | | | | | | | | | | | | | | | | | | | | | | | | | | | |
| MUSC_DCIS | COL1A1 | 20.2 | GPAGARGNDGATGAA | 15 | Unmodified | -0.80 | 1242.582 | 1242.583 | 7.070 | 3.676 | 5.372 | 5.685 | 5.577 | 6.774 | 7.393 | 7.997 | 8.479 | 8.767 | 8.915 | 8.953 | 7.070 | 4.550 | 5.650 | 5.823 | 7.381 | 7.759 | 8.011 | 8.023 | 8.029 | 8.051 | 8.134 | 8.372 | 8.459 | 8.268 | 8.725 | 8.702 | 8.865 | 8.711 | 9.189 | 4.239 | 5.045 | 5.421 | 5.823 | 5.589 | 5.200 |
| 2022 DCIS Lumpectomies | COL1A1 | 131.1 | GPAGARGNDGATGAA | 15 | Unmodified | -0.80 | 1242.582 | | | | | | | | | | | | | | | | | | | | | | | | | | | | | | | | | | | | | | |
| MUSC_DCIS | COL3A1 | 24.4 | GPRGQPGVMGFP | 12 | GP(0.0367)RGQP(0.9878)GVM(0.9878)GFP( | 1.28 | 1247.584 | 1247.582 | 5.266 | 3.913 | 4.215 | 4.519 | 4.706 | 5.219 | 5.588 | 5.762 | 6.151 | 6.674 | 6.487 | 7.125 | 5.266 | 4.556 | 4.605 | 4.943 | 6.272 | 5.810 | 6.100 | 5.757 | 6.013 | 6.191 | 6.033 | 6.488 | 6.027 | 6.233 | 6.133 | 6.822 | 6.456 | 7.791 | 7.762 | 3.649 | 4.208 | 4.284 | 4.077 | 4.414 | 4.799 |
| 2022 DCIS Lumpectomies | COL3A1 | 81.6 | GPRGQPGVMGFP | 12 | GP(0.006)RGQP(0.994)GVM(1)GFP(1) | 1.28 | 1247.584 | | | | | | | | | | | | | | | | | | | | | | | | | | | | | | | | | | | | | | |
| MUSC_DCIS | COL1A1 | 23.0 | GPAGEKGSPGADGPA | 15 | Unmodified | -3.79 | 1267.591 | 1267.596 | 3.912 | 2.582 | 3.051 | 3.394 | 3.304 | 3.900 | 4.624 | 5.029 | 5.220 | 5.440 | 5.166 | 5.481 | 3.912 | 3.297 | 3.205 | 3.444 | 4.175 | 4.322 | 4.709 | 4.484 | 4.755 | 4.425 | 5.024 | 5.039 | 4.799 | 4.639 | 5.709 | 5.272 | 5.250 | 5.263 | 5.607 | 2.652 | 2.848 | 3.221 | 3.209 | 3.208 | 3.099 |
| 2022 DCIS Lumpectomies | COL1A1 | 192.5 | GPAGEKGSPGADGPA | 15 | Unmodified | -3.79 | 1267.591 | | | | | | | | | | | | | | | | | | | | | | | | | | | | | | | | | | | | | | |
| MUSC_DCIS | COL1A1 | 20.4 | GQRGERGFPGLP | 12 | Unmodified | 3.15 | 1270.665 | 1270.661 | 2.559 | 1.693 | 1.776 | 2.975 | 1.908 | 2.169 | 3.333 | 3.176 | 3.612 | 3.613 | 3.623 | 3.315 | 2.559 | 2.147 | 1.995 | 2.213 | 2.516 | 2.717 | 3.070 | 2.446 | 3.188 | 3.162 | 3.915 | 3.162 | 2.987 | 2.713 | 4.532 | 3.228 | 3.657 | 3.667 | 4.466 | 1.834 | 1.893 | 2.125 | 1.926 | 1.949 | 1.956 |
| MUSC_DCIS | COL1A1 | 20.1 | GAPGNDGAKGDAGAP | 15 | GAP(0.9856)GNDGAKGDAGAP(0.0144) | -1.02 | 1271.550 | 1271.551 | 3.981 | 2.965 | 3.317 | 3.625 | 3.757 | 4.074 | 4.585 | 5.124 | 5.141 | 5.221 | 5.259 | 5.734 | 3.981 | 3.228 | 3.468 | 3.868 | 4.444 | 4.723 | 4.491 | 4.782 | 4.729 | 4.979 | 4.851 | 4.820 | 4.961 | 5.103 | 5.335 | 5.324 | 5.319 | 5.804 | 6.103 | 2.826 | 3.097 | 3.113 | 3.199 | 3.326 | 3.612 |
| MUSC_DCIS | COL5A1 | 23.5 | GLRGFPGDRGLP | 12 | GLRGFP(1.0000)GDRGLP(1.0000) | 0.39 | 1273.665 | 1273.664 | 3.198 | 1.707 | 2.112 | 2.350 | 2.355 | 3.030 | 3.575 | 4.065 | 4.154 | 4.309 | 4.773 | 5.131 | 3.198 | 2.382 | 2.634 | 2.695 | 3.710 | 3.863 | 3.771 | 3.536 | 3.857 | 3.839 | 3.813 | 4.110 | 3.789 | 4.034 | 4.389 | 4.534 | 4.244 | 4.699 | 5.522 | 1.954 | 2.172 | 2.191 | 2.251 | 2.345 | 2.253 |
| 2022 DCIS Lumpectomies | COL5A1 | 157.8 | GLRGFPGDRGLP | 12 | GLRGFP(1)GDRGLP(1) | 0.39 | 1273.665 | | | | | | | | | | | | | | | | | | | | | | | | | | | | | | | | | | | | | | |
| MUSC_DCIS | COL1A2 | 20.9 | GPAGARGSDGSVGPV | 15 | Unmodified | -2.49 | 1283.634 | 1283.637 | 4.944 | 2.532 | 3.066 | 3.544 | 3.442 | 4.232 | 4.977 | 5.623 | 6.747 | 6.397 | 6.826 | 6.801 | 4.944 | 3.125 | 3.837 | 3.541 | 5.228 | 5.995 | 5.563 | 5.758 | 5.582 | 6.105 | 5.860 | 6.095 | 6.524 | 5.934 | 6.009 | 6.079 | 7.145 | 6.494 | 6.838 | 2.865 | 3.618 | 3.422 | 3.931 | 3.304 | 3.426 |
| 2022 DCIS Lumpectomies | COL1A2 | 130.0 | GPAGARGSDGSVGPV | 15 | Unmodified | -2.49 | 1283.634 | | | | | | | | | | | | | | | | | | | | | | | | | | | | | | | | | | | | | | |
| MUSC_DCIS | COL1A1 | 20.1 | GQRGERGFPGLP | 12 | GQRGERGFP(0.0120)GLP(0.9880) | 2.25 | 1286.660 | 1286.657 | 5.917 | 3.079 | 3.240 | 5.167 | 4.024 | 4.713 | 5.755 | 5.756 | 7.129 | 7.144 | 7.188 | 6.872 | 5.917 | 4.244 | 4.837 | 4.777 | 5.432 | 6.210 | 6.250 | 5.347 | 6.594 | 6.330 | 7.000 | 6.351 | 6.269 | 5.511 | 7.242 | 6.760 | 7.195 | 7.215 | 8.046 | 3.616 | 3.957 | 4.614 | 4.018 | 4.980 | 4.517 |
| 2022 DCIS Lumpectomies | COL2A1;COL1A1 | 121.6 | GQRGERGFPGLP | 12 | GQRGERGFPGLP(1) | 2.25 | 1286.660 | | | | | | | | | | | | | | | | | | | | | | | | | | | | | | | | | | | | | | |
| MUSC_DCIS | COL6A1 | 21.7 | GPAGNGTEGFPGFP | 14 | GP(0.0068)AGNGTEGFP(0.0071)GFP(0.9861 | 2.57 | 1321.569 | 1321.566 | 4.418 | 3.055 | 3.928 | 4.110 | 4.286 | 4.485 | 5.791 | 6.099 | 5.458 | 5.666 | 5.585 | 5.808 | 4.418 | 3.694 | 4.123 | 4.314 | 4.766 | 5.054 | 5.533 | 5.674 | 5.463 | 5.333 | 5.377 | 5.412 | 5.391 | 5.681 | 6.287 | 5.839 | 5.722 | 6.386 | 6.223 | 3.184 | 3.474 | 3.573 | 4.215 | 3.812 | 3.903 |
| MUSC_DCIS | COL3A1 | 26.6 | GPLGIAGITGARGLA | 15 | Unmodified | -0.53 | 1323.774 | 1323.775 | 3.195 | 3.897 | 3.554 | 3.450 | 3.477 | 4.030 | 5.516 | 6.040 | 3.818 | 5.540 | 4.771 | 5.088 | 3.195 | 3.473 | 3.533 | 3.623 | 4.583 | 4.140 | 5.637 | 5.689 | 5.449 | 4.634 | 4.716 | 5.690 | 4.832 | 5.543 | 4.955 | 5.682 | 4.666 | 6.431 | 6.053 | 3.182 | 3.308 | 3.377 | 4.882 | 3.256 | 3.151 |
| 2022 DCIS Lumpectomies | COL3A1 | 142.8 | GPLGIAGITGARGLA | 15 | Unmodified | -0.53 | 1323.774 | | | | | | | | | | | | | | | | | | | | | | | | | | | | | | | | | | | | | | |
| MUSC_DCIS | COL1A1 | 25.5 | GATGFPGAAGRVGPP | 15 | GATGFP(0.7145)GAAGRVGP(0.6428)P(0.642 | 2.23 | 1343.670 | 1343.667 | 6.553 | 4.428 | 5.190 | 6.088 | 5.428 | 6.251 | 7.122 | 7.383 | 7.853 | 7.950 | 7.806 | 8.174 | 6.553 | 5.367 | 5.578 | 5.928 | 6.823 | 7.009 | 7.317 | 6.775 | 7.384 | 7.873 | 7.657 | 7.216 | 7.834 | 7.051 | 8.535 | 7.977 | 8.407 | 8.617 | 9.232 | 4.699 | 5.335 | 5.640 | 6.021 | 5.870 | 5.891 |
| MUSC_DCIS | COL1A1 | 25.5 | GAPGIAGAPGFPGAR | 15 | GAP(1.0000)GIAGAP(1.0000)GFP(1.0000)GA | 2.23 | 1343.670 | | | | | | | | | | | | | | | | | | | | | | | | | | | | | | | | | | | | | | |
| 2022 DCIS Lumpectomies | COL1A1 | 155.8 | GAPGIAGAPGFPGAR | 15 | GAP(1)GIAGAP(1)GFP(1)GAR | 2.23 | 1343.670 | | | | | | | | | | | | | | | | | | | | | | | | | | | | | | | | | | | | | | |
| MUSC_DCIS | COL1A1 | 20.2 | GPAGPAGERGEQGPA | 15 | Unmodified | 1.18 | 1350.640 | 1350.638 | 5.997 | 3.850 | 5.222 | 5.480 | 5.059 | 6.050 | 6.195 | 6.690 | 7.084 | 7.449 | 7.018 | 8.079 | 5.997 | 4.991 | 5.194 | 5.580 | 6.201 | 6.501 | 7.037 | 6.400 | 7.181 | 6.803 | 7.430 | 6.526 | 7.089 | 6.873 | 7.241 | 7.365 | 7.758 | 8.873 | 8.893 | 4.035 | 4.621 | 5.179 | 4.543 | 5.442 | 4.790 |
| MUSC_DCIS | COL1A1 | 21.6 | GPAGKSGDRGETGPA | 15 | Unmodified | 0.81 | 1356.650 | 1356.649 | 3.765 | 2.701 | 3.183 | 3.228 | 3.184 | 3.693 | 3.859 | 4.252 | 5.210 | 4.740 | 4.809 | 5.262 | 3.765 | 3.271 | 3.293 | 3.313 | 4.202 | 4.612 | 3.841 | 3.777 | 3.942 | 4.707 | 4.232 | 4.328 | 4.326 | 4.318 | 4.607 | 4.584 | 4.987 | 5.178 | 5.175 | 2.833 | 3.224 | 3.044 | 2.786 | 3.221 | 3.118 |
| MUSC_DCIS | COL1A1 | 27.5 | GATGFPGAAGRVGPP | 15 | GATGFP(1.0000)GAAGRVGP(1.0000)P(1.000 | -3.68 | 1359.665 | 1359.670 | 5.555 | 3.425 | 4.283 | 4.763 | 4.789 | 6.090 | 6.016 | 6.653 | 7.487 | 7.229 | 7.252 | 8.700 | 5.555 | 4.422 | 5.329 | 5.187 | 6.337 | 7.021 | 6.694 | 6.899 | 7.171 | 6.868 | 6.630 | 7.259 | 7.210 | 7.562 | 7.064 | 7.471 | 7.949 | 8.622 | 8.811 | 3.774 | 4.437 | 4.329 | 4.615 | 4.501 | 4.664 |
| MUSC_DCIS | COL18A1 | 20.0 | GPPGQFPFDFLQ | 12 | GP(0.0483)P(0.9398)GQFP(0.0119)FDFLQ | -3.51 | 1365.647 | 1365.652 | 5.047 | 3.156 | 4.060 | 4.709 | 4.242 | 4.783 | 5.880 | 6.003 | 6.375 | 6.532 | 6.237 | 6.333 | 5.047 | 3.877 | 3.876 | 4.798 | 5.307 | 5.744 | 5.642 | 5.467 | 6.034 | 6.229 | 6.425 | 5.341 | 5.889 | 5.669 | 6.823 | 6.439 | 6.523 | 7.555 | 7.799 | 3.190 | 3.830 | 4.267 | 4.290 | 4.712 | 4.941 |
| 2022 DCIS Lumpectomies | COL18A1 | 96.7 | GPPGQFPFDFLQ | 12 | GP(0.2)P(0.8)GQFPFDFLQ | -3.51 | 1365.647 | | | | | | | | | | | | | | | | | | | | | | | | | | | | | | | | | | | | | | |
| MUSC_DCIS | COL1A2 | 25.1 | GPVGRTGEVGAVGPP | 15 | GP(0.0065)VGRTGEVGAVGP(0.0106)P(0.982 | -1.46 | 1365.712 | 1365.714 | 4.500 | 1.889 | 2.104 | 2.816 | 2.836 | 3.959 | 4.014 | 4.345 | 5.719 | 6.098 | 6.449 | 6.251 | 4.500 | 2.698 | 3.123 | 3.031 | 4.722 | 5.057 | 5.129 | 5.431 | 5.174 | 5.546 | 5.329 | 5.713 | 5.706 | 5.726 | 6.171 | 5.917 | 6.119 | 5.463 | 6.206 | 1.952 | 2.249 | 2.296 | 3.852 | 2.976 | 2.802 |
| 2022 DCIS Lumpectomies | COL1A2 | 132.9 | GPVGRTGEVGAVGPP | 15 | GPVGRTGEVGAVGP(0.5)P(0.5) | -1.46 | 1365.712 | | | | | | | | | | | | | | | | | | | | | | | | | | | | | | | | | | | | | | |
| MUSC_DCIS | COL6A2 | 30.6 | FVFLTDGVTGNDS | 13 | Unmodified | 4.15 | 1372.627 | 1372.621 | 4.383 | 2.725 | 3.972 | 4.029 | 3.748 | 4.293 | 4.980 | 5.495 | 5.202 | 5.488 | 5.292 | 5.960 | 4.383 | 3.404 | 3.631 | 4.183 | 4.571 | 4.777 | 5.131 | 4.835 | 5.348 | 5.238 | 5.542 | 4.904 | 5.090 | 5.313 | 5.857 | 5.630 | 5.642 | 7.314 | 7.231 | 2.895 | 3.191 | 3.558 | 3.233 | 3.904 | 3.554 |
| MUSC_DCIS | COL6A1 | 20.9 | LASAVDAMDFIND | 13 | Unmodified | 1.66 | 1381.630 | 1381.628 | 3.998 | 2.624 | 3.570 | 3.853 | 3.767 | 4.195 | 4.659 | 5.564 | 5.305 | 5.704 | 5.260 | 6.399 | 3.998 | 3.265 | 3.765 | 4.133 | 4.263 | 5.204 | 4.592 | 5.317 | 5.421 | 4.914 | 5.810 | 5.160 | 4.913 | 5.703 | 5.456 | 6.281 | 5.580 | 7.183 | 7.391 | 2.964 | 3.079 | 3.222 | 3.456 | 3.522 | 4.431 |
| MUSC_DCIS | COL1A1 | 25.8 | GPPGEPGEPGASGPM | 15 | GP(0.0111)P(0.0111)GEP(0.8185)GEP(0.818 | 1.08 | 1384.569 | 1384.567 | 3.730 | 2.406 | 3.003 | 3.073 | 3.105 | 3.450 | 4.042 | 4.302 | 4.748 | 4.987 | 4.662 | 4.883 | 3.730 | 2.711 | 2.976 | 3.418 | 3.781 | 4.207 | 4.301 | 4.295 | 4.594 | 4.226 | 4.430 | 4.342 | 4.711 | 4.418 | 4.970 | 4.839 | 4.779 | 5.328 | 5.351 | 2.611 | 2.550 | 2.906 | 2.752 | 2.972 | 2.946 |
| 2022 DCIS Lumpectomies | COL1A1 | 109.0 | GPPGEPGEPGASGPM | 15 | GP(0.003)P(0.005)GEP(0.992)GEP(1)GASG | 1.08 | 1384.569 | | | | | | | | | | | | | | | | | | | | | | | | | | | | | | | | | | | | | | |
| MUSC_DCIS | COL6A1 | 21.1 | LASAVDAMDFIND | 13 | LASAVDAM(1.0000)DFIND | 2.29 | 1397.625 | 1397.622 | 3.276 | 2.442 | 3.037 | 3.064 | 3.156 | 3.462 | 3.727 | 4.380 | 4.442 | 4.650 | 4.537 | 5.762 | 3.276 | 2.724 | 3.101 | 3.451 | 3.715 | 4.260 | 3.957 | 4.874 | 4.880 | 4.154 | 4.473 | 4.568 | 4.297 | 5.593 | 4.586 | 5.651 | 4.924 | 6.279 | 6.959 | 2.574 | 2.650 | 2.665 | 2.743 | 2.854 | 3.254 |
| MUSC_DCIS | COL1A1 | 23.5 | GSPGEAGRPGEAGLP | 15 | GSP(1.0000)GEAGRP(1.0000)GEAGLP(1.000 | -3.72 | 1399.645 | 1399.650 | 6.166 | 3.556 | 4.498 | 4.769 | 4.705 | 5.903 | 5.984 | 7.096 | 7.777 | 7.762 | 7.949 | 8.128 | 6.166 | 4.184 | 5.048 | 5.068 | 6.476 | 7.039 | 6.987 | 6.899 | 7.034 | 6.978 | 7.007 | 7.313 | 7.671 | 7.312 | 7.347 | 7.568 | 8.186 | 7.592 | 8.131 | 3.830 | 4.460 | 4.590 | 4.978 | 4.610 | 4.717 |
| 2022 DCIS Lumpectomies | COL1A1 | 124.2 | GSPGEAGRPGEAGLP | 15 | GSP(1)GEAGRP(1)GEAGLP(1) | -3.72 | 1399.645 | | | | | | | | | | | | | | | | | | | | | | | | | | | | | | | | | | | | | | |
| MUSC_DCIS | COL1A1 | 40.7 | GPQGIAGQRGVVGLP | 15 | GP(0.0166)QGIAGQRGVVGLP(0.9834) | -3.59 | 1421.786 | 1421.791 | 4.374 | 3.019 | 3.292 | 3.888 | 3.616 | 4.443 | 5.232 | 6.022 | 5.627 | 6.398 | 5.974 | 6.153 | 4.374 | 3.597 | 4.175 | 4.147 | 4.967 | 5.254 | 6.326 | 5.872 | 6.123 | 5.541 | 5.883 | 6.007 | 5.847 | 5.756 | 5.984 | 6.365 | 5.985 | 7.470 | 7.442 | 3.046 | 3.379 | 3.640 | 4.851 | 3.851 | 3.548 |
| 2022 DCIS Lumpectomies | COL1A1 | 153.1 | GPQGIAGQRGVVGLP | 15 | GPQGIAGQRGVVGLP(1) | -3.59 | 1421.786 | | | | | | | | | | | | | | | | | | | | | | | | | | | | | | | | | | | | | | |
| MUSC_DCIS | COL1A2 | 22.0 | GPAGKDGRTGHPGTV | 15 | GP(0.0129)AGKDGRTGHP(0.9871)GTV | 0.21 | 1422.708 | 1422.708 | 4.291 | 2.830 | 3.860 | 3.795 | 4.780 | 5.813 | 5.946 | 6.806 | 6.685 | 6.355 | 6.748 | 8.541 | 4.291 | 4.227 | 4.917 | 4.705 | 5.653 | 6.504 | 6.096 | 6.661 | 6.636 | 6.089 | 5.410 | 6.971 | 6.491 | 7.372 | 6.763 | 6.956 | 7.401 | 8.475 | 8.537 | 2.899 | 4.143 | 3.071 | 3.731 | 3.581 | 3.912 |
| MUSC_DCIS | COL1A2 | 23.4 | GSRGLPGADGRAGVM | 15 | GSRGLP(1.0000)GADGRAGVM(1.0000) | 0.70 | 1432.696 | 1432.695 | 3.704 | 1.964 | 2.427 | 2.794 | 2.294 | 2.929 | 3.326 | 3.634 | 5.641 | 5.451 | 6.162 | 5.622 | 3.704 | 2.938 | 2.473 | 2.656 | 4.000 | 4.573 | 3.832 | 4.273 | 4.444 | 4.996 | 4.310 | 4.823 | 5.129 | 4.643 | 4.613 | 5.130 | 5.975 | 4.904 | 5.694 | 2.184 | 2.704 | 2.246 | 2.276 | 2.462 | 2.565 |
| MUSC_DCIS | COL1A2 | 20.1 | GMTGFPGAAGRTGPP | 15 | GM(1.0000)TGFP(1.0000)GAAGRTGP(1.0000 | 2.50 | 1437.643 | 1437.639 | 5.551 | 2.959 | 4.049 | 4.384 | 4.029 | 5.297 | 5.670 | 6.324 | 7.097 | 7.014 | 7.341 | 7.681 | 5.551 | 3.628 | 3.942 | 4.530 | 5.914 | 6.432 | 6.135 | 6.122 | 6.513 | 6.975 | 6.496 | 6.396 | 6.944 | 6.603 | 7.489 | 7.075 | 7.543 | 7.533 | 8.123 | 3.282 | 4.010 | 4.152 | 4.013 | 4.471 | 4.523 |
| MUSC_DCIS | COL5A2 | 22.0 | GPPGHLTAALGDIMG | 15 | GP(0.6430)P(0.6430)GHLTAALGDIM(0.7140) | -2.57 | 1438.699 | 1438.703 | 6.248 | 3.095 | 4.281 | 4.959 | 4.671 | 5.967 | 6.056 | 6.653 | 7.728 | 7.800 | 7.856 | 8.444 | 6.248 | 4.554 | 4.677 | 5.191 | 6.531 | 7.019 | 6.848 | 6.681 | 7.247 | 7.351 | 7.302 | 6.996 | 7.311 | 7.277 | 7.786 | 7.687 | 8.113 | 8.509 | 9.089 | 3.274 | 5.001 | 4.578 | 4.443 | 5.207 | 5.172 |
| 2022 DCIS Lumpectomies | COL5A2 | 111.1 | GPPGHLTAALGDIMG | 15 | GP(0.5)P(0.5)GHLTAALGDIM(1)G | -2.57 | 1438.700 | | | | | | | | | | | | | | | | | | | | | | | | | | | | | | | | | | | | | | |
| MUSC_DCIS | COL1A1 | 23.3 | GPPGARGQAGVMGFP | 15 | GP(0.0755)P(0.9747)GARGQAGVM(0.9749)G | 3.66 | 1446.679 | 1446.674 | 6.041 | 4.067 | 4.840 | 5.545 | 5.262 | 5.987 | 6.562 | 6.763 | 7.309 | 7.264 | 7.127 | 7.610 | 6.041 | 5.444 | 5.324 | 5.675 | 6.428 | 6.687 | 6.707 | 6.518 | 6.959 | 7.161 | 7.021 | 6.714 | 6.751 | 6.860 | 7.455 | 7.445 | 7.409 | 8.651 | 8.601 | 4.185 | 5.156 | 5.336 | 4.812 | 5.605 | 5.976 |
| MUSC_DCIS | COL1A1 | 25.2 | GLQGMPGERGAAGLP | 15 | GLQGM(1.0000)P(1.0000)GERGAAGLP(1.00 | 3.70 | 1458.700 | 1458.695 | 5.904 | 2.605 | 3.713 | 4.440 | 4.307 | 5.518 | 5.728 | 6.477 | 7.614 | 7.416 | 7.823 | 7.924 | 5.904 | 3.488 | 4.869 | 4.709 | 6.291 | 6.950 | 6.377 | 6.660 | 6.722 | 7.448 | 6.981 | 6.838 | 7.144 | 7.058 | 7.622 | 7.421 | 7.901 | 7.265 | 7.956 | 3.255 | 3.984 | 3.989 | 4.800 | 4.718 | 4.501 |
| 2022 DCIS Lumpectomies | COL1A1 | 101.6 | GLQGMPGERGAAGLP | 15 | GLQGM(1)P(1)GERGAAGLP(1) | 3.84 | 1458.701 | | | | | | | | | | | | | | | | | | | | | | | | | | | | | | | | | | | | | | |
| MUSC_DCIS | COL1A1 | 20.5 | GPPGERGGPGSRGFP | 15 | GP(0.0421)P(0.9860)GERGGP(0.9860)GSRG | -2.99 | 1472.688 | 1472.692 | 3.733 | 2.343 | 2.592 | 2.715 | 2.618 | 3.069 | 3.824 | 4.156 | 4.893 | 4.412 | 5.152 | 4.712 | 3.733 | 2.571 | 2.950 | 2.824 | 3.931 | 4.357 | 4.097 | 3.780 | 3.771 | 4.801 | 4.038 | 4.060 | 4.382 | 4.088 | 4.180 | 4.290 | 5.013 | 4.830 | 5.094 | 2.334 | 2.855 | 2.659 | 3.023 | 2.736 | 2.735 |
| MUSC_DCIS | COL3A1 | 24.2 | GPRGERGEAGIPGVP | 15 | GP(0.0314)RGERGEAGIP(0.9843)GVP(0.984 | -3.31 | 1480.750 | 1480.755 | 5.041 | 1.954 | 2.434 | 2.888 | 3.264 | 4.467 | 3.963 | 4.547 | 6.465 | 6.278 | 6.549 | 7.161 | 5.041 | 2.328 | 3.031 | 3.521 | 6.174 | 6.081 | 4.981 | 5.120 | 5.302 | 6.290 | 5.464 | 6.193 | 6.006 | 5.939 | 5.830 | 6.081 | 6.748 | 6.303 | 7.022 | 2.207 | 3.212 | 2.321 | 3.176 | 3.393 | 3.224 |
| 2022 DCIS Lumpectomies | COL3A1 | 99.8 | GPRGERGEAGIPGVP | 15 | GPRGERGEAGIP(1)GVP(1) | -3.31 | 1480.750 | | | | | | | | | | | | | | | | | | | | | | | | | | | | | | | | | | | | | | |
| MUSC_DCIS | COL1A2 | 21.2 | GPAGARGSDGSVGPVGPA | 18 | Unmodified | -3.91 | 1508.745 | 1508.751 | 5.944 | 3.866 | 4.810 | 5.374 | 5.141 | 6.047 | 6.510 | 6.763 | 7.108 | 7.648 | 7.449 | 8.129 | 5.944 | 5.410 | 5.308 | 5.498 | 6.248 | 6.526 | 6.883 | 6.618 | 7.114 | 6.813 | 7.298 | 6.934 | 7.107 | 6.971 | 7.498 | 7.464 | 7.633 | 8.455 | 8.623 | 4.162 | 4.703 | 5.268 | 5.128 | 5.383 | 5.412 |
| 2022 DCIS Lumpectomies | COL1A2 | 132.1 | GPAGARGSDGSVGPVGPA | 18 | Unmodified | -3.91 | 1508.745 | | | | | | | | | | | | | | | | | | | | | | | | | | | | | | | | | | | | | | |
| 2022 DCIS Lumpectomies | COL1A1 | 109.6 | PPQEKAHDGGRYY | 13 | Unmodified | 2.77 | 1517.713 | 1517.709 | 3.924 | 2.694 | 3.736 | 4.065 | 3.766 | 4.254 | 5.628 | 5.906 | 5.042 | 5.524 | 4.866 | 5.729 | 3.924 | 3.888 | 3.762 | 4.305 | 4.396 | 4.832 | 5.161 | 4.742 | 5.381 | 5.031 | 5.484 | 4.670 | 4.821 | 5.182 | 6.143 | 5.627 | 5.287 | 7.313 | 7.300 | 2.782 | 3.189 | 3.692 | 3.393 | 3.865 | 3.773 |
| MUSC_DCIS | COL1A2 | 20.4 | GPRGDQGPVGRTGEVG | 16 | Unmodified | -0.71 | 1538.767 | 1538.768 | 2.785 | 1.864 | 2.011 | 2.078 | 2.108 | 2.421 | 2.906 | 2.860 | 4.112 | 3.495 | 3.707 | 4.348 | 2.785 | 2.092 | 2.140 | 2.305 | 3.058 | 3.630 | 2.662 | 2.690 | 2.941 | 3.844 | 3.280 | 3.083 | 3.375 | 3.333 | 3.622 | 3.413 | 4.354 | 4.340 | 4.484 | 1.924 | 2.126 | 2.070 | 1.990 | 2.152 | 2.457 |
| MUSC_DCIS | COL1A1 | 27.8 | GPPGAVGPAGKDGEAGAQ | 18 | GP(0.0118)P(0.9813)GAVGP(0.0069)AGKDG | -0.26 | 1551.740 | | | | | | | | | | | | | | | | | | | | | | | | | | | | | | | | | | | | | | |
| 2022 DCIS Lumpectomies | COL1A1 | 78.1 | GPPGAVGPAGKDGEAGAQ | 18 | GP(0.09)P(0.91)GAVGPAGKDGEAGAQ | -0.26 | 1551.740 | | | | | | | | | | | | | | | | | | | | | | | | | | | | | | | | | | | | | | |
| MUSC_DCIS | COL1A2 | 45.9 | GPVGAAGATGARGLVGEP | 18 | GP(0.0143)VGAAGATGARGLVGEP(0.9857) | -2.19 | 1552.808 | 1552.811 | 3.847 | 3.071 | 3.612 | 4.194 | 3.809 | 4.445 | 5.714 | 6.165 | 5.221 | 5.812 | 5.282 | 6.068 | 3.847 | 3.981 | 4.266 | 4.191 | 4.413 | 4.903 | 5.466 | 5.469 | 5.683 | 5.406 | 5.813 | 5.257 | 5.102 | 5.418 | 6.250 | 5.918 | 5.464 | 7.358 | 7.218 | 3.366 | 3.509 | 3.774 | 4.536 | 4.056 | 3.768 |
| 2022 DCIS Lumpectomies | COL1A2 | 90.0 | GPVGAAGATGARGLVGEP | 18 | GPVGAAGATGARGLVGEP(1) | -2.19 | 1552.808 | | | | | | | | | | | | | | | | | | | | | | | | | | | | | | | | | | | | | | |
| MUSC_DCIS | COL1A1 | 25.0 | GARGERGFPGERGVQ | 15 | GARGERGFP(1.0000)GERGVQ | -2.08 | 1588.794 | 1588.797 | 3.826 | 1.658 | 1.937 | 2.134 | 2.000 | 2.567 | 2.811 | 3.030 | 5.809 | 4.628 | 5.965 | 5.381 | 3.826 | 1.807 | 2.082 | 2.309 | 3.877 | 4.972 | 3.452 | 3.610 | 4.083 | 5.201 | 3.903 | 4.480 | 4.952 | 4.169 | 4.228 | 4.243 | 6.019 | 5.159 | 5.717 | 1.787 | 2.835 | 1.948 | 1.979 | 1.983 | 2.225 |
| MUSC_DCIS | COL1A2 | 22.6 | GAVGSPGVNGAPGEAGRD | 18 | GAVGSP(1.0000)GVNGAP(1.0000)GEAGRD | -3.38 | 1599.736 | 1599.741 | 3.441 | 2.340 | 2.721 | 2.973 | 2.715 | 3.434 | 3.760 | 4.184 | 4.733 | 4.674 | 4.932 | 5.397 | 3.441 | 2.740 | 2.942 | 3.160 | 3.686 | 4.071 | 3.950 | 3.852 | 4.405 | 4.199 | 4.322 | 4.206 | 4.490 | 4.382 | 5.018 | 4.643 | 5.077 | 5.086 | 5.275 | 2.266 | 2.594 | 2.672 | 2.813 | 2.834 | 2.619 |
| 2022 DCIS Lumpectomies | COL1A2 | 111.1 | GAVGSPGVNGAPGEAGRD | 18 | GAVGSP(1)GVNGAP(1)GEAGRD | -3.38 | 1599.736 | | | | | | | | | | | | | | | | | | | | | | | | | | | | | | | | | | | | | | |
| MUSC_DCIS | COL12A1 | 31.1 | VANLPYKGGNTLTGMA | 16 | VANLP(0.0142)YKGGNTLTGM(0.9858)A | -0.92 | 1622.821 | 1622.822 | 3.826 | 1.658 | 1.897 | 2.229 | 2.177 | 3.370 | 3.339 | 3.581 | 5.442 | 5.638 | 5.584 | 6.472 | 3.826 | 2.012 | 2.280 | 2.505 | 4.055 | 4.754 | 4.346 | 3.975 | 4.291 | 4.917 | 4.988 | 4.532 | 5.088 | 4.758 | 5.544 | 5.075 | 5.985 | 5.683 | 6.085 | 1.801 | 2.536 | 1.985 | 2.038 | 2.451 | 2.938 |
| MUSC_DCIS | COL1A1 | 21.4 | GPAGKEGGKGPRGETGPA | 18 | Unmodified | 1.48 | 1622.824 | | | | | | | | | | | | | | | | | | | | | | | | | | | | | | | | | | | | | | |
| MUSC_DCIS | COL3A1 | 20.7 | GVKGERGSPGGPGAAGFP | 18 | GVKGERGSP(1.0000)GGP(1.0000)GAAGFP( | 0.43 | 1645.793 | 1645.792 | 3.307 | 1.947 | 2.455 | 2.658 | 2.557 | 3.235 | 3.796 | 4.098 | 4.619 | 4.715 | 4.925 | 5.163 | 3.307 | 2.363 | 2.609 | 2.948 | 3.868 | 4.070 | 3.882 | 3.815 | 4.220 | 4.489 | 4.366 | 4.189 | 4.144 | 4.377 | 4.837 | 4.651 | 4.949 | 5.713 | 5.760 | 2.004 | 2.423 | 2.312 | 2.617 | 2.692 | 2.752 |
| MUSC_DCIS | COL3A1 | 24.4 | GPRGAAGEPGRDGVPGGP | 18 | GP(0.0472)RGAAGEP(0.9842)GRDGVP(0.98 | 0.73 | 1651.778 | | | | | | | | | | | | | | | | | | | | | | | | | | | | | | | | | | | | | | |
| MUSC_DCIS | COL1A2 | 20.9 | GPSGEEGKRGPNGEAGSA | 18 | Unmodified | 0.06 | 1656.757 | 1656.757 | 4.180 | 2.384 | 2.863 | 3.019 | 2.901 | 3.360 | 3.893 | 4.284 | 6.101 | 5.419 | 6.010 | 6.012 | 4.180 | 2.641 | 2.820 | 3.084 | 4.365 | 5.235 | 4.104 | 4.222 | 4.469 | 5.366 | 4.712 | 4.837 | 5.278 | 4.855 | 5.313 | 4.966 | 6.364 | 5.484 | 5.901 | 2.553 | 3.248 | 2.660 | 2.702 | 2.786 | 2.628 |
| MUSC_DCIS | COL1A1 | 22.0 | GERGGPGSRGFPGADGVA | 18 | GERGGP(1.0000)GSRGFP(1.0000)GADGVA | 0.66 | 1675.778 | 1675.777 | 4.202 | 2.204 | 2.608 | 2.866 | 2.764 | 3.474 | 3.503 | 3.911 | 6.025 | 5.182 | 5.661 | 5.927 | 4.202 | 2.643 | 2.727 | 3.156 | 4.387 | 5.368 | 3.959 | 4.088 | 4.470 | 5.479 | 4.539 | 4.607 | 5.066 | 4.595 | 4.914 | 4.808 | 6.136 | 5.585 | 6.224 | 2.233 | 3.385 | 2.618 | 2.411 | 2.893 | 3.120 |
| MUSC_DCIS | COL1A1 | 20.4 | GPKGSPGEAGRPGEAGLP | 18 | GP(0.0500)KGSP(0.9833)GEAGRP(0.9833)G | 1.07 | 1681.814 | 1681.812 | 5.385 | 2.942 | 3.572 | 4.154 | 3.858 | 5.239 | 5.344 | 5.836 | 6.887 | 7.355 | 7.419 | 8.011 | 5.385 | 3.818 | 3.681 | 4.289 | 5.669 | 6.074 | 6.295 | 5.947 | 6.536 | 6.363 | 6.845 | 6.460 | 6.879 | 6.495 | 6.979 | 6.971 | 7.507 | 7.636 | 8.161 | 2.897 | 3.863 | 3.768 | 3.810 | 4.268 | 4.340 |
| MUSC_DCIS | COL1A2 | 24.4 | GDQGPVGRTGEVGAVGPP | 18 | GDQGP(0.0344)VGRTGEVGAVGP(0.9828)P( | 1.07 | 1681.814 | | | | | | | | | | | | | | | | | | | | | | | | | | | | | | | | | | | | | | |
| MUSC_DCIS | COL3A1 | 21.9 | GPNGIPGEKGPAGERGAP | 18 | GP(0.0166)NGIP(0.9499)GEKGP(0.0859)AGE | -0.12 | 1692.830 | 1692.830 | 3.913 | 2.172 | 2.831 | 2.927 | 3.098 | 3.996 | 4.095 | 4.592 | 4.978 | 5.515 | 5.505 | 6.244 | 3.913 | 2.872 | 2.899 | 3.292 | 4.727 | 4.590 | 4.836 | 4.194 | 4.585 | 4.953 | 5.100 | 5.170 | 4.765 | 4.959 | 5.381 | 5.588 | 5.448 | 6.441 | 6.595 | 2.164 | 2.631 | 2.622 | 2.625 | 3.095 | 3.218 |
| 2022 DCIS Lumpectomies | COL3A1 | 140.9 | GPNGIPGEKGPAGERGAP | 18 | GPNGIP(1)GEKGP(0.002)AGERGAP(0.998) | -0.12 | 1692.830 | | | | | | | | | | | | | | | | | | | | | | | | | | | | | | | | | | | | | | |
| MUSC_DCIS | COL1A2 | 21.4 | GPAGVRGPNGDAGRPGEP | 18 | GP(0.0138)AGVRGP(0.0154)NGDAGRP(0.98 | -3.66 | 1693.789 | 1693.795 | 4.861 | 4.405 | 4.645 | 4.983 | 4.952 | 5.131 | 5.587 | 5.333 | 6.395 | 5.550 | 5.865 | 7.011 | 4.861 | 5.708 | 5.491 | 5.195 | 5.007 | 5.751 | 4.765 | 4.314 | 5.347 | 6.179 | 5.800 | 4.829 | 5.484 | 4.716 | 6.008 | 5.196 | 6.640 | 7.191 | 7.374 | 3.727 | 4.778 | 5.292 | 3.811 | 4.952 | 4.505 |
| MUSC_DCIS | COL2A1 | 20.2 | QGPKGEPGPAGPQGAPGPA | 19 | QGP(0.3968)KGEP(0.3968)GP(0.3968)AGP( | -3.52 | 1703.787 | 1703.793 | 3.560 | 2.247 | 2.696 | 2.922 | 2.817 | 3.566 | 4.316 | 4.534 | 4.937 | 5.395 | 5.381 | 5.798 | 3.560 | 2.676 | 2.684 | 3.086 | 3.839 | 4.447 | 4.404 | 4.243 | 4.731 | 4.737 | 4.964 | 4.583 | 4.807 | 4.781 | 5.580 | 5.262 | 5.386 | 6.151 | 6.418 | 2.286 | 2.673 | 2.624 | 2.633 | 2.844 | 3.206 |
| MUSC_DCIS | COL1A1 | 20.2 | GPRGETGPAGRPGEVGPP | 18 | GP(0.0093)RGETGP(0.0095)AGRP(0.7851)G | 5.06 | 1719.841 | 1719.832 | 4.093 | 2.374 | 2.944 | 3.267 | 3.039 | 3.959 | 4.685 | 4.988 | 5.618 | 5.461 | 5.487 | 5.995 | 4.093 | 2.878 | 3.241 | 3.622 | 4.311 | 5.003 | 4.789 | 4.620 | 5.139 | 5.401 | 5.294 | 4.755 | 5.144 | 5.031 | 5.796 | 5.447 | 6.030 | 6.836 | 6.939 | 2.290 | 3.193 | 2.943 | 3.023 | 3.319 | 3.203 |
| MUSC_DCIS | COL1A2 | 22.1 | VRGPNGDAGRPGEPGLM | 17 | VRGP(0.0404)NGDAGRP(0.9866)GEP(0.9864 | -0.12 | 1727.813 | 1727.813 | 3.468 | 2.175 | 2.360 | 2.566 | 2.379 | 2.768 | 3.073 | 3.247 | 4.923 | 4.046 | 4.305 | 4.705 | 3.468 | 2.383 | 2.431 | 2.628 | 3.501 | 4.079 | 3.019 | 3.069 | 3.264 | 4.413 | 3.612 | 3.379 | 3.843 | 3.624 | 4.023 | 3.816 | 4.849 | 4.432 | 4.703 | 2.268 | 2.414 | 2.212 | 2.162 | 2.380 | 2.693 |
| MUSC_DCIS | COL12A1 | 20.6 | LFGEGTTLEERGSPQD | 16 | Unmodified | 4.15 | 1735.813 | 1735.806 | 2.864 | 2.051 | 2.465 | 2.773 | 2.708 | 2.941 | 3.561 | 3.990 | 4.004 | 4.379 | 4.149 | 4.719 | 2.864 | 2.533 | 2.692 | 3.016 | 3.224 | 3.456 | 3.487 | 3.715 | 4.197 | 3.880 | 4.292 | 3.755 | 3.689 | 4.234 | 4.394 | 4.573 | 4.284 | 5.791 | 6.001 | 2.235 | 2.240 | 2.379 | 2.365 | 2.465 | 2.724 |
| 2022 DCIS Lumpectomies | COL12A1 | 274.5 | LFGEGTTLEERGSPQD | 16 | Unmodified | 4.15 | 1735.813 | | | | | | | | | | | | | | | | | | | | | | | | | | | | | | | | | | | | | | |
| MUSC_DCIS | COL3A1 | 20.6 | GPAGQPGDKGEGGAPGLPGIA | 21 | GP(0.0395)AGQP(0.9868)GDKGEGGAP(0.98 | -4.43 | 1851.872 | 1851.880 | 2.795 | 1.704 | 2.091 | 2.136 | 2.083 | 2.324 | 2.806 | 3.021 | 3.972 | 3.915 | 4.349 | 4.052 | 2.795 | 2.315 | 2.170 | 2.388 | 3.095 | 3.556 | 3.131 | 3.247 | 3.483 | 3.872 | 3.486 | 3.474 | 3.674 | 3.493 | 3.714 | 3.651 | 4.343 | 4.706 | 5.186 | 1.803 | 2.046 | 1.966 | 1.813 | 2.046 | 1.985 |
| 2022 DCIS Lumpectomies | COL3A1 | 138.0 | GPAGQPGDKGEGGAPGLPGIA | 21 | GP(0.003)AGQ(1)P(0.997)GDKGEGGAP(1)G | -4.43 | 1851.872 | | | | | | | | | | | | | | | | | | | | | | | | | | | | | | | | | | | | | | |
| MUSC_DCIS | COL1A1 | 20.2 | GAPGTPGPQGIAGQRGVVGLP | 21 | GAP(0.9752)GTP(0.0263)GP(0.0231)QGIAG | -1.09 | 1918.014 | 1918.016 | 2.194 | 1.892 | 1.915 | 2.457 | 1.929 | 2.452 | 3.917 | 4.166 | 3.375 | 4.147 | 3.818 | 4.014 | 2.194 | 2.433 | 2.486 | 2.443 | 2.682 | 3.032 | 4.055 | 3.385 | 4.229 | 3.326 | 4.005 | 3.682 | 3.234 | 3.308 | 4.089 | 4.132 | 3.580 | 6.039 | 5.302 | 1.620 | 1.832 | 2.384 | 2.342 | 2.133 | 2.091 |
| MUSC_DCIS | COL1A1 | 20.6 | GPSGEPGKQGPSGASGERGPP | 21 | GP(0.0034)SGEP(0.0038)GKQGP(0.0039)SG | 2.39 | 1921.900 | 1921.895 | 3.178 | 2.156 | 2.778 | 2.773 | 2.799 | 3.154 | 3.918 | 4.157 | 4.082 | 4.744 | 4.090 | 5.195 | 3.178 | 3.656 | 3.061 | 3.200 | 3.329 | 3.485 | 4.162 | 3.463 | 4.185 | 3.775 | 4.604 | 3.628 | 3.864 | 3.813 | 4.902 | 4.542 | 4.264 | 6.508 | 6.181 | 2.169 | 2.418 | 3.060 | 2.149 | 3.036 | 2.532 |
| MUSC_DCIS | COL3A1 | 20.8 | GAAGTPGLQGMPGERGGLGSP | 21 | GAAGTP(1.0000)GLQGM(1.0000)P(1.0000)G | 0.62 | 1930.892 | 1930.891 | 2.762 | 1.852 | 2.190 | 2.374 | 2.320 | 2.904 | 3.194 | 3.914 | 4.072 | 4.611 | 4.739 | 5.401 | 2.762 | 2.131 | 2.524 | 2.701 | 3.641 | 3.749 | 3.607 | 3.657 | 4.064 | 4.117 | 4.094 | 4.149 | 3.853 | 4.059 | 4.403 | 4.637 | 4.574 | 5.517 | 5.660 | 1.831 | 2.054 | 2.017 | 2.202 | 2.330 | 2.300 |
| 2022 DCIS Lumpectomies | COL3A1 | 152.2 | GAAGTPGLQGMPGERGGLGSP | 21 | GAAGTP(1)GLQGM(1)P(1)GERGGLGSP(1) | 0.62 | 1930.892 | | | | | | | | | | | | | | | | | | | | | | | | | | | | | | | | | | | | | | |
| MUSC_DCIS | COL1A1 | 24.3 | GPSGEPGKQGPSGASGERGPP | 21 | GP(0.0071)SGEP(0.7943)GKQGP(0.0104)SG | 0.77 | 1937.895 | 1937.893 | 3.398 | 2.332 | 3.099 | 3.386 | 2.930 | 3.558 | 4.041 | 4.453 | 4.475 | 4.904 | 4.799 | 5.742 | 3.398 | 3.485 | 3.199 | 3.352 | 3.665 | 3.977 | 4.455 | 3.641 | 4.599 | 4.366 | 4.999 | 4.032 | 4.299 | 4.198 | 4.936 | 4.671 | 5.238 | 6.728 | 6.564 | 2.212 | 2.706 | 3.182 | 2.407 | 3.251 | 2.781 |
| MUSC_DCIS | COL1A1 | 22.8 | GPPGEAGKPGEQGVPGDLGAP | 21 | GP(0.7742)P(0.7742)GEAGKP(0.8171)GEQG | -3.69 | 1950.904 | 1950.911 | 2.896 | 1.967 | 2.215 | 2.517 | 2.282 | 2.901 | 3.192 | 3.678 | 4.252 | 4.340 | 4.513 | 4.978 | 2.896 | 2.543 | 2.555 | 2.628 | 3.402 | 3.706 | 3.810 | 3.398 | 3.992 | 4.062 | 3.969 | 3.830 | 3.919 | 3.818 | 4.507 | 4.250 | 4.695 | 5.384 | 5.382 | 1.710 | 2.065 | 2.359 | 1.994 | 2.418 | 2.514 |
| 2022 DCIS Lumpectomies | COL1A1 | 157.4 | GPPGEAGKPGEQGVPGDLGAP | 21 | GP(0.191)P(0.809)GEAGKP(1)GEQGVP(1)G | -3.69 | 1950.904 | | | | | | | | | | | | | | | | | | | | | | | | | | | | | | | | | | | | | | |
| MUSC_DCIS | COL1A2 | 26.4 | GPRGDQGPVGRTGEVGAVGPP | 21 | GP(0.0040)RGDQGP(0.0036)VGRTGEVGAV | 0.10 | 1975.994 | 1975.994 | 3.418 | 1.186 | 1.627 | 2.284 | 2.029 | 3.301 | 3.090 | 3.628 | 4.922 | 5.172 | 5.327 | 5.634 | 3.418 | 1.552 | 1.877 | 2.707 | 3.739 | 4.204 | 4.341 | 4.076 | 4.676 | 4.639 | 4.818 | 4.578 | 4.734 | 4.335 | 4.926 | 4.890 | 5.475 | 6.253 | 6.250 | 1.289 | 1.975 | 1.789 | 2.172 | 2.357 | 2.633 |
| 2022 DCIS Lumpectomies | COL1A2 | 120.0 | GPRGDQGPVGRTGEVGAVGPP | 21 | GPRGDQGPVGRTGEVGAVGP(0.5)P(0.5) | 0.10 | 1975.994 | | | | | | | | | | | | | | | | | | | | | | | | | | | | | | | | | | | | | | |
| MUSC_DCIS | COL3A1 | 25.7 | GPAGKDGESGRPGRPGERGLP | 21 | GP(0.0558)AGKDGESGRP(0.9814)GRP(0.98 | 0.14 | 2095.027 | 2095.027 | 3.100 | 1.347 | 1.735 | 1.886 | 2.000 | 2.974 | 3.284 | 3.823 | 4.541 | 5.151 | 5.355 | 6.082 | 3.100 | 2.043 | 1.845 | 2.466 | 4.188 | 4.078 | 4.185 | 3.647 | 4.357 | 4.418 | 4.387 | 4.845 | 4.404 | 4.355 | 4.817 | 4.934 | 5.094 | 6.245 | 6.297 | 1.176 | 1.763 | 1.619 | 1.453 | 1.887 | 1.712 |
| 2022 DCIS Lumpectomies | COL3A1 | 115.2 | GPAGKDGESGRPGRPGERGLP | 21 | GPAGKDGESGRP(1)GRP(1)GERGLP(1) | 0.24 | 2095.028 | | | | | | | | | | | | | | | | | | | | | | | | | | | | | | | | | | | | | | |

### Supplemental Table 2 Continued.

| 4590A | 44010J | 43816D | 44507D | 43662A | 43867A | 43963A | 43634A | 46002A | 46043A | 44015D | 44096A | 43143A | 44243A | 43263J | 43972A | 43018A | 43785D | 43077A | 43945D | 46034A | 43854A | 46003A | 43268A | 43088A | 4494A | 46268L | 43961D | 42312J | 43094A | 43146A | 43163A | 43066A | 43095A | 43074A | 46086A | 46109A | 42313J | 43187A | 43906A | 43327A | 44087A | 42309J | 43060A | 43071A | 43160A | 43148A | 43330A | 44095A | 43275A | 47001D | 42288J | 46001A | 43184A | 43855A | 44050A | 42306J |
|---|---|---|---|---|---|---|---|---|---|---|---|---|---|---|---|---|---|---|---|---|---|---|---|---|---|---|---|---|---|---|---|---|---|---|---|---|---|---|---|---|---|---|---|---|---|---|---|---|---|---|---|---|---|---|---|---|
| Non-progre | Non-progre | Non-progre | Non-progre | Non-progre | Non-progre | Non-progre | Non-progre | Non-progre | Non-progre | Non-progre | Non-progre | Non-progre | Non-progre | Non-progre | Non-progre | Non-progre | Non-progre | Non-progre | Non-progre | Non-progre | Non-progre | Non-progre | Non-progre | Non-progre | Non-progre | Non-progre | Non-progre | Non-progre | Non-progre | Non-progre | Non-progre | Non-progre | Non-progre | Non-progre | Non-progre | Non-progre | Non-progre | Non-progre | Non-progre | Non-progre | Non-progre | Non-progre | Non-progre | Non-progre | Non-progre | Non-progre | Non-progre | Non-progre | Non-progre | Non-progre | Non-progre | Non-progre | Non-progre | Non-progre | Non-progre | Non-progre |
| NP | NP | NP | NP | NP | NP | NP | NP | NP | NP | NP | NP | NP | NP | NP | NP | NP | NP | NP | NP | NP | NP | NP | NP | NP | NP | NP | NP | NP | NP | NP | NP | NP | NP | NP | NP | NP | NP | NP | NP | NP | NP | NP | NP | NP | NP | NP | NP | NP | NP | NP | NP | NP | NP | NP | NP | NP |
| [illegible] | [illegible] | [illegible] | [illegible] | [illegible] | [illegible] | [illegible] | [illegible] | [illegible] | [illegible] | [illegible] | [illegible] | [illegible] | [illegible] | [illegible] | [illegible] | [illegible] | [illegible] | [illegible] | [illegible] | [illegible] | [illegible] | [illegible] | [illegible] | [illegible] | [illegible] | [illegible] | [illegible] | [illegible] | [illegible] | [illegible] | [illegible] | [illegible] | [illegible] | [illegible] | [illegible] | [illegible] | [illegible] | [illegible] | [illegible] | [illegible] | [illegible] | [illegible] | [illegible] | [illegible] | [illegible] | [illegible] | [illegible] | [illegible] | [illegible] | [illegible] | [illegible] | [illegible] | [illegible] | [illegible] | [illegible] | [illegible] |

### Supplemental Table 2 Continued.

| 44059A Non-progre | 44065A Non-progre | 43067A Non-progre | 43078A Non-progre | 43090A Non-progre | 43359A Non-progre | 43987D Non-progre | 43820A Non-progre | 46011A Non-progre | 43726D Non-progre | 44005J Non-progre | 42531C Non-progre | 44094A Non-progre | 43179A Non-progre | 43216A Non-progre | 43098A Non-progre | 43084A Non-progre | 43621A Non-progre | 43178A Non-progre | 43000A Non-progre |
|---|---|---|---|---|---|---|---|---|---|---|---|---|---|---|---|---|---|---|---|
| NP | NP | NP | NP | NP | NP | NP | NP | NP | NP | NP | NP | NP | NP | NP | NP | NP | NP | NP | NP |
| 3.627 | 3.745 | 3.517 | 3.931 | 3.353 | 3.478 | 3.717 | 3.529 | 3.779 | 4.024 | 3.829 | 3.538 | 3.795 | 3.520 | 3.548 | 3.560 | 3.445 | 4.025 | 3.854 | 3.650 |
| 5.476 | 5.690 | 5.294 | 5.356 | 4.745 | 4.590 | 5.462 | 5.140 | 5.499 | 5.490 | 5.421 | 5.508 | 4.942 | 4.936 | 4.712 | 4.722 | 4.993 | 5.406 | 5.617 | 5.508 |
| 4.949 | 4.789 | 4.681 | 4.308 | 4.023 | 4.951 | 4.546 | 4.923 | 4.767 | 4.417 | 4.093 | 4.608 | 4.692 | 4.560 | 4.114 | 4.167 | 4.315 | 4.731 | 4.400 | 4.968 |
| 7.159 | 7.200 | 7.193 | 6.983 | 6.744 | 7.500 | 7.229 | 7.518 | 6.274 | 7.532 | 7.053 | 7.370 | 7.275 | 7.379 | 7.062 | 6.709 | 6.792 | 7.402 | 7.432 | 7.553 |
| 6.883 | 6.965 | 6.539 | 7.212 | 6.374 | 6.507 | 6.727 | 6.618 | 6.268 | 6.833 | 6.942 | 7.117 | 6.677 | 6.937 | 7.332 | 7.154 | 7.187 | 6.888 | 7.112 | 6.833 |
| 5.905 | 6.157 | 4.968 | 5.567 | 4.631 | 5.833 | 5.300 | 5.416 | 4.897 | 6.543 | 5.709 | 5.922 | 6.139 | 5.774 | 5.763 | 5.370 | 5.780 | 4.920 | 6.243 | 6.035 |
| 8.153 | 8.206 | 7.956 | 8.416 | 7.663 | 7.957 | 7.789 | 8.082 | 7.869 | 7.799 | 7.830 | 8.257 | 8.500 | 8.293 | 8.260 | 8.517 | 8.522 | 8.132 | 8.417 | 8.292 |
| 4.948 | 5.117 | 4.672 | 5.484 | 4.340 | 4.292 | 4.987 | 4.496 | 4.501 | 5.179 | 5.235 | 5.323 | 5.255 | 4.765 | 5.301 | 5.292 | 4.981 | 4.927 | 5.533 | 4.847 |
| 5.905 | 5.926 | 5.499 | 5.959 | 5.323 | 5.229 | 5.719 | 5.472 | 5.725 | 6.132 | 5.944 | 5.874 | 5.574 | 5.577 | 5.675 | 5.316 | 5.800 | 5.475 | 6.174 | 5.938 |
| 3.400 | 3.345 | 3.812 | 3.935 | 3.690 | 3.742 | 4.028 | 3.747 | 3.464 | 3.861 | 4.203 | 3.994 | 4.420 | 4.039 | 4.110 | 4.684 | 4.028 | 4.142 | 4.022 | 4.066 |
| 5.798 | 5.777 | 5.576 | 5.532 | 5.562 | 5.970 | 5.506 | 5.893 | 5.870 | 6.089 | 5.540 | 5.491 | 6.269 | 5.771 | 5.158 | 5.214 | 5.931 | 5.677 | 5.506 | 5.956 |
| 7.378 | 7.368 | 7.892 | 7.516 | 7.439 | 7.827 | 7.982 | 7.592 | 7.549 | 7.835 | 8.028 | 8.027 | 7.778 | 7.973 | 7.873 | 8.084 | 7.761 | 7.860 | 7.923 | 8.039 |
| 5.909 | 6.072 | 5.343 | 5.785 | 4.743 | 5.458 | 5.120 | 5.582 | 5.441 | 5.783 | 5.339 | 5.996 | 6.096 | 5.331 | 5.546 | 4.978 | 5.565 | 5.328 | 5.634 | 5.636 |
| 5.575 | 5.405 | 4.991 | 5.231 | 5.084 | 5.711 | 5.011 | 5.640 | 5.120 | 5.436 | 4.963 | 5.404 | 6.365 | 5.309 | 4.950 | 5.137 | 5.201 | 4.182 | 4.845 | 5.428 |
| 4.398 | 4.399 | 4.594 | 4.410 | 4.364 | 4.655 | 4.798 | 4.432 | 4.964 | 4.694 | 4.789 | 4.462 | 4.745 | 4.462 | 4.300 | 4.449 | 4.746 | 4.521 | 4.590 | 4.761 |
| 6.272 | 6.744 | 6.111 | 6.418 | 5.498 | 5.192 | 6.221 | 5.879 | 5.932 | 5.580 | 6.033 | 6.030 | 6.109 | 6.290 | 6.689 | 7.155 | 6.633 | 7.561 | 6.839 | 6.497 |
| 5.841 | 5.839 | 4.736 | 4.643 | 4.326 | 4.137 | 4.906 | 4.991 | 5.044 | 5.014 | 4.606 | 4.813 | 4.916 | 4.520 | 4.338 | 4.061 | 4.532 | 4.864 | 5.038 | 5.252 |
| 5.231 | 5.284 | 5.297 | 5.586 | 4.834 | 5.353 | 5.575 | 5.064 | 5.141 | 5.432 | 5.437 | 5.364 | 6.348 | 5.399 | 5.663 | 6.169 | 5.410 | 5.520 | 5.591 | 5.413 |
| 4.707 | 4.796 | 6.042 | 6.162 | 6.059 | 5.640 | 6.377 | 5.456 | 5.377 | 6.203 | 6.620 | 6.628 | 4.989 | 6.347 | 6.672 | 6.916 | 6.111 | 6.340 | 6.182 | 5.912 |
| 4.178 | 4.680 | 3.682 | 4.803 | 3.698 | 3.313 | 4.057 | 3.640 | 3.656 | 3.519 | 4.023 | 3.892 | 3.979 | 3.916 | 4.658 | 5.231 | 4.702 | 5.061 | 4.924 | 4.275 |
| 5.223 | 5.329 | 4.960 | 5.219 | 4.569 | 4.924 | 5.052 | 4.998 | 5.032 | 5.019 | 5.133 | 5.091 | 6.354 | 5.048 | 5.416 | 5.894 | 5.150 | 5.297 | 5.336 | 5.073 |
| 7.528 | 7.473 | 7.862 | 7.376 | 8.091 | 8.198 | 7.699 | 7.944 | 8.112 | 7.802 | 7.606 | 7.438 | 8.483 | 7.669 | 7.431 | 7.499 | 8.022 | 7.696 | 7.336 | 7.967 |
| 7.430 | 7.384 | 7.661 | 7.363 | 7.253 | 7.784 | 7.680 | 7.640 | 7.608 | 7.617 | 7.648 | 7.817 | 7.281 | 7.722 | 7.481 | 7.602 | 7.642 | 7.673 | 7.682 | 7.892 |
| 5.153 | 5.191 | 4.316 | 4.572 | 4.521 | 4.676 | 4.523 | 4.798 | 4.948 | 4.655 | 4.444 | 4.335 | 5.073 | 4.278 | 4.085 | 4.260 | 4.730 | 4.281 | 4.801 | 4.748 |
| 5.367 | 5.375 | 5.198 | 5.288 | 5.039 | 5.412 | 5.366 | 5.360 | 5.792 | 5.568 | 5.425 | 5.154 | 5.386 | 5.125 | 4.876 | 5.042 | 5.307 | 5.475 | 5.368 | 5.426 |
| 4.446 | 4.437 | 4.614 | 4.844 | 4.384 | 4.573 | 4.953 | 4.471 | 4.735 | 5.238 | 5.197 | 4.962 | 5.053 | 4.714 | 4.762 | 4.662 | 4.892 | 4.538 | 5.106 | 4.782 |
| 4.766 | 4.758 | 4.656 | 4.721 | 4.701 | 4.883 | 4.824 | 4.888 | 5.022 | 4.802 | 4.786 | 4.658 | 5.206 | 4.561 | 4.255 | 4.226 | 4.658 | 4.665 | 4.804 | 4.839 |
| 3.678 | 3.927 | 4.049 | 4.455 | 3.886 | 4.090 | 4.300 | 3.958 | 3.667 | 4.451 | 4.615 | 4.420 | 3.711 | 4.391 | 5.250 | 5.261 | 4.338 | 4.521 | 4.370 | 4.417 |
| 4.984 | 4.927 | 4.126 | 4.003 | 4.053 | 4.818 | 4.149 | 4.529 | 5.274 | 4.688 | 4.382 | 4.019 | 5.142 | 4.436 | 3.849 | 4.859 | 5.183 | 4.081 | 4.370 | 4.917 |
| 5.032 | 5.091 | 5.384 | 5.574 | 5.070 | 5.427 | 5.368 | 5.267 | 5.362 | 5.359 | 5.356 | 5.472 | 5.512 | 5.604 | 5.535 | 5.872 | 5.649 | 5.683 | 5.469 | 5.556 |
| 7.085 | 7.192 | 7.284 | 7.268 | 6.698 | 7.214 | 7.260 | 7.143 | 6.981 | 7.279 | 7.295 | 7.277 | 7.153 | 7.393 | 7.162 | 7.447 | 7.256 | 7.265 | 7.419 | 7.431 |
| 6.530 | 6.556 | 6.909 | 7.069 | 6.216 | 6.603 | 6.988 | 6.531 | 6.418 | 6.858 | 7.021 | 7.065 | 6.306 | 7.131 | 6.989 | 7.114 | 7.066 | 6.888 | 7.234 | 7.144 |
| 6.414 | 6.246 | 7.078 | 6.994 | 6.734 | 6.954 | 7.008 | 7.001 | 6.828 | 6.956 | 7.060 | 7.283 | 6.525 | 7.319 | 7.146 | 7.438 | 7.512 | 6.754 | 7.206 | 7.390 |
| 5.649 | 5.784 | 5.969 | 6.108 | 5.508 | 5.745 | 6.133 | 5.768 | 5.762 | 6.158 | 6.352 | 6.120 | 6.019 | 6.126 | 6.016 | 6.312 | 5.859 | 6.528 | 6.198 | 6.044 |
| 6.892 | 6.910 | 7.257 | 7.414 | 7.035 | 7.314 | 7.195 | 7.154 | 7.056 | 7.226 | 7.294 | 7.593 | 7.525 | 7.597 | 7.570 | 7.693 | 7.812 | 7.126 | 7.506 | 7.611 |
| 5.081 | 4.894 | 4.707 | 4.604 | 4.832 | 5.261 | 4.617 | 4.762 | 4.716 | 4.401 | 4.834 | 4.830 | 6.801 | 4.734 | 4.964 | 5.517 | 4.933 | 5.336 | 4.887 | 4.796 |
| 6.448 | 6.443 | 7.122 | 6.843 | 7.110 | 6.820 | 7.499 | 6.780 | 7.236 | 7.573 | 7.611 | 7.075 | 6.607 | 7.070 | 6.480 | 6.591 | 6.968 | 6.626 | 7.302 | 7.179 |
| 4.573 | 4.632 | 4.813 | 5.183 | 4.613 | 4.560 | 5.087 | 4.489 | 4.870 | 5.272 | 5.477 | 5.151 | 4.971 | 4.827 | 5.199 | 5.081 | 4.953 | 4.905 | 5.520 | 4.906 |
| 7.593 | 7.666 | 7.975 | 7.764 | 7.859 | 8.129 | 8.266 | 7.827 | 8.568 | 8.060 | 8.230 | 7.805 | 8.264 | 7.546 | 7.310 | 7.605 | 7.923 | 7.785 | 7.693 | 7.954 |
| 6.619 | 6.533 | 7.432 | 7.107 | 7.726 | 7.459 | 7.540 | 7.223 | 7.197 | 7.523 | 7.596 | 7.433 | 7.105 | 7.443 | 7.423 | 7.232 | 7.338 | 7.480 | 7.205 | 7.342 |
| 3.605 | 3.959 | 4.317 | 4.907 | 3.875 | 3.528 | 4.130 | 3.730 | 3.704 | 2.809 | 4.048 | 4.232 | 4.459 | 4.523 | 5.400 | 5.948 | 3.778 | 5.459 | 4.695 | 4.274 |
| 3.591 | 3.602 | 4.125 | 3.972 | 4.175 | 3.804 | 4.284 | 3.827 | 3.945 | 4.157 | 4.505 | 4.366 | 3.479 | 4.243 | 4.230 | 4.571 | 4.271 | 4.006 | 4.254 | 4.265 |
| 5.463 | 5.453 | 6.039 | 5.659 | 6.425 | 6.126 | 6.062 | 6.028 | 6.054 | 5.817 | 6.172 | 5.906 | 6.320 | 6.212 | 6.383 | 6.676 | 6.077 | 6.946 | 5.877 | 6.148 |
| 3.808 | 4.249 | 4.067 | 4.624 | 3.687 | 3.350 | 4.309 | 3.680 | 3.780 | 3.724 | 4.458 | 4.020 | 4.869 | 4.294 | 5.082 | 5.568 | 4.589 | 5.535 | 4.768 | 4.364 |
| 8.299 | 8.314 | 8.504 | 8.397 | 8.248 | 8.617 | 8.522 | 8.491 | 8.302 | 8.533 | 8.411 | 8.603 | 8.246 | 8.671 | 8.343 | 8.468 | 8.695 | 8.363 | 8.531 | 8.705 |
| 5.265 | 5.490 | 6.209 | 5.941 | 6.829 | 6.172 | 6.477 | 6.131 | 6.506 | 6.698 | 6.580 | 6.153 | 6.244 | 6.321 | 6.399 | 6.127 | 6.044 | 6.605 | 6.374 | 6.223 |
| 5.045 | 5.007 | 5.162 | 5.080 | 4.931 | 5.239 | 5.087 | 5.383 | 5.290 | 5.381 | 5.082 | 4.914 | 5.481 | 5.139 | 4.731 | 4.991 | 5.346 | 5.495 | 5.192 | 5.402 |
| 3.331 | 3.129 | 3.528 | 3.582 | 3.750 | 3.372 | 3.321 | 3.881 | 3.066 | 4.317 | 3.524 | 3.118 | 3.737 | 3.690 | 3.641 | 3.350 | 3.331 | 5.294 | 3.554 | 3.548 |
| 5.156 | 5.117 | 4.942 | 5.147 | 4.999 | 5.244 | 5.015 | 4.937 | 5.342 | 4.949 | 5.173 | 5.196 | 5.262 | 5.058 | 5.245 | 5.186 | 5.423 | 4.946 | 5.245 | 5.243 |
| 3.499 | 3.654 | 4.060 | 4.120 | 4.240 | 3.713 | 4.222 | 4.028 | 4.153 | 4.121 | 4.531 | 4.149 | 4.239 | 4.377 | 4.304 | 4.280 | 4.108 | 4.233 | 4.306 | 3.926 |
| 6.113 | 6.338 | 6.063 | 6.040 | 5.696 | 5.938 | 6.208 | 5.761 | 5.649 | 6.189 | 6.303 | 6.255 | 5.796 | 6.203 | 6.688 | 6.792 | 6.048 | 6.629 | 6.264 | 6.253 |
| 6.024 | 5.948 | 6.915 | 6.600 | 6.942 | 6.481 | 6.793 | 6.920 | 6.204 | 6.979 | 6.674 | 6.716 | 6.602 | 6.965 | 7.224 | 6.859 | 6.764 | 7.790 | 6.753 | 6.765 |
| 6.278 | 6.338 | 5.502 | 6.040 | 5.326 | 6.027 | 5.536 | 6.016 | 6.047 | 5.588 | 5.396 | 5.588 | 6.059 | 5.606 | 5.245 | 5.554 | 6.078 | 5.567 | 5.891 | 6.044 |
| 6.357 | 6.182 | 5.477 | 5.455 | 5.377 | 5.988 | 5.192 | 5.985 | 5.856 | 6.210 | 5.278 | 5.510 | 6.388 | 5.699 | 5.087 | 4.405 | 5.266 | 4.996 | 5.361 | 5.777 |
| 6.995 | 6.903 | 7.985 | 7.733 | 8.225 | 7.983 | 7.954 | 7.678 | 7.755 | 7.811 | 7.926 | 7.619 | 8.221 | 7.695 | 8.120 | 7.636 | 7.829 | 8.117 | 7.410 | 7.711 |
| 7.084 | 6.665 | 7.455 | 6.524 | 7.914 | 7.723 | 7.274 | 7.594 | 7.612 | 7.270 | 7.205 | 7.241 | 7.319 | 7.474 | 7.038 | 6.951 | 7.127 | 7.443 | 6.765 | 7.500 |
| 4.007 | 4.055 | 4.327 | 4.242 | 4.325 | 4.284 | 4.445 | 4.096 | 4.412 | 4.490 | 4.632 | 4.245 | 4.530 | 4.402 | 4.764 | 5.341 | 4.519 | 5.086 | 4.509 | 4.519 |
| 6.862 | 6.897 | 7.081 | 6.467 | 7.231 | 7.528 | 7.257 | 7.118 | 7.589 | 7.517 | 7.414 | 7.323 | 7.109 | 7.489 | 7.122 | 7.690 | 7.696 | 6.821 | 7.261 | 7.509 |
| 6.253 | 6.173 | 6.101 | 6.336 | 6.517 | 6.132 | 5.881 | 6.455 | 6.103 | 5.955 | 5.933 | 6.252 | 6.539 | 6.229 | 6.578 | 6.249 | 6.153 | 6.449 | 6.070 | 6.260 |
| 4.793 | 4.897 | 5.711 | 6.017 | 5.522 | 5.420 | 5.707 | 5.426 | 5.336 | 5.814 | 5.831 | 5.838 | 4.803 | 5.964 | 6.043 | 5.728 | 6.071 | 5.451 | 5.948 | 5.809 |
| 5.725 | 5.316 | 5.296 | 5.394 | 5.748 | 5.516 | 5.188 | 5.704 | 5.668 | 5.354 | 5.290 | 5.425 | 5.771 | 5.354 | 5.280 | 5.276 | 5.548 | 5.412 | 5.275 | 5.567 |
| 5.539 | 5.604 | 4.965 | 4.974 | 5.449 | 5.366 | 4.991 | 5.515 | 5.426 | 5.300 | 4.968 | 5.232 | 5.484 | 5.152 | 5.875 | 5.841 | 6.034 | 5.872 | 5.870 | 5.450 |
| 4.422 | 4.455 | 4.871 | 4.606 | 4.804 | 4.692 | 4.797 | 4.715 | 4.507 | 5.113 | 4.815 | 4.742 | 4.564 | 4.629 | 4.604 | 4.614 | 4.952 | 4.467 | 4.695 | 4.951 |
| 4.779 | 4.885 | 4.198 | 4.161 | 4.494 | 4.836 | 4.340 | 4.605 | 4.942 | 4.530 | 4.602 | 4.629 | 4.557 | 4.577 | 4.950 | 4.771 | 5.079 | 4.262 | 5.525 | 4.987 |
| 7.096 | 7.094 | 7.604 | 6.937 | 7.304 | 7.703 | 7.686 | 7.426 | 7.500 | 7.542 | 7.690 | 7.677 | 7.187 | 7.667 | 7.271 | 7.700 | 7.291 | 7.409 | 7.491 | 7.735 |
| 6.181 | 5.968 | 6.408 | 5.763 | 6.770 | 6.919 | 6.283 | 6.475 | 6.362 | 6.809 | 6.246 | 6.504 | 6.217 | 6.594 | 5.940 | 5.514 | 5.912 | 5.088 | 5.546 | 6.339 |
| 6.387 | 6.245 | 6.138 | 6.206 | 6.254 | 6.966 | 6.461 | 6.151 | 7.148 | 7.080 | 7.004 | 6.801 | 6.893 | 7.138 | 6.613 | 7.421 | 7.654 | 5.395 | 6.739 | 7.047 |
| 3.798 | 3.926 | 4.871 | 4.784 | 4.615 | 4.233 | 4.837 | 4.334 | 4.082 | 4.884 | 5.102 | 5.192 | 3.891 | 4.946 | 5.478 | 5.522 | 4.891 | 5.323 | 5.049 | 4.662 |
| 6.101 | 5.972 | 6.834 | 6.878 | 6.924 | 6.881 | 7.022 | 6.703 | 6.781 | 6.875 | 7.067 | 6.872 | 6.676 | 6.921 | 7.151 | 7.216 | 7.424 | 7.000 | 6.756 | 6.835 |
| 6.748 | 6.465 | 7.424 | 7.052 | 7.732 | 7.486 | 7.336 | 7.455 | 7.405 | 7.317 | 7.402 | 7.526 | 7.251 | 7.724 | 7.637 | 7.787 | 7.685 | 7.827 | 7.330 | 7.573 |
| 6.529 | 6.335 | 7.004 | 6.871 | 7.351 | 7.121 | 6.712 | 7.180 | 6.984 | 6.980 | 6.673 | 7.087 | 7.293 | 7.286 | 7.451 | 7.221 | 7.180 | 7.111 | 6.833 | 7.006 |
| 6.138 | 6.407 | 7.003 | 7.316 | 7.202 | 6.573 | 6.918 | 6.759 | 6.387 | 7.026 | 7.057 | 7.175 | 6.745 | 7.388 | 7.793 | 7.615 | 7.482 | 7.279 | 7.270 | 7.215 |
| 4.468 | 4.490 | 4.045 | 4.257 | 4.333 | 4.036 | 4.089 | 4.171 | 3.858 | 4.264 | 4.603 | 4.364 | 4.426 | 4.466 | 5.038 | 5.250 | 4.147 | 4.269 | 4.255 | 4.341 |
| 4.231 | 5.164 | 5.686 | 5.788 | 6.094 | 4.841 | 6.119 | 4.995 | 5.523 | 6.206 | 6.570 | 5.875 | 5.102 | 5.961 | 6.558 | 6.668 | 5.851 | 6.404 | 6.133 | 5.736 |
| 7.210 | 7.085 | 7.413 | 6.836 | 7.641 | 7.576 | 7.244 | 7.548 | 7.508 | 7.355 | 7.279 | 7.380 | 7.396 | 7.494 | 6.931 | 7.023 | 7.215 | 7.271 | 7.033 | 7.489 |
| 5.725 | 5.345 | 5.280 | 5.178 | 5.809 | 5.737 | 5.166 | 5.761 | 5.809 | 5.312 | 5.021 | 5.314 | 6.488 | 5.453 | 5.088 | 5.043 | 5.533 | 4.952 | 5.112 | 5.537 |
| 2.575 | 2.814 | 3.127 | 3.183 | 3.580 | 2.839 | 3.181 | 3.030 | 2.915 | 3.180 | 3.361 | 3.061 | 3.000 | 3.424 | 4.017 | 4.291 | 3.409 | 4.361 | 3.470 | 3.121 |
| 6.445 | 6.327 | 5.681 | 5.351 | 5.945 | 6.246 | 5.570 | 6.323 | 6.242 | 5.747 | 5.444 | 5.566 | 6.839 | 5.896 | 5.305 | 5.078 | 5.966 | 5.195 | 5.384 | 6.100 |
| 3.033 | 3.339 | 4.171 | 4.347 | 4.600 | 3.910 | 4.428 | 3.631 | 3.384 | 4.446 | 4.976 | 4.835 | 3.292 | 4.498 | 5.738 | 6.061 | 4.286 | 5.403 | 4.386 | 4.088 |
| 4.368 | 4.287 | 4.592 | 4.204 | 4.409 | 4.726 | 4.675 | 4.544 | 4.683 | 4.433 | 4.539 | 4.619 | 4.581 | 4.586 | 4.559 | 4.804 | 4.772 | 4.645 | 4.403 | 4.743 |
| 3.674 | 3.793 | 4.925 | 4.811 | 5.433 | 4.439 | 4.813 | 4.662 | 4.662 | 5.327 | 5.402 | 5.054 | 4.104 | 5.390 | 5.588 | 5.457 | 4.887 | 6.323 | 4.815 | 5.190 |
| 4.359 | 4.308 | 4.141 | 4.590 | 4.566 | 4.262 | 4.295 | 4.381 | 4.303 | 4.573 | 4.653 | 4.571 | 4.428 | 4.428 | 4.911 | 4.866 | 4.580 | 4.748 | 4.491 | 4.406 |
| 4.698 | 4.706 | 4.685 | 4.978 | 4.933 | 4.431 | 4.941 | 4.596 | 4.372 | 4.914 | 5.313 | 5.033 | 4.442 | 5.082 | 5.933 | 6.255 | 5.061 | 5.995 | 5.117 | 5.067 |
| 3.871 | 3.863 | 4.768 | 4.469 | 5.043 | 4.511 | 4.949 | 4.332 | 4.404 | 4.713 | 5.146 | 5.023 | 4.193 | 5.025 | 5.582 | 6.167 | 4.910 | 5.479 | 4.756 | 4.736 |
| 6.234 | 6.222 | 7.010 | 6.253 | 7.194 | 6.898 | 6.900 | 6.872 | 6.985 | 6.960 | 6.943 | 6.915 | 6.448 | 7.126 | 6.827 | 7.039 | 6.798 | 7.490 | 6.563 | 7.022 |
| 4.039 | 4.108 | 4.968 | 4.946 | 5.847 | 4.662 | 5.104 | 4.711 | 5.284 | 5.416 | 5.441 | 4.903 | 5.251 | 5.007 | 5.423 | 5.215 | 4.707 | 6.289 | 4.988 | 4.824 |
| 5.573 | 5.179 | 5.576 | 4.663 | 6.443 | 5.620 | 5.499 | 5.596 | 5.919 | 5.308 | 5.293 | 5.500 | 6.416 | 5.636 | 5.988 | 6.498 | 5.539 | 7.080 | 4.767 | 5.389 |
| 5.005 | 5.022 | 4.862 | 5.062 | 5.239 | 4.859 | 4.792 | 5.031 | 4.997 | 4.956 | 4.953 | 4.999 | 4.902 | 5.058 | 5.233 | 5.312 | 5.153 | 5.296 | 4.937 | 5.096 |
| 5.339 | 5.220 | 5.241 | 5.095 | 5.637 | 5.199 | 5.166 | 5.247 | 5.106 | 5.029 | 5.122 | 5.456 | 5.380 | 5.508 | 5.780 | 5.818 | 5.544 | 5.568 | 5.108 | 5.407 |
| 3.257 | 3.300 | 3.550 | 3.605 | 3.656 | 3.484 | 3.688 | 3.322 | 3.361 | 3.449 | 3.661 | 3.573 | 3.418 | 3.818 | 4.423 | 4.965 | 3.792 | 4.479 | 3.787 | 3.560 |
| 4.476 | 4.511 | 3.940 | 3.717 | 4.509 | 4.332 | 3.904 | 4.384 | 4.430 | 3.938 | 4.056 | 4.115 | 4.385 | 4.094 | 4.255 | 3.961 | 4.253 | 4.208 | 4.368 | 4.331 |
| 3.098 | 3.146 | 3.543 | 3.811 | 3.866 | 3.263 | 3.652 | 3.237 | 3.255 | 3.683 | 3.783 | 3.948 | 3.257 | 3.746 | 4.535 | 4.542 | 3.649 | 3.955 | 3.555 | 3.443 |
| 4.443 | 4.149 | 4.157 | 3.437 | 4.637 | 4.691 | 3.780 | 4.512 | 4.428 | 4.000 | 3.568 | 4.146 | 5.010 | 4.256 | 3.708 | 3.351 | 3.745 | 3.329 | 3.267 | 3.965 |
| 4.374 | 4.124 | 4.489 | 3.811 | 4.864 | 4.418 | 4.035 | 4.897 | 4.777 | 4.101 | 3.823 | 3.961 | 5.116 | 4.579 | 4.034 | 4.261 | 4.255 | 5.091 | 3.830 | 4.274 |
| 3.608 | 3.624 | 4.011 | 3.910 | 4.786 | 3.967 | 4.290 | 3.903 | 4.323 | 4.332 | 4.441 | 4.238 | 4.219 | 4.214 | 4.350 | 4.330 | 4.313 | 4.589 | 4.183 | 4.042 |
| 4.537 | 4.229 | 4.847 | 4.079 | 5.563 | 5.124 | 4.541 | 4.973 | 5.286 | 4.361 | 4.355 | 4.707 | 5.108 | 5.013 | 4.523 | 4.756 | 4.421 | 5.624 | 4.101 | 4.706 |
| 3.652 | 3.574 | 4.056 | 3.873 | 4.560 | 4.293 | 4.027 | 4.109 | 4.342 | 4.226 | 4.269 | 4.189 | 4.315 | 4.410 | 4.336 | 4.417 | 4.026 | 4.467 | 3.733 | 4.131 |
| 3.650 | 4.125 | 4.935 | 4.235 | 5.436 | 4.768 | 4.794 | 4.735 | 4.635 | 4.809 | 4.850 | 5.167 | 4.398 | 5.172 | 5.330 | 5.115 | 4.664 | 5.145 | 4.368 | 4.709 |
| 3.374 | 3.460 | 4.378 | 4.343 | 5.346 | 4.229 | 4.600 | 4.288 | 4.660 | 4.759 | 4.999 | 4.685 | 4.215 | 4.726 | 4.781 | 5.178 | 4.421 | 5.202 | 4.452 | 4.435 |

| 46124A Non-progre | 43747D Non-progre | 44466D Non-progre | 46093A Non-progre | 4423A Non-progre | 43311A Non-progre | 43005A Non-progre | 46103A Non-progre | 43081A Non-progre | 44088A Non-progre | 42287J Non-progre | 43366A Non-progre | 43061A Non-progre | 44045D Non-progre | 43068A Non-progre | 46105A Non-progre | 43174A Non-progre | 43252J Non-progre | 43262A Non-progre |
|---|---|---|---|---|---|---|---|---|---|---|---|---|---|---|---|---|---|---|
| NP | NP | NP | NP | NP | NP | NP | NP | NP | NP | NP | NP | NP | NP | NP | NP | NP | NP | NP |
| 3.834 | 3.609 | 3.446 | 3.588 | 3.948 | 3.749 | 3.837 | 3.907 | 3.656 | 3.717 | 3.799 | 3.758 | 3.805 | 3.736 | 3.825 | 4.163 | 3.832 | 3.800 | 3.638 |
| 5.505 | 5.216 | 5.262 | 4.607 | 5.678 | 5.415 | 5.194 | 5.277 | 5.205 | 5.330 | 5.492 | 5.531 | 5.669 | 5.604 | 5.358 | 5.653 | 5.053 | 5.491 | 5.210 |
| 4.808 | 4.512 | 4.552 | 4.413 | 5.001 | 4.668 | 4.733 | 4.616 | 4.925 | 4.543 | 5.013 | 4.699 | 4.856 | 4.737 | 4.683 | 4.564 | 4.297 | 4.431 | 4.313 |
| 7.757 | 7.272 | 6.993 | 7.055 | 7.521 | 7.529 | 7.192 | 7.468 | 7.802 | 7.110 | 7.244 | 7.451 | 7.504 | 7.858 | 7.532 | 7.569 | 6.976 | 7.429 | 7.318 |
| 6.850 | 6.717 | 7.214 | 7.087 | 7.422 | 7.416 | 6.790 | 7.197 | 7.499 | 7.357 | 7.718 | 7.183 | 7.575 | 7.452 | 7.331 | 7.527 | 8.057 | 7.604 | 7.763 |
| 5.627 | 6.365 | 5.617 | 5.037 | 6.126 | 5.945 | 5.304 | 5.809 | 5.659 | 4.789 | 5.807 | 5.308 | 6.175 | 6.074 | 5.972 | 6.724 | 7.331 | 5.078 | 6.165 |
| 8.041 | 8.243 | 8.341 | 8.456 | 8.610 | 8.653 | 8.344 | 8.612 | 8.532 | 8.668 | 8.685 | 8.091 | 8.491 | 8.411 | 8.557 | 8.675 | 8.908 | 8.721 | 8.651 |
| 4.660 | 4.792 | 4.618 | 4.462 | 5.305 | 5.105 | 4.697 | 5.042 | 4.774 | 5.205 | 5.119 | 4.640 | 5.180 | 5.420 | 5.047 | 5.742 | 5.840 | 5.656 | 5.695 |
| 5.691 | 5.585 | 5.695 | 5.390 | 5.950 | 5.895 | 5.663 | 5.959 | 5.869 | 5.774 | 6.103 | 5.856 | 6.283 | 6.189 | 6.037 | 6.596 | 5.840 | 6.140 | 6.053 |
| 3.944 | 4.031 | 4.286 | 3.907 | 3.814 | 4.245 | 3.850 | 4.084 | 3.969 | 4.222 | 3.950 | 4.211 | 3.938 | 4.490 | 4.134 | 4.106 | 5.057 | 4.447 | 4.697 |
| 6.114 | 5.840 | 5.612 | 6.298 | 6.075 | 5.704 | 6.225 | 6.134 | 6.192 | 6.393 | 6.156 | 6.113 | 5.935 | 5.793 | 6.220 | 6.144 | 5.196 | 5.783 | 5.395 |
| 7.969 | 8.047 | 8.074 | 7.312 | 7.810 | 8.127 | 7.615 | 8.025 | 7.906 | 7.571 | 8.031 | 8.071 | 7.863 | 8.326 | 7.969 | 7.891 | 8.582 | 8.349 | 8.421 |
| 5.378 | 5.100 | 5.325 | 5.378 | 6.007 | 5.772 | 5.458 | 5.511 | 6.144 | 5.713 | 6.364 | 5.864 | 6.307 | 6.156 | 5.901 | 6.210 | 6.297 | 5.841 | 6.040 |
| 5.296 | 5.158 | 4.924 | 5.477 | 5.613 | 5.184 | 5.367 | 5.119 | 6.058 | 5.315 | 6.266 | 5.412 | 5.906 | 5.524 | 5.329 | 5.534 | 3.877 | 4.921 | 4.905 |
| 4.817 | 4.887 | 4.726 | 4.744 | 4.891 | 4.722 | 4.821 | 4.823 | 4.670 | 4.541 | 4.745 | 4.857 | 4.889 | 4.870 | 4.789 | 5.034 | 4.223 | 4.778 | 4.777 |
| 6.501 | 6.480 | 7.077 | 6.094 | 6.859 | 6.902 | 6.448 | 6.804 | 5.682 | 7.110 | 5.927 | 6.739 | 6.329 | 6.732 | 6.764 | 6.769 | 7.914 | 7.185 | 7.414 |
| 4.981 | 4.552 | 4.450 | 4.070 | 5.602 | 4.918 | 4.903 | 4.769 | 5.323 | 4.598 | 5.541 | 4.748 | 5.721 | 4.848 | 5.041 | 5.200 | 4.091 | 4.759 | 4.444 |
| 5.655 | 5.623 | 5.486 | 5.235 | 5.574 | 5.722 | 5.337 | 5.597 | 5.396 | 5.552 | 5.539 | 5.565 | 5.400 | 5.665 | 5.335 | 5.622 | 6.458 | 5.847 | 6.019 |
| 6.088 | 6.235 | 6.772 | 6.001 | 5.554 | 6.605 | 5.731 | 6.176 | 6.571 | 6.077 | 6.606 | 6.837 | 6.631 | 7.083 | 6.159 | 6.445 | 7.609 | 6.764 | 7.361 |
| 4.118 | 4.173 | 4.319 | 4.025 | 4.628 | 4.592 | 4.165 | 4.551 | 3.614 | 5.377 | 3.774 | 4.237 | 4.127 | 4.338 | 4.757 | 4.910 | 5.923 | 5.319 | 5.201 |
| 5.109 | 5.088 | 5.143 | 5.094 | 5.612 | 5.440 | 5.167 | 5.214 | 5.116 | 5.363 | 5.233 | 5.235 | 5.289 | 5.398 | 5.218 | 5.569 | 6.325 | 5.605 | 5.887 |
| 8.404 | 8.253 | 8.034 | 8.690 | 7.717 | 7.675 | 8.154 | 7.866 | 7.899 | 8.388 | 7.935 | 8.627 | 7.769 | 7.948 | 8.242 | 8.059 | 7.146 | 7.490 | 7.605 |
| 7.904 | 7.852 | 7.867 | 7.598 | 7.860 | 7.886 | 7.730 | 7.925 | 7.937 | 7.686 | 8.050 | 8.054 | 7.883 | 8.141 | 7.845 | 7.766 | 8.163 | 8.015 | 8.045 |
| 4.724 | 4.660 | 4.284 | 4.780 | 5.335 | 4.604 | 5.071 | 4.920 | 4.943 | 4.984 | 4.934 | 4.654 | 5.056 | 4.646 | 4.891 | 5.356 | 4.515 | 4.795 | 4.486 |
| 5.689 | 5.389 | 5.365 | 5.585 | 5.600 | 5.302 | 5.537 | 5.422 | 5.354 | 5.639 | 5.433 | 5.533 | 5.418 | 5.340 | 5.479 | 5.650 | 5.337 | 5.299 | 5.204 |
| 4.702 | 4.813 | 5.029 | 4.802 | 4.675 | 4.770 | 4.745 | 4.968 | 4.953 | 4.572 | 5.109 | 4.905 | 4.978 | 5.138 | 4.928 | 5.258 | 5.069 | 4.995 | 4.955 |
| 5.037 | 4.891 | 4.603 | 5.030 | 5.156 | 4.579 | 5.158 | 4.713 | 4.940 | 4.818 | 5.080 | 5.007 | 5.025 | 4.841 | 4.973 | 4.992 | 4.288 | 4.643 | 4.359 |
| 4.111 | 4.218 | 4.762 | 3.924 | 4.093 | 4.621 | 3.818 | 4.255 | 4.404 | 4.229 | 4.702 | 4.396 | 4.450 | 4.988 | 4.336 | 4.480 | 6.080 | 4.935 | 5.850 |
| 4.776 | 5.402 | 5.170 | 4.727 | 4.923 | 4.194 | 5.017 | 4.695 | 4.275 | 4.215 | 4.782 | 4.710 | 4.788 | 4.401 | 4.941 | 4.999 | 4.705 | 4.021 | 4.594 |
| 5.458 | 5.516 | 5.457 | 5.350 | 5.524 | 5.703 | 5.424 | 5.793 | 5.366 | 5.749 | 5.518 | 5.669 | 5.435 | 5.790 | 5.672 | 5.841 | 6.163 | 5.674 | 5.737 |
| 7.425 | 7.363 | 7.335 | 6.850 | 7.559 | 7.498 | 7.226 | 7.588 | 7.375 | 7.289 | 7.482 | 7.551 | 7.382 | 7.627 | 7.353 | 7.449 | 7.998 | 7.705 | 7.699 |
| 6.817 | 7.160 | 7.096 | 6.442 | 6.942 | 7.328 | 6.647 | 7.309 | 6.869 | 7.102 | 7.097 | 6.906 | 6.851 | 7.288 | 6.928 | 7.158 | 7.513 | 7.431 | 7.497 |
| 7.094 | 7.377 | 7.503 | 7.279 | 6.887 | 7.482 | 7.085 | 7.567 | 7.408 | 7.484 | 7.483 | 7.394 | 7.306 | 7.725 | 7.528 | 7.498 | 7.722 | 7.643 | 7.808 |
| 6.134 | 6.051 | 5.885 | 5.566 | 6.091 | 6.208 | 5.759 | 6.055 | 5.777 | 6.162 | 5.944 | 6.275 | 5.913 | 6.424 | 5.965 | 6.247 | 6.866 | 6.532 | 6.548 |
| 7.212 | 7.513 | 7.754 | 7.564 | 7.450 | 7.815 | 7.418 | 7.787 | 7.590 | 7.720 | 7.727 | 7.491 | 7.597 | 7.838 | 7.679 | 7.862 | 8.398 | 7.924 | 8.050 |
| 4.616 | 4.832 | 5.058 | 5.079 | 5.268 | 5.128 | 4.971 | 4.999 | 4.877 | 5.193 | 4.910 | 4.916 | 4.786 | 4.886 | 4.816 | 5.258 | 6.023 | 5.372 | 5.348 |
| 7.508 | 7.419 | 7.210 | 6.918 | 6.947 | 7.053 | 7.098 | 7.264 | 7.083 | 7.024 | 7.154 | 7.534 | 7.203 | 7.503 | 7.412 | 7.506 | 6.638 | 7.344 | 7.091 |
| 4.877 | 4.867 | 4.870 | 4.759 | 4.909 | 5.137 | 4.826 | 5.205 | 4.850 | 4.901 | 4.889 | 4.944 | 4.952 | 5.302 | 5.055 | 5.595 | 5.743 | 5.723 | 5.631 |
| 8.475 | 8.067 | 7.660 | 8.349 | 8.014 | 7.652 | 8.009 | 7.828 | 7.950 | 8.226 | 7.918 | 8.452 | 8.123 | 8.055 | 7.997 | 8.124 | 7.751 | 7.725 | 7.683 |
| 7.788 | 7.731 | 7.744 | 8.040 | 7.127 | 7.462 | 7.422 | 7.366 | 7.657 | 7.699 | 7.542 | 8.273 | 7.586 | 7.940 | 7.621 | 7.328 | 7.579 | 7.705 | 7.841 |
| 4.195 | 3.870 | 4.122 | 3.944 | 4.513 | 5.041 | 4.182 | 4.915 | 3.716 | 4.902 | 3.607 | 4.343 | 3.895 | 4.810 | 4.262 | 4.218 | 6.164 | 5.412 | 5.846 |
| 4.336 | 4.301 | 4.472 | 4.161 | 3.985 | 4.387 | 3.939 | 4.345 | 4.199 | 4.280 | 4.339 | 4.472 | 4.314 | 4.834 | 3.925 | 4.235 | 4.884 | 4.703 | 4.905 |
| 6.384 | 6.500 | 6.850 | 6.703 | 5.888 | 6.244 | 6.625 | 6.336 | 6.148 | 6.388 | 5.962 | 6.820 | 6.122 | 6.344 | 6.523 | 6.131 | 6.462 | 6.336 | 6.608 |
| 4.478 | 4.490 | 4.811 | 4.106 | 4.713 | 4.688 | 4.086 | 4.656 | 3.391 | 5.137 | 3.549 | 4.703 | 3.999 | 4.853 | 4.527 | 4.737 | 6.723 | 5.480 | 5.790 |
| 8.726 | 8.775 | 8.803 | 8.730 | 8.802 | 8.760 | 8.638 | 8.823 | 8.932 | 8.845 | 8.982 | 8.920 | 8.858 | 8.992 | 8.870 | 8.763 | 8.989 | 8.875 | 8.881 |
| 6.668 | 6.733 | 6.734 | 6.678 | 6.240 | 6.239 | 6.700 | 6.384 | 6.689 | 6.224 | 6.696 | 7.187 | 7.014 | 6.720 | 6.753 | 6.734 | 6.032 | 6.341 | 6.580 |
| 5.589 | 5.438 | 5.493 | 5.684 | 5.420 | 5.283 | 5.668 | 5.669 | 5.465 | 5.681 | 5.429 | 5.484 | 5.173 | 5.394 | 5.693 | 5.351 | 4.919 | 5.488 | 5.215 |
| 4.062 | 3.689 | 3.979 | 4.443 | 3.636 | 3.463 | 4.006 | 4.171 | 4.089 | 4.130 | 3.776 | 3.534 | 3.080 | 3.546 | 5.196 | 3.128 | 3.065 | 4.568 | 3.735 |
| 5.099 | 5.195 | 5.216 | 5.355 | 5.383 | 5.332 | 5.251 | 5.434 | 5.263 | 5.435 | 5.354 | 5.496 | 5.341 | 5.350 | 5.540 | 5.766 | 5.571 | 5.684 | 5.628 |
| 4.394 | 4.152 | 4.427 | 4.431 | 4.006 | 4.528 | 4.329 | 4.279 | 4.295 | 4.556 | 4.392 | 4.450 | 4.336 | 4.462 | 4.847 | 4.845 | 4.420 | 4.490 | 4.834 |
| 6.282 | 6.309 | 6.617 | 5.511 | 6.690 | 6.363 | 6.119 | 6.328 | 6.285 | 6.193 | 6.438 | 6.614 | 6.607 | 6.824 | 6.033 | 6.189 | 7.387 | 6.636 | 6.973 |
| 7.262 | 7.139 | 7.537 | 7.051 | 6.482 | 7.149 | 7.199 | 7.207 | 7.258 | 7.147 | 6.970 | 7.392 | 6.493 | 7.131 | 7.670 | 6.261 | 6.333 | 7.532 | 7.377 |
| 5.816 | 5.642 | 5.278 | 6.038 | 6.421 | 5.940 | 6.200 | 6.144 | 6.141 | 6.483 | 6.169 | 5.810 | 6.179 | 5.700 | 6.103 | 6.424 | 5.911 | 6.095 | 5.435 |
| 5.449 | 5.157 | 4.720 | 5.642 | 6.340 | 5.258 | 5.897 | 5.208 | 6.569 | 5.196 | 6.645 | 5.414 | 6.385 | 5.139 | 5.783 | 6.157 | 3.765 | 4.753 | 4.657 |
| 8.413 | 7.874 | 8.029 | 8.901 | 7.438 | 8.029 | 8.269 | 8.098 | 8.184 | 8.658 | 8.013 | 8.824 | 7.825 | 8.177 | 8.357 | 8.023 | 7.242 | 8.048 | 8.103 |
| 7.839 | 7.820 | 7.828 | 7.648 | 7.241 | 7.300 | 7.826 | 7.509 | 8.022 | 7.199 | 7.760 | 8.239 | 7.855 | 7.690 | 7.844 | 7.236 | 6.808 | 7.230 | 7.291 |
| 4.689 | 4.685 | 4.770 | 4.530 | 4.568 | 4.596 | 4.685 | 4.651 | 4.390 | 4.718 | 4.367 | 4.795 | 4.611 | 4.767 | 4.613 | 4.673 | 5.370 | 4.798 | 5.140 |
| 7.447 | 8.181 | 8.275 | 7.201 | 7.124 | 7.396 | 7.475 | 7.534 | 7.464 | 6.587 | 7.696 | 8.000 | 7.886 | 7.792 | 7.837 | 7.650 | 8.494 | 7.211 | 8.078 |
| 6.293 | 6.074 | 6.131 | 7.007 | 6.482 | 6.439 | 6.723 | 6.554 | 7.032 | 7.107 | 6.718 | 6.584 | 6.636 | 6.448 | 7.052 | 6.741 | 5.500 | 6.825 | 6.614 |
| 5.853 | 5.826 | 5.858 | 5.569 | 5.294 | 6.196 | 5.339 | 6.072 | 5.720 | 6.230 | 5.842 | 6.176 | 5.743 | 6.487 | 6.025 | 6.289 | 6.551 | 6.234 | 6.690 |
| 5.554 | 5.608 | 5.336 | 6.207 | 5.679 | 5.372 | 5.878 | 5.694 | 6.317 | 6.043 | 6.029 | 5.824 | 6.031 | 5.665 | 6.227 | 5.897 | 5.177 | 5.687 | 5.570 |
| 5.237 | 5.816 | 5.806 | 5.423 | 5.744 | 6.106 | 6.251 | 6.176 | 5.783 | 5.454 | 5.944 | 5.481 | 6.020 | 5.370 | 5.988 | 6.101 | 6.530 | 6.827 | 6.100 |
| 4.982 | 5.071 | 4.942 | 5.091 | 4.692 | 4.743 | 4.780 | 4.877 | 5.068 | 4.909 | 5.027 | 5.259 | 4.682 | 5.134 | 5.315 | 4.751 | 4.483 | 5.015 | 4.812 |
| 4.431 | 5.282 | 5.183 | 4.643 | 5.136 | 5.286 | 5.285 | 5.579 | 4.921 | 4.259 | 5.162 | 4.792 | 5.210 | 4.873 | 4.932 | 5.021 | 6.134 | 5.596 | 5.658 |
| 7.829 | 7.899 | 8.057 | 7.357 | 7.556 | 7.732 | 7.579 | 7.773 | 7.715 | 7.513 | 7.710 | 8.008 | 7.681 | 8.022 | 7.651 | 7.474 | 8.395 | 7.882 | 8.024 |
| 6.666 | 6.600 | 6.350 | 6.680 | 6.362 | 6.117 | 6.545 | 6.174 | 7.290 | 6.196 | 7.154 | 7.047 | 6.819 | 6.598 | 6.577 | 6.189 | 4.782 | 5.546 | 5.716 |
| 6.842 | 7.860 | 7.919 | 6.837 | 6.600 | 6.787 | 6.786 | 6.988 | 6.428 | 6.053 | 7.314 | 7.222 | 7.495 | 7.119 | 7.387 | 7.597 | 8.518 | 6.143 | 7.772 |
| 4.677 | 4.754 | 5.555 | 4.423 | 4.475 | 5.452 | 4.575 | 5.109 | 5.116 | 4.690 | 5.188 | 5.487 | 5.291 | 5.886 | 4.903 | 5.082 | 6.108 | 5.623 | 6.137 |
| 7.223 | 7.041 | 7.323 | 7.744 | 6.531 | 7.052 | 7.073 | 7.148 | 7.179 | 7.434 | 7.183 | 7.581 | 7.164 | 7.334 | 7.470 | 7.334 | 7.552 | 7.118 | 7.601 |
| 7.685 | 7.875 | 8.124 | 8.066 | 6.847 | 7.722 | 7.902 | 7.842 | 7.989 | 7.778 | 7.840 | 8.134 | 7.967 | 8.036 | 8.031 | 7.646 | 7.939 | 7.720 | 8.119 |
| 7.092 | 7.517 | 7.603 | 7.981 | 6.970 | 7.312 | 7.513 | 7.310 | 7.665 | 7.592 | 7.721 | 7.625 | 7.547 | 7.390 | 7.693 | 7.342 | 7.205 | 7.114 | 7.524 |
| 6.900 | 7.331 | 7.727 | 7.398 | 6.625 | 7.572 | 7.051 | 7.483 | 7.081 | 7.718 | 7.286 | 7.388 | 7.234 | 7.871 | 7.649 | 7.613 | 8.110 | 7.735 | 8.175 |
| 4.588 | 4.336 | 4.453 | 4.099 | 4.521 | 4.507 | 4.145 | 4.196 | 4.375 | 4.457 | 4.604 | 4.349 | 4.488 | 4.859 | 4.451 | 4.403 | 5.424 | 4.568 | 5.399 |
| 5.853 | 5.944 | 6.397 | 5.715 | 5.100 | 6.083 | 5.785 | 6.065 | 5.670 | 5.623 | 5.739 | 6.247 | 6.288 | 6.382 | 6.149 | 6.371 | 6.686 | 6.466 | 7.126 |
| 7.768 | 7.727 | 7.908 | 7.622 | 7.433 | 7.455 | 7.770 | 7.529 | 7.853 | 7.499 | 7.711 | 8.126 | 7.786 | 7.841 | 7.845 | 7.508 | 7.178 | 7.344 | 7.487 |
| 5.781 | 5.770 | 5.519 | 6.699 | 5.811 | 5.321 | 6.183 | 5.691 | 6.213 | 6.346 | 5.949 | 5.927 | 5.980 | 5.606 | 6.267 | 5.926 | 4.420 | 5.385 | 5.165 |
| 3.384 | 3.377 | 3.678 | 3.434 | 3.138 | 3.634 | 3.382 | 3.480 | 3.280 | 3.631 | 3.187 | 3.744 | 3.578 | 3.606 | 3.696 | 3.480 | 4.395 | 3.833 | 4.446 |
| 6.240 | 5.994 | 5.848 | 6.824 | 6.354 | 5.718 | 6.611 | 5.990 | 6.668 | 6.285 | 6.619 | 6.380 | 6.462 | 5.905 | 6.466 | 6.165 | 4.465 | 5.143 | 5.182 |
| 4.125 | 4.294 | 5.076 | 4.167 | 3.844 | 4.970 | 3.904 | 4.383 | 4.852 | 4.329 | 4.682 | 4.868 | 4.971 | 5.435 | 4.310 | 4.568 | 6.194 | 5.313 | 6.273 |
| 4.915 | 4.858 | 5.118 | 4.995 | 4.627 | 4.755 | 4.814 | 4.863 | 4.820 | 4.852 | 4.798 | 4.991 | 4.790 | 5.022 | 4.900 | 4.708 | 5.237 | 4.806 | 5.048 |
| 5.279 | 5.138 | 5.647 | 5.238 | 4.358 | 5.348 | 5.165 | 5.413 | 4.934 | 5.535 | 4.662 | 5.797 | 5.015 | 5.674 | 5.348 | 5.268 | 6.249 | 5.666 | 5.992 |
| 4.431 | 4.398 | 4.555 | 4.930 | 4.618 | 4.523 | 4.508 | 4.666 | 4.912 | 5.022 | 4.847 | 4.665 | 4.996 | 4.829 | 4.890 | 5.116 | 5.097 | 4.980 | 5.090 |
| 4.928 | 4.919 | 5.473 | 4.989 | 5.060 | 5.240 | 4.882 | 5.205 | 5.095 | 5.407 | 5.020 | 5.275 | 5.440 | 5.655 | 5.060 | 5.254 | 6.786 | 5.744 | 6.437 |
| 4.832 | 5.071 | 5.713 | 4.972 | 4.373 | 5.162 | 4.777 | 5.029 | 5.148 | 4.840 | 4.963 | 5.212 | 5.255 | 5.550 | 5.076 | 4.913 | 6.028 | 5.309 | 6.097 |
| 7.433 | 7.163 | 7.544 | 7.085 | 6.842 | 6.983 | 7.322 | 7.186 | 7.415 | 7.049 | 7.054 | 7.770 | 7.267 | 7.453 | 7.318 | 7.004 | 7.549 | 7.283 | 7.304 |
| 5.405 | 5.178 | 5.521 | 5.833 | 4.896 | 4.834 | 5.598 | 5.089 | 5.413 | 5.320 | 5.217 | 5.622 | 5.783 | 5.605 | 5.328 | 5.611 | 5.259 | 5.313 | 5.476 |
| 5.832 | 6.005 | 6.585 | 6.810 | 5.380 | 5.616 | 6.221 | 5.134 | 5.846 | 5.931 | 5.488 | 6.343 | 5.947 | 6.144 | 6.026 | 5.287 | 5.712 | 5.418 | 5.920 |
| 5.243 | 5.061 | 5.200 | 5.490 | 5.356 | 5.139 | 5.378 | 5.360 | 5.675 | 5.797 | 5.366 | 5.426 | 5.559 | 5.317 | 5.591 | 5.635 | 5.723 | 5.685 | 5.372 |
| 5.319 | 5.618 | 5.944 | 6.000 | 5.367 | 5.470 | 5.515 | 5.512 | 5.937 | 5.936 | 5.764 | 5.646 | 5.878 | 5.792 | 5.863 | 5.667 | 5.384 | 5.484 | 5.910 |
| 3.755 | 3.941 | 4.375 | 3.788 | 3.711 | 3.974 | 3.665 | 3.861 | 3.731 | 4.057 | 3.662 | 4.143 | 3.830 | 4.255 | 3.946 | 3.934 | 5.179 | 4.321 | 4.998 |
| 4.285 | 4.463 | 4.519 | 4.702 | 4.599 | 4.412 | 4.915 | 4.722 | 4.721 | 4.421 | 4.567 | 4.546 | 4.678 | 4.487 | 4.638 | 4.377 | 4.198 | 4.669 | 4.602 |
| 3.460 | 3.514 | 3.957 | 3.817 | 3.748 | 3.853 | 3.631 | 3.689 | 4.417 | 4.013 | 4.150 | 3.933 | 4.296 | 4.202 | 3.997 | 4.150 | 4.709 | 4.364 | 4.837 |
| 4.434 | 4.621 | 4.510 | 5.117 | 4.391 | 3.757 | 4.926 | 4.012 | 5.289 | 4.134 | 4.894 | 4.757 | 4.693 | 4.031 | 4.444 | 3.921 | 2.705 | 3.341 | 3.438 |
| 5.031 | 4.836 | 4.896 | 5.385 | 4.574 | 4.276 | 5.419 | 4.576 | 5.290 | 4.925 | 4.581 | 5.315 | 4.798 | 4.366 | 4.895 | 4.278 | 3.899 | 4.178 | 4.057 |
| 4.473 | 4.487 | 5.026 | 4.655 | 4.151 | 4.184 | 4.521 | 4.371 | 4.632 | 4.479 | 4.451 | 4.830 | 5.055 | 4.581 | 4.572 | 4.803 | 4.655 | 4.421 | 4.713 |
| 5.219 | 5.115 | 5.401 | 5.605 | 4.631 | 4.601 | 5.454 | 4.864 | 5.624 | 4.989 | 5.045 | 5.682 | 5.703 | 5.202 | 5.024 | 4.710 | 4.551 | 4.428 | 4.705 |
| 4.372 | 4.377 | 4.804 | 4.801 | 3.904 | 4.138 | 4.489 | 4.337 | 4.607 | 4.382 | 4.303 | 4.820 | 4.650 | 4.540 | 4.508 | 4.421 | 4.545 | 4.313 | 4.440 |
| 5.069 | 5.068 | 5.746 | 5.156 | 4.421 | 5.167 | 4.996 | 4.932 | 5.554 | 4.905 | 5.327 | 5.575 | 5.439 | 5.365 | 4.960 | 4.700 | 5.149 | 4.849 | 5.331 |
| 4.952 | 4.658 | 5.278 | 5.043 | 4.062 | 4.443 | 4.947 | 4.631 | 5.041 | 4.856 | 4.584 | 5.370 | 5.478 | 5.103 | 4.784 | 5.121 | 5.292 | 4.624 | 5.167 |

| 43260A Non-progre | 43907A Non-progre | 46171A Non-progre | 46071A Non-progre | 43079A Non-progre | 44053A Non-progre | 41836J Non-progre | 43176A Non-progre | 43928A Non-progre | 43382A Non-progre | 46142A Non-progre | 43097A Non-progre | 43442D Non-progre | 43930A Non-progre | 43643A Non-progre | 43535D Non-progre | 43326A Non-progre | 43895D Non-progre |
|---|---|---|---|---|---|---|---|---|---|---|---|---|---|---|---|---|---|
| NP | NP | NP | NP | NP | NP | NP | NP | NP | NP | NP | NP | NP | NP | NP | NP | NP | NP |
| 3.570 | 3.418 | 3.983 | 3.866 | 3.622 | 3.942 | 3.832 | 3.808 | 3.991 | 3.720 | 3.888 | 3.993 | 3.904 | 3.867 | 3.735 | 3.719 | 3.481 | 3.836 |
| 5.546 | 5.015 | 5.736 | 5.512 | 5.748 | 5.278 | 5.300 | 5.293 | 5.212 | 5.329 | 5.847 | 5.613 | 5.588 | 5.761 | 5.441 | 5.848 | 5.451 | 6.013 |
| 4.480 | 4.296 | 4.545 | 4.560 | 4.740 | 4.341 | 4.640 | 4.418 | 4.681 | 4.798 | 4.601 | 4.750 | 4.696 | 4.630 | 4.388 | 4.847 | 4.403 | 4.722 |
| 7.495 | 7.074 | 7.427 | 7.432 | 7.473 | 7.508 | 7.455 | 7.467 | 7.415 | 7.373 | 7.361 | 7.563 | 7.694 | 7.246 | 7.255 | 7.715 | 7.186 | 7.852 |
| 7.905 | 7.743 | 7.564 | 7.416 | 7.758 | 6.963 | 7.837 | 7.685 | 7.886 | 7.986 | 7.530 | 8.023 | 8.034 | 7.124 | 7.403 | 8.038 | 7.994 | 8.238 |
| 6.689 | 5.620 | 6.228 | 5.254 | 6.246 | 5.523 | 6.277 | 5.159 | 6.341 | 5.960 | 5.958 | 5.824 | 6.341 | 5.898 | 5.145 | 6.143 | 6.029 | 6.672 |
| 8.672 | 8.514 | 8.525 | 8.583 | 8.538 | 8.393 | 9.022 | 8.399 | 8.842 | 8.816 | 8.493 | 8.753 | 8.906 | 8.522 | 8.461 | 8.838 | 8.636 | 8.763 |
| 5.619 | 5.652 | 5.438 | 5.535 | 5.254 | 4.710 | 5.706 | 5.044 | 5.411 | 5.050 | 5.192 | 5.084 | 5.941 | 5.011 | 5.106 | 5.681 | 5.228 | 5.727 |
| 6.217 | 5.914 | 6.408 | 6.073 | 6.227 | 5.889 | 6.062 | 6.109 | 6.148 | 5.991 | 6.383 | 6.363 | 6.631 | 6.175 | 6.052 | 6.366 | 6.295 | 6.623 |
| 4.774 | 4.749 | 4.522 | 4.702 | 4.115 | 4.113 | 5.189 | 4.502 | 4.535 | 4.258 | 4.502 | 4.330 | 4.690 | 4.120 | 4.319 | 4.248 | 4.884 | 4.486 |
| 5.231 | 5.559 | 5.839 | 5.558 | 5.926 | 6.123 | 6.352 | 5.796 | 6.312 | 6.468 | 5.973 | 6.401 | 6.342 | 6.417 | 5.958 | 6.436 | 5.636 | 6.506 |
| 8.430 | 8.177 | 8.297 | 8.427 | 7.891 | 7.765 | 8.515 | 8.064 | 8.152 | 8.097 | 8.301 | 8.047 | 8.345 | 7.896 | 8.093 | 8.161 | 8.477 | 8.302 |
| 6.322 | 5.838 | 6.103 | 5.921 | 6.351 | 5.066 | 6.375 | 5.761 | 6.058 | 6.280 | 6.258 | 6.457 | 6.623 | 5.775 | 5.763 | 6.404 | 6.111 | 6.927 |
| 4.642 | 5.381 | 5.112 | 4.821 | 5.858 | 4.943 | 6.540 | 5.342 | 5.408 | 6.205 | 5.072 | 5.829 | 6.014 | 5.425 | 5.120 | 6.145 | 4.722 | 6.204 |
| 4.803 | 4.533 | 5.058 | 5.074 | 4.891 | 5.184 | 4.831 | 4.698 | 4.939 | 4.933 | 5.112 | 5.013 | 4.443 | 5.057 | 4.812 | 4.940 | 5.026 | 4.765 |
| 7.605 | 7.187 | 7.293 | 7.407 | 6.954 | 6.905 | 7.227 | 7.538 | 7.422 | 6.503 | 7.421 | 7.018 | 7.523 | 7.157 | 7.329 | 7.474 | 7.818 | 7.229 |
| 4.640 | 4.724 | 4.911 | 4.783 | 5.766 | 4.636 | 5.048 | 4.979 | 4.826 | 5.063 | 5.181 | 5.389 | 5.217 | 5.100 | 4.751 | 5.748 | 4.565 | 5.925 |
| 5.994 | 5.855 | 5.702 | 5.843 | 5.510 | 5.202 | 6.752 | 5.579 | 5.970 | 5.648 | 5.589 | 5.640 | 6.236 | 5.419 | 5.515 | 5.783 | 5.738 | 5.763 |
| 7.458 | 7.309 | 7.170 | 7.203 | 6.680 | 6.507 | 6.843 | 7.040 | 6.877 | 7.000 | 7.158 | 7.191 | 6.806 | 6.303 | 6.851 | 6.441 | 7.636 | 7.278 |
| 5.383 | 5.481 | 4.730 | 5.218 | 5.113 | 4.657 | 5.195 | 5.323 | 5.365 | 4.279 | 5.035 | 4.489 | 5.500 | 4.968 | 5.289 | 5.556 | 5.422 | 5.029 |
| 5.799 | 5.754 | 5.440 | 5.626 | 5.330 | 5.116 | 6.540 | 5.341 | 5.765 | 5.264 | 5.375 | 5.303 | 6.060 | 5.326 | 5.242 | 5.712 | 5.566 | 5.600 |
| 7.333 | 7.596 | 8.146 | 7.575 | 7.976 | 8.393 | 8.403 | 8.003 | 8.466 | 8.490 | 8.394 | 8.224 | 8.383 | 8.330 | 8.351 | 8.033 | 8.292 | 7.855 |
| 8.069 | 7.804 | 8.143 | 8.174 | 7.871 | 7.919 | 7.986 | 7.810 | 8.060 | 8.248 | 8.220 | 8.186 | 8.054 | 8.011 | 8.040 | 8.196 | 8.246 | 8.221 |
| 4.420 | 4.595 | 4.548 | 4.760 | 5.312 | 5.152 | 4.850 | 4.518 | 4.822 | 4.967 | 4.896 | 4.850 | 5.095 | 5.552 | 5.168 | 5.714 | 4.654 | 5.197 |
| 5.247 | 5.014 | 5.642 | 5.479 | 5.392 | 5.581 | 5.407 | 5.195 | 5.542 | 5.620 | 5.474 | 5.610 | 5.441 | 5.636 | 5.301 | 5.572 | 5.318 | 5.331 |
| 5.196 | 4.945 | 5.323 | 5.271 | 5.030 | 5.032 | 5.052 | 4.697 | 5.184 | 5.215 | 5.324 | 5.277 | 5.298 | 4.984 | 4.721 | 4.782 | 5.211 | 5.120 |
| 4.428 | 4.388 | 4.947 | 4.872 | 4.843 | 5.026 | 4.762 | 4.680 | 4.838 | 5.040 | 4.886 | 4.992 | 5.041 | 5.215 | 4.710 | 5.015 | 4.377 | 4.711 |
| 5.716 | 6.079 | 4.958 | 5.282 | 4.676 | 4.055 | 5.162 | 5.797 | 5.068 | 4.622 | 4.822 | 4.780 | 5.229 | 4.046 | 4.560 | 4.589 | 5.742 | 5.292 |
| 4.552 | 4.227 | 5.006 | 4.434 | 5.032 | 5.055 | 4.463 | 4.301 | 5.086 | 5.232 | 5.456 | 4.846 | 4.912 | 5.720 | 4.611 | 5.242 | 5.179 | 5.061 |
| 5.755 | 5.623 | 5.879 | 5.841 | 5.426 | 5.555 | 6.083 | 5.471 | 5.836 | 5.634 | 5.869 | 5.668 | 5.995 | 5.726 | 5.653 | 5.531 | 5.704 | 5.495 |
| 7.701 | 7.441 | 7.776 | 7.742 | 7.337 | 7.073 | 7.807 | 7.273 | 7.673 | 7.537 | 7.695 | 7.555 | 7.745 | 7.531 | 7.543 | 7.758 | 7.506 | 7.738 |
| 7.375 | 7.013 | 7.278 | 7.323 | 6.875 | 6.602 | 7.261 | 6.999 | 7.231 | 7.130 | 7.227 | 7.045 | 7.334 | 6.730 | 7.009 | 7.121 | 7.181 | 7.209 |
| 7.704 | 7.382 | 7.733 | 7.716 | 7.474 | 7.511 | 7.752 | 7.377 | 7.883 | 7.809 | 7.889 | 7.681 | 7.828 | 7.561 | 7.647 | 7.402 | 8.109 | 7.754 |
| 6.324 | 6.329 | 6.664 | 6.617 | 5.921 | 5.807 | 6.609 | 5.851 | 6.337 | 6.043 | 6.386 | 6.045 | 6.580 | 6.072 | 6.158 | 6.257 | 6.200 | 6.182 |
| 8.132 | 7.902 | 7.889 | 8.022 | 7.677 | 7.671 | 8.384 | 7.700 | 8.084 | 8.056 | 8.001 | 7.974 | 8.073 | 7.710 | 7.755 | 7.761 | 8.280 | 7.959 |
| 5.375 | 5.682 | 5.043 | 5.417 | 5.324 | 5.131 | 7.185 | 5.393 | 5.460 | 5.181 | 5.173 | 5.066 | 6.047 | 5.045 | 5.313 | 5.901 | 5.497 | 5.498 |
| 7.125 | 6.617 | 7.794 | 7.549 | 7.129 | 7.673 | 6.944 | 6.763 | 7.215 | 7.184 | 7.742 | 7.326 | 7.323 | 7.663 | 7.393 | 7.278 | 7.334 | 7.170 |
| 5.624 | 5.485 | 5.737 | 5.653 | 5.084 | 5.240 | 5.588 | 5.089 | 5.311 | 5.097 | 5.539 | 5.174 | 5.682 | 5.312 | 5.214 | 5.429 | 5.415 | 5.424 |
| 7.700 | 7.460 | 8.247 | 8.195 | 7.835 | 8.499 | 8.114 | 7.572 | 8.217 | 8.091 | 8.263 | 8.143 | 8.172 | 8.367 | 8.061 | 8.182 | 7.827 | 7.968 |
| 7.860 | 7.934 | 8.011 | 7.920 | 7.648 | 7.973 | 7.988 | 8.278 | 8.059 | 7.982 | 8.100 | 7.994 | 7.888 | 7.924 | 8.172 | 7.598 | 8.214 | 8.067 |
| 5.238 | 5.646 | 4.845 | 5.552 | 4.289 | 4.633 | 5.525 | 5.421 | 5.161 | 4.223 | 4.925 | 4.414 | 5.139 | 4.417 | 5.212 | 4.499 | 5.076 | 4.488 |
| 4.878 | 4.816 | 4.968 | 4.953 | 4.501 | 4.539 | 4.440 | 4.485 | 4.551 | 4.662 | 5.042 | 4.680 | 4.370 | 4.596 | 4.855 | 4.498 | 5.343 | 4.911 |
| 6.589 | 6.983 | 6.774 | 6.805 | 6.204 | 7.086 | 6.922 | 7.320 | 6.873 | 6.687 | 6.890 | 6.847 | 6.633 | 6.948 | 6.958 | 6.272 | 7.220 | 6.585 |
| 5.604 | 5.598 | 5.459 | 5.725 | 4.357 | 4.836 | 5.891 | 5.467 | 5.650 | 4.291 | 5.521 | 4.772 | 5.623 | 5.038 | 5.298 | 5.027 | 5.808 | 4.952 |
| 8.947 | 8.630 | 9.083 | 8.998 | 8.960 | 8.753 | 8.972 | 8.752 | 9.072 | 9.240 | 9.119 | 9.198 | 9.098 | 8.975 | 8.947 | 9.201 | 9.195 | 9.215 |
| 6.418 | 6.849 | 7.089 | 6.795 | 6.736 | 7.473 | 6.278 | 7.474 | 6.691 | 7.044 | 7.128 | 7.304 | 6.677 | 7.277 | 7.119 | 6.293 | 7.055 | 6.662 |
| 5.116 | 4.933 | 5.786 | 5.558 | 5.337 | 5.428 | 5.568 | 5.333 | 5.847 | 5.704 | 5.812 | 5.652 | 5.850 | 5.797 | 5.581 | 5.649 | 5.389 | 5.556 |
| 3.408 | 3.764 | 4.194 | 3.994 | 3.561 | 3.484 | 4.472 | 5.462 | 4.676 | 4.089 | 3.631 | 3.967 | 5.045 | 4.144 | 4.290 | 3.443 | 3.596 | 3.959 |
| 5.561 | 5.738 | 5.315 | 5.608 | 5.791 | 5.813 | 5.764 | 5.577 | 5.631 | 5.581 | 5.559 | 5.682 | 5.785 | 5.669 | 5.813 | 5.868 | 5.730 | 6.049 |
| 4.516 | 4.596 | 4.903 | 4.728 | 4.483 | 5.049 | 4.853 | 5.010 | 5.087 | 4.634 | 4.970 | 5.005 | 4.735 | 4.668 | 4.679 | 4.372 | 5.102 | 4.761 |
| 7.073 | 6.978 | 6.934 | 6.963 | 6.750 | 6.160 | 6.989 | 7.216 | 6.924 | 6.744 | 6.900 | 6.899 | 7.045 | 6.401 | 6.666 | 7.077 | 7.252 | 7.286 |
| 6.832 | 7.399 | 7.429 | 7.425 | 6.955 | 6.998 | 7.705 | 8.523 | 7.831 | 7.404 | 7.226 | 7.636 | 7.848 | 7.418 | 7.796 | 6.252 | 7.341 | 7.356 |
| 5.540 | 5.357 | 5.607 | 5.692 | 6.394 | 5.870 | 6.300 | 5.597 | 6.110 | 6.096 | 5.719 | 6.026 | 6.375 | 6.413 | 6.194 | 6.908 | 5.797 | 6.394 |
| 4.822 | 4.997 | 4.870 | 5.076 | 6.242 | 5.483 | 5.939 | 5.034 | 5.866 | 6.119 | 5.120 | 6.210 | 5.774 | 5.231 | 4.759 | 6.124 | 4.252 | 5.934 |
| 7.795 | 8.317 | 8.211 | 8.134 | 8.138 | 8.608 | 8.776 | 8.881 | 8.767 | 8.694 | 8.354 | 8.824 | 8.664 | 8.141 | 8.571 | 7.717 | 8.465 | 8.100 |
| 7.376 | 7.365 | 7.856 | 7.614 | 7.671 | 8.478 | 7.479 | 8.250 | 7.671 | 8.303 | 8.168 | 8.259 | 7.400 | 8.273 | 8.191 | 7.573 | 8.333 | 7.786 |
| 4.943 | 5.076 | 5.276 | 5.221 | 4.750 | 5.075 | 5.149 | 5.335 | 5.083 | 4.768 | 5.128 | 4.885 | 5.068 | 5.129 | 5.053 | 4.848 | 5.296 | 4.868 |
| 8.224 | 7.778 | 8.246 | 7.691 | 7.849 | 8.161 | 7.475 | 7.690 | 7.980 | 8.313 | 8.468 | 8.021 | 7.597 | 8.320 | 7.780 | 7.851 | 8.784 | 8.113 |
| 6.238 | 6.962 | 6.390 | 6.717 | 7.042 | 7.113 | 7.177 | 7.380 | 7.013 | 7.034 | 6.682 | 7.179 | 7.217 | 6.983 | 7.320 | 7.053 | 6.996 | 7.162 |
| 6.374 | 6.187 | 6.480 | 6.245 | 5.952 | 6.067 | 6.615 | 6.122 | 6.685 | 6.058 | 6.547 | 6.124 | 6.727 | 5.882 | 6.248 | 5.780 | 6.744 | 6.399 |
| 5.567 | 5.774 | 5.629 | 5.719 | 6.195 | 6.446 | 5.973 | 6.201 | 6.035 | 6.284 | 6.091 | 6.369 | 6.044 | 6.464 | 6.488 | 6.470 | 6.335 | 6.287 |
| 6.102 | 6.438 | 5.795 | 5.934 | 6.410 | 6.658 | 5.725 | 6.532 | 6.127 | 6.258 | 6.189 | 6.302 | 5.822 | 6.575 | 7.165 | 6.651 | 6.599 | 6.705 |
| 4.908 | 4.944 | 5.101 | 5.116 | 5.138 | 4.842 | 5.127 | 5.219 | 5.307 | 5.236 | 5.322 | 5.157 | 5.264 | 5.062 | 5.083 | 5.069 | 5.338 | 5.249 |
| 5.996 | 5.736 | 5.061 | 5.141 | 5.673 | 5.667 | 4.886 | 5.084 | 5.291 | 5.685 | 5.334 | 5.378 | 4.904 | 6.263 | 5.875 | 6.230 | 5.764 | 6.127 |
| 8.184 | 7.833 | 8.104 | 8.084 | 7.772 | 7.807 | 7.880 | 7.783 | 7.685 | 7.943 | 8.206 | 7.913 | 7.660 | 7.982 | 7.987 | 8.064 | 8.383 | 8.147 |
| 5.487 | 6.094 | 6.095 | 5.671 | 6.608 | 6.524 | 6.443 | 6.662 | 5.966 | 7.029 | 6.308 | 6.926 | 6.237 | 6.579 | 6.462 | 6.698 | 6.093 | 6.844 |
| 8.039 | 7.349 | 7.958 | 7.065 | 7.771 | 7.480 | 6.727 | 6.772 | 7.832 | 8.066 | 8.279 | 7.323 | 7.129 | 7.854 | 6.769 | 7.451 | 8.567 | 7.790 |
| 6.071 | 5.931 | 5.918 | 5.897 | 5.369 | 4.893 | 5.865 | 6.002 | 5.768 | 5.642 | 5.877 | 5.855 | 5.841 | 4.962 | 5.505 | 5.174 | 6.500 | 5.945 |
| 7.388 | 7.496 | 7.571 | 7.571 | 7.332 | 7.606 | 7.827 | 7.670 | 8.078 | 7.943 | 7.697 | 7.849 | 7.761 | 7.399 | 7.576 | 7.135 | 8.099 | 7.452 |
| 8.122 | 8.076 | 8.281 | 8.156 | 8.117 | 8.530 | 8.060 | 8.552 | 8.350 | 8.528 | 8.481 | 8.519 | 7.986 | 8.320 | 8.382 | 7.862 | 8.858 | 8.247 |
| 7.246 | 7.783 | 7.524 | 7.333 | 7.656 | 7.825 | 7.703 | 8.485 | 7.813 | 8.158 | 7.806 | 8.111 | 7.662 | 7.849 | 7.708 | 7.079 | 8.017 | 7.510 |
| 8.032 | 8.230 | 7.880 | 7.888 | 7.586 | 7.673 | 8.203 | 8.376 | 8.133 | 7.638 | 7.988 | 7.800 | 8.220 | 7.536 | 7.909 | 7.359 | 8.531 | 7.918 |
| 5.089 | 5.600 | 4.790 | 5.187 | 4.653 | 4.558 | 4.763 | 5.789 | 4.877 | 4.579 | 4.841 | 4.807 | 4.768 | 4.684 | 4.711 | 4.962 | 5.469 | 4.868 |
| 6.853 | 7.099 | 6.960 | 6.905 | 6.143 | 7.030 | 6.328 | 7.464 | 6.621 | 6.165 | 6.830 | 6.516 | 6.533 | 6.309 | 6.717 | 5.783 | 7.253 | 6.538 |
| 7.493 | 7.248 | 8.064 | 7.679 | 7.756 | 8.197 | 7.655 | 7.788 | 7.846 | 8.187 | 8.195 | 8.161 | 7.707 | 8.119 | 8.105 | 7.950 | 8.315 | 7.978 |
| 5.018 | 5.297 | 5.552 | 5.308 | 6.259 | 6.483 | 5.884 | 5.952 | 5.967 | 6.479 | 5.954 | 6.211 | 5.768 | 6.498 | 6.273 | 6.169 | 5.958 | 6.128 |
| 4.302 | 4.696 | 4.148 | 4.268 | 3.858 | 4.348 | 3.940 | 5.337 | 4.173 | 3.759 | 4.223 | 4.054 | 3.961 | 4.034 | 4.188 | 3.524 | 4.931 | 3.881 |
| 5.105 | 5.458 | 5.873 | 5.360 | 6.439 | 6.558 | 6.252 | 5.957 | 6.257 | 6.841 | 6.155 | 6.582 | 6.268 | 6.672 | 6.148 | 6.698 | 5.908 | 6.596 |
| 6.094 | 6.377 | 5.413 | 5.753 | 4.810 | 4.910 | 5.245 | 6.882 | 5.512 | 5.190 | 5.362 | 5.503 | 5.371 | 4.521 | 5.186 | 4.284 | 6.463 | 5.422 |
| 5.089 | 4.863 | 5.115 | 5.189 | 4.997 | 5.154 | 5.143 | 5.042 | 5.275 | 5.233 | 5.314 | 5.173 | 5.093 | 5.207 | 5.074 | 5.019 | 5.565 | 5.172 |
| 5.898 | 6.042 | 6.307 | 6.097 | 4.699 | 6.179 | 5.708 | 6.664 | 6.281 | 5.503 | 6.246 | 5.955 | 5.606 | 5.888 | 6.248 | 5.149 | 6.569 | 5.915 |
| 4.931 | 5.407 | 5.194 | 5.234 | 5.284 | 5.252 | 5.240 | 5.908 | 5.404 | 5.284 | 5.233 | 5.379 | 5.218 | 5.245 | 5.427 | 5.577 | 5.446 | 5.569 |
| 6.227 | 6.488 | 6.130 | 6.288 | 5.437 | 5.608 | 5.807 | 6.658 | 6.037 | 5.515 | 6.075 | 5.706 | 5.841 | 5.520 | 6.011 | 5.712 | 6.596 | 6.048 |
| 6.074 | 6.230 | 5.710 | 5.809 | 5.606 | 5.681 | 5.699 | 6.587 | 5.791 | 5.523 | 5.757 | 5.676 | 5.509 | 5.174 | 5.695 | 4.874 | 6.601 | 5.690 |
| 7.568 | 7.208 | 7.862 | 7.661 | 7.286 | 7.889 | 7.105 | 7.906 | 7.470 | 7.709 | 7.904 | 7.902 | 7.216 | 7.816 | 7.891 | 7.427 | 8.278 | 7.826 |
| 5.255 | 5.842 | 6.130 | 5.774 | 5.375 | 6.528 | 5.087 | 6.591 | 5.862 | 5.867 | 6.002 | 6.350 | 5.383 | 6.215 | 5.877 | 5.205 | 6.270 | 5.648 |
| 6.378 | 6.531 | 6.598 | 6.298 | 5.670 | 7.045 | 6.240 | 7.473 | 6.359 | 6.584 | 6.453 | 6.564 | 5.957 | 6.153 | 6.499 | 6.152 | 7.411 | 6.223 |
| 5.571 | 5.616 | 5.595 | 5.648 | 5.895 | 5.887 | 5.677 | 6.012 | 5.904 | 5.741 | 5.758 | 5.878 | 5.800 | 6.042 | 6.140 | 6.269 | 6.177 | 6.279 |
| 5.731 | 6.189 | 5.839 | 5.786 | 6.127 | 6.178 | 6.107 | 6.975 | 6.152 | 6.251 | 6.065 | 6.314 | 6.056 | 5.945 | 6.053 | 6.110 | 6.765 | 6.162 |
| 4.840 | 5.124 | 4.571 | 4.772 | 4.184 | 4.525 | 4.356 | 5.544 | 4.742 | 4.097 | 4.624 | 4.256 | 4.484 | 4.218 | 4.710 | 4.318 | 5.353 | 4.291 |
| 4.503 | 4.701 | 4.504 | 4.451 | 5.025 | 5.323 | 4.712 | 5.115 | 4.663 | 5.024 | 4.662 | 5.049 | 4.651 | 5.384 | 5.313 | 5.407 | 5.008 | 5.327 |
| 4.620 | 5.050 | 4.294 | 4.404 | 4.451 | 4.332 | 4.362 | 5.583 | 4.281 | 4.387 | 4.316 | 4.624 | 4.451 | 4.133 | 4.448 | 4.168 | 4.992 | 4.810 |
| 3.383 | 3.916 | 3.936 | 3.708 | 4.256 | 4.630 | 4.264 | 4.604 | 4.276 | 4.990 | 4.149 | 4.742 | 4.137 | 4.567 | 4.326 | 4.252 | 4.117 | 4.420 |
| 4.241 | 4.410 | 5.073 | 4.622 | 4.632 | 5.550 | 4.583 | 5.679 | 5.118 | 5.297 | 5.206 | 5.647 | 4.469 | 5.594 | 5.289 | 4.932 | 5.209 | 4.999 |
| 4.713 | 4.815 | 5.154 | 4.854 | 4.798 | 5.538 | 4.614 | 5.769 | 4.922 | 4.950 | 5.155 | 5.299 | 4.742 | 5.124 | 5.000 | 4.639 | 5.680 | 4.844 |
| 4.914 | 5.103 | 5.574 | 4.978 | 5.034 | 6.355 | 5.014 | 6.210 | 5.217 | 5.980 | 5.659 | 5.987 | 4.708 | 5.970 | 5.780 | 5.246 | 6.056 | 5.290 |
| 4.491 | 4.748 | 4.938 | 4.588 | 4.347 | 5.118 | 4.480 | 5.322 | 4.823 | 4.998 | 4.968 | 5.102 | 4.476 | 4.871 | 4.905 | 4.536 | 5.321 | 4.787 |
| 5.259 | 5.738 | 5.456 | 5.379 | 5.186 | 5.729 | 5.259 | 6.941 | 5.429 | 5.766 | 5.575 | 5.953 | 5.133 | 5.319 | 5.634 | 4.574 | 6.246 | 5.521 |
| 5.195 | 5.383 | 5.929 | 5.530 | 5.084 | 6.259 | 4.815 | 6.458 | 5.343 | 5.292 | 5.843 | 5.792 | 4.751 | 5.649 | 5.688 | 4.571 | 6.161 | 5.193 |

### Supplemental Table 2 Continued.

| 43080A | 43049A | 45008A | 45011A | 44067A | 45040A | 44062A | 43099A | 46115A | 44652D | 45015A | 43120A | 43157A | 45005A | 43144A | 43117A |
|---|---|---|---|---|---|---|---|---|---|---|---|---|---|---|---|
| Non-progre | Non-progre | Non-progre | Non-progre | Non-progre | Non-progre | Non-progre | Non-progre | Non-progre | Non-progre | Non-progre | Non-progre | Non-progre | Non-progre | Non-progre | Non-progre |
| NP | NP | NP | NP | NP | NP | NP | NP | NP | NP | NP | NP | NP | NP | NP | NP |
| 3.643 | 3.617 | 3.738 | 3.812 | 3.967 | 3.699 | 3.935 | 3.906 | 4.025 | 3.850 | 3.954 | 3.784 | 3.828 | 4.178 | 3.925 | 4.040 |
| 5.464 | 5.766 | 5.709 | 5.607 | 5.887 | 5.450 | 5.701 | 5.916 | 5.423 | 5.692 | 5.723 | 5.769 | 5.523 | 5.774 | 5.375 | 5.440 |
| 4.658 | 4.617 | 4.694 | 4.755 | 4.626 | 4.524 | 4.555 | 4.560 | 4.359 | 4.425 | 4.445 | 4.496 | 4.475 | 4.726 | 4.284 | 4.754 |
| 7.813 | 7.806 | 7.653 | 7.837 | 7.974 | 7.717 | 7.751 | 7.725 | 6.921 | 7.246 | 7.748 | 7.562 | 7.573 | 8.072 | 7.521 | 6.896 |
| 7.771 | 8.302 | 8.391 | 8.271 | 8.240 | 8.054 | 7.855 | 8.089 | 7.981 | 8.228 | 8.235 | 8.445 | 8.197 | 8.964 | 8.712 | 8.729 |
| 6.119 | 5.290 | 5.420 | 6.348 | 6.623 | 5.515 | 6.852 | 6.451 | 6.354 | 6.309 | 6.439 | 6.267 | 6.439 | 6.402 | 6.495 | 4.921 |
| 8.440 | 8.786 | 8.708 | 8.801 | 8.961 | 8.465 | 8.647 | 8.639 | 8.620 | 8.793 | 8.814 | 8.938 | 8.614 | 9.113 | 8.983 | 8.842 |
| 4.743 | 4.857 | 5.022 | 5.144 | 5.669 | 5.452 | 5.200 | 5.129 | 4.807 | 5.223 | 5.194 | 5.444 | 4.726 | 6.107 | 5.835 | 5.161 |
| 6.217 | 6.259 | 6.803 | 6.318 | 6.531 | 6.381 | 6.642 | 6.500 | 6.311 | 6.390 | 6.362 | 6.533 | 6.212 | 6.872 | 6.398 | 6.207 |
| 4.241 | 4.575 | 4.666 | 4.485 | 4.600 | 4.932 | 4.315 | 4.323 | 4.510 | 4.612 | 4.495 | 4.804 | 4.577 | 5.366 | 5.260 | 5.014 |
| 6.641 | 6.716 | 6.719 | 6.501 | 6.519 | 5.436 | 6.736 | 6.556 | 6.613 | 6.334 | 6.539 | 6.510 | 6.433 | 6.345 | 6.091 | 6.276 |
| 7.819 | 7.966 | 8.260 | 8.181 | 8.399 | 8.421 | 7.863 | 7.896 | 7.867 | 8.220 | 8.114 | 8.302 | 7.871 | 8.831 | 8.639 | 8.246 |
| 6.287 | 6.563 | 6.759 | 6.515 | 6.607 | 6.232 | 6.541 | 6.459 | 6.073 | 6.248 | 6.236 | 6.494 | 6.086 | 7.128 | 6.634 | 6.220 |
| 6.078 | 6.295 | 6.482 | 6.106 | 5.734 | 5.109 | 6.124 | 5.869 | 5.718 | 5.941 | 5.860 | 5.803 | 5.761 | 5.958 | 5.351 | 5.482 |
| 5.195 | 4.910 | 5.215 | 5.040 | 5.150 | 5.247 | 5.314 | 5.301 | 5.303 | 5.328 | 5.292 | 5.335 | 5.214 | 5.254 | 5.296 | 5.435 |
| 6.738 | 7.208 | 6.954 | 7.249 | 7.659 | 7.964 | 7.148 | 7.470 | 7.315 | 7.223 | 7.543 | 7.636 | 7.627 | 8.153 | 8.444 | 8.288 |
| 5.378 | 5.348 | 5.776 | 5.431 | 5.383 | 4.569 | 5.505 | 5.496 | 5.287 | 5.177 | 5.272 | 5.372 | 5.340 | 5.550 | 4.660 | 4.969 |
| 5.277 | 5.552 | 5.657 | 5.690 | 5.994 | 5.665 | 5.493 | 5.418 | 5.296 | 5.559 | 5.564 | 5.832 | 5.349 | 6.371 | 6.210 | 5.784 |
| 7.046 | 7.205 | 7.386 | 7.531 | 7.187 | 7.842 | 6.593 | 6.927 | 7.418 | 7.761 | 7.203 | 7.567 | 7.587 | 8.503 | 8.730 | 7.922 |
| 4.400 | 4.563 | 4.358 | 4.754 | 5.359 | 5.564 | 4.827 | 5.083 | 4.805 | 4.872 | 5.366 | 5.301 | 5.243 | 5.639 | 6.319 | 5.966 |
| 5.165 | 5.294 | 5.440 | 5.451 | 5.923 | 5.694 | 5.599 | 5.356 | 5.097 | 5.364 | 5.491 | 5.696 | 5.334 | 6.271 | 6.181 | 5.630 |
| 8.796 | 8.864 | 8.604 | 8.624 | 8.028 | 8.651 | 8.885 | 8.779 | 8.981 | 8.480 | 8.811 | 8.533 | 9.293 | 8.202 | 8.318 | 9.403 |
| 8.123 | 8.230 | 8.211 | 8.316 | 8.410 | 8.069 | 8.073 | 8.119 | 8.176 | 8.372 | 8.273 | 8.399 | 8.212 | 8.697 | 8.547 | 8.402 |
| 5.387 | 5.027 | 4.760 | 4.992 | 5.265 | 4.710 | 5.582 | 5.611 | 5.159 | 5.252 | 5.219 | 5.357 | 5.243 | 4.918 | 5.104 | 5.230 |
| 5.491 | 5.597 | 5.474 | 5.700 | 5.615 | 5.230 | 5.656 | 5.584 | 5.641 | 5.617 | 5.730 | 5.742 | 5.585 | 5.616 | 5.822 | 5.730 |
| 4.963 | 4.948 | 5.200 | 5.110 | 5.290 | 5.203 | 5.121 | 5.021 | 5.085 | 5.378 | 5.119 | 5.198 | 5.100 | 5.358 | 5.199 | 5.122 |
| 4.749 | 4.815 | 4.679 | 4.954 | 5.008 | 4.599 | 4.894 | 4.935 | 4.873 | 4.895 | 4.869 | 4.818 | 4.772 | 4.788 | 4.722 | 4.780 |
| 4.364 | 4.962 | 5.574 | 5.067 | 5.184 | 6.474 | 4.553 | 4.829 | 4.593 | 5.080 | 4.843 | 5.308 | 4.706 | 7.356 | 7.097 | 5.952 |
| 5.679 | 4.672 | 4.822 | 4.939 | 5.097 | 4.835 | 6.310 | 5.823 | 6.239 | 5.315 | 5.540 | 5.400 | 6.139 | 4.868 | 5.117 | 5.402 |
| 5.144 | 5.591 | 5.398 | 5.609 | 5.903 | 5.424 | 5.404 | 5.397 | 5.485 | 5.493 | 5.397 | 5.643 | 5.277 | 5.822 | 5.624 | 5.475 |
| 6.992 | 7.301 | 7.493 | 7.432 | 8.008 | 7.261 | 7.381 | 7.306 | 7.175 | 7.443 | 7.351 | 7.631 | 7.206 | 7.825 | 7.710 | 7.474 |
| 6.389 | 6.864 | 7.078 | 7.012 | 7.427 | 6.850 | 6.633 | 6.768 | 6.635 | 7.027 | 6.792 | 7.138 | 6.427 | 7.458 | 7.127 | 6.584 |
| 7.716 | 7.900 | 8.049 | 7.934 | 7.815 | 7.855 | 7.639 | 7.703 | 7.874 | 8.048 | 7.858 | 7.985 | 7.892 | 8.430 | 8.020 | 8.064 |
| 5.499 | 5.966 | 5.998 | 6.016 | 6.548 | 6.068 | 5.975 | 5.883 | 5.845 | 5.984 | 6.024 | 6.259 | 5.647 | 6.731 | 6.663 | 6.021 |
| 7.646 | 8.124 | 8.157 | 8.121 | 8.214 | 7.976 | 7.722 | 7.765 | 7.896 | 8.197 | 7.977 | 8.228 | 7.802 | 8.678 | 8.508 | 7.996 |
| 5.082 | 5.283 | 5.307 | 5.215 | 5.695 | 5.984 | 5.555 | 5.500 | 5.393 | 5.535 | 5.653 | 5.805 | 5.626 | 6.641 | 7.135 | 6.073 |
| 7.432 | 7.221 | 7.275 | 7.350 | 7.359 | 7.124 | 7.623 | 7.506 | 7.680 | 7.688 | 7.423 | 7.433 | 7.371 | 6.915 | 6.743 | 7.020 |
| 5.148 | 5.023 | 5.263 | 5.219 | 5.587 | 5.634 | 5.436 | 5.420 | 5.113 | 5.381 | 5.435 | 5.648 | 5.297 | 5.886 | 5.730 | 5.485 |
| 8.268 | 8.124 | 7.970 | 8.067 | 8.271 | 8.042 | 8.234 | 8.303 | 8.215 | 8.353 | 8.318 | 8.286 | 8.204 | 8.161 | 8.188 | 8.119 |
| 8.342 | 8.469 | 8.427 | 8.437 | 8.149 | 8.818 | 8.163 | 8.375 | 8.648 | 8.540 | 8.583 | 8.582 | 8.971 | 9.047 | 9.402 | 9.362 |
| 3.909 | 4.636 | 4.515 | 4.532 | 5.324 | 5.325 | 4.176 | 4.325 | 4.106 | 4.093 | 4.401 | 4.751 | 4.349 | 5.870 | 5.853 | 5.189 |
| 4.561 | 4.972 | 5.172 | 4.995 | 5.002 | 5.320 | 4.738 | 4.824 | 5.129 | 5.224 | 5.024 | 5.213 | 5.026 | 5.578 | 5.655 | 5.470 |
| 7.140 | 7.169 | 7.293 | 7.121 | 6.782 | 7.689 | 7.227 | 7.141 | 7.404 | 7.351 | 7.412 | 7.269 | 7.651 | 7.403 | 8.092 | 8.095 |
| 4.281 | 4.752 | 4.622 | 4.827 | 5.749 | 6.026 | 5.045 | 4.959 | 5.075 | 4.793 | 5.346 | 5.401 | 5.237 | 6.578 | 6.821 | 6.042 |
| 9.123 | 9.334 | 9.288 | 9.369 | 9.394 | 9.019 | 9.144 | 9.157 | 9.233 | 9.395 | 9.243 | 9.425 | 9.249 | 9.628 | 9.485 | 9.478 |
| 7.628 | 7.449 | 7.485 | 7.592 | 7.041 | 7.965 | 7.740 | 7.757 | 8.158 | 7.866 | 7.788 | 7.491 | 8.392 | 7.830 | 8.125 | 8.559 |
| 5.525 | 5.690 | 5.663 | 5.565 | 5.830 | 5.403 | 5.562 | 5.912 | 5.707 | 5.648 | 6.032 | 5.896 | 5.669 | 5.839 | 5.762 | 5.907 |
| 4.516 | 4.866 | 4.811 | 4.069 | 3.886 | 4.996 | 3.948 | 4.549 | 4.107 | 3.884 | 5.258 | 4.556 | 4.712 | 5.403 | 5.054 | 6.582 |
| 5.711 | 5.765 | 5.764 | 5.737 | 5.892 | 5.919 | 5.725 | 5.978 | 5.790 | 5.932 | 6.010 | 6.151 | 6.073 | 6.058 | 6.390 | 6.186 |
| 4.574 | 5.601 | 5.374 | 5.070 | 4.985 | 5.296 | 5.124 | 5.378 | 5.441 | 5.015 | 5.232 | 5.531 | 5.549 | 5.800 | 6.046 | 5.835 |
| 6.639 | 7.214 | 7.282 | 7.070 | 7.437 | 7.645 | 6.973 | 6.994 | 6.833 | 7.174 | 7.183 | 7.429 | 7.044 | 8.324 | 8.154 | 7.756 |
| 7.953 | 8.248 | 8.266 | 7.888 | 7.267 | 8.671 | 7.603 | 7.875 | 7.755 | 7.688 | 8.416 | 8.051 | 8.574 | 8.909 | 8.915 | 9.760 |
| 6.072 | 6.055 | 5.991 | 6.061 | 6.276 | 5.590 | 6.238 | 6.288 | 5.847 | 6.036 | 6.054 | 6.098 | 5.990 | 5.927 | 6.216 | 5.890 |
| 6.250 | 5.947 | 6.613 | 5.846 | 5.547 | 4.317 | 6.667 | 5.748 | 5.809 | 5.225 | 5.411 | 5.292 | 5.523 | 5.630 | 4.395 | 4.810 |
| 8.863 | 9.390 | 9.164 | 9.150 | 8.313 | 9.115 | 8.553 | 8.844 | 8.751 | 8.730 | 9.138 | 8.933 | 9.226 | 9.418 | 9.207 | 10.202 |
| 8.989 | 8.643 | 8.666 | 8.745 | 8.006 | 8.686 | 8.731 | 8.726 | 9.285 | 8.929 | 8.938 | 8.586 | 9.542 | 8.447 | 8.776 | 9.681 |
| 4.928 | 5.026 | 5.177 | 5.114 | 5.417 | 5.538 | 5.173 | 5.154 | 5.381 | 5.160 | 5.461 | 5.391 | 5.491 | 5.810 | 6.058 | 5.816 |
| 8.664 | 8.088 | 8.308 | 8.540 | 8.209 | 8.296 | 9.034 | 8.779 | 9.535 | 9.185 | 8.935 | 8.783 | 9.472 | 8.632 | 9.348 | 8.913 |
| 7.680 | 7.656 | 7.585 | 7.442 | 7.109 | 7.732 | 7.134 | 7.611 | 7.035 | 7.337 | 7.688 | 7.619 | 7.817 | 7.632 | 7.891 | 8.439 |
| 6.204 | 6.379 | 6.747 | 6.417 | 6.233 | 6.737 | 6.127 | 6.266 | 6.498 | 6.444 | 6.421 | 6.569 | 6.582 | 7.510 | 6.889 | 6.886 |
| 7.167 | 6.579 | 6.580 | 6.614 | 6.268 | 6.710 | 6.985 | 7.078 | 7.119 | 7.039 | 7.140 | 6.891 | 7.576 | 6.316 | 7.037 | 7.735 |
| 6.868 | 6.613 | 6.639 | 6.323 | 6.700 | 6.470 | 7.298 | 7.266 | 7.202 | 7.263 | 7.308 | 7.385 | 7.383 | 6.565 | 7.817 | 8.092 |
| 5.369 | 5.144 | 5.459 | 5.326 | 5.279 | 5.580 | 5.316 | 5.372 | 5.473 | 5.415 | 5.534 | 5.515 | 5.341 | 5.661 | 5.655 | 5.782 |
| 6.141 | 5.472 | 5.487 | 5.779 | 6.304 | 5.239 | 7.054 | 6.884 | 6.639 | 6.728 | 6.862 | 6.936 | 6.619 | 5.764 | 7.426 | 6.603 |
| 8.041 | 8.021 | 8.192 | 8.071 | 8.351 | 8.195 | 8.142 | 8.093 | 8.177 | 8.379 | 8.244 | 8.418 | 8.139 | 8.534 | 8.694 | 8.157 |
| 7.506 | 7.376 | 7.916 | 7.219 | 6.572 | 6.348 | 7.405 | 7.050 | 7.444 | 7.116 | 6.792 | 6.740 | 7.574 | 6.800 | 5.939 | 7.097 |
| 8.380 | 7.215 | 7.581 | 8.257 | 7.768 | 7.473 | 8.927 | 8.539 | 9.598 | 8.942 | 8.730 | 8.433 | 9.367 | 8.083 | 8.887 | 8.214 |
| 5.195 | 6.286 | 6.357 | 6.193 | 6.147 | 6.706 | 5.398 | 5.525 | 5.773 | 6.143 | 5.980 | 6.254 | 5.916 | 7.659 | 7.205 | 6.631 |
| 7.975 | 8.339 | 8.156 | 8.241 | 7.633 | 8.065 | 7.872 | 7.918 | 8.365 | 8.314 | 8.266 | 8.153 | 8.448 | 8.703 | 8.567 | 8.826 |
| 8.982 | 9.003 | 9.002 | 8.967 | 8.439 | 9.042 | 8.787 | 8.844 | 9.325 | 9.239 | 9.185 | 9.036 | 9.564 | 9.358 | 9.654 | 9.851 |
| 8.366 | 8.506 | 8.354 | 8.560 | 7.834 | 8.630 | 8.420 | 8.350 | 8.879 | 8.403 | 8.605 | 8.288 | 9.137 | 8.783 | 8.907 | 9.618 |
| 7.628 | 8.222 | 8.559 | 8.179 | 7.975 | 8.845 | 7.831 | 7.921 | 8.116 | 8.202 | 8.123 | 8.264 | 8.162 | 9.336 | 9.103 | 8.963 |
| 4.765 | 5.030 | 5.340 | 5.029 | 5.071 | 6.229 | 5.012 | 4.772 | 5.278 | 5.258 | 5.164 | 5.273 | 5.562 | 6.702 | 6.722 | 6.273 |
| 6.765 | 6.790 | 7.212 | 6.934 | 6.482 | 8.092 | 6.648 | 6.934 | 7.390 | 7.359 | 7.153 | 7.085 | 7.520 | 8.176 | 8.239 | 7.930 |
| 8.762 | 8.596 | 8.498 | 8.500 | 8.102 | 8.238 | 8.615 | 8.532 | 8.897 | 8.626 | 8.651 | 8.519 | 9.150 | 8.099 | 8.088 | 8.951 |
| 7.264 | 6.748 | 6.557 | 6.612 | 6.129 | 6.153 | 7.250 | 7.053 | 7.188 | 6.831 | 7.019 | 6.698 | 7.439 | 6.016 | 6.528 | 7.224 |
| 4.290 | 4.458 | 4.434 | 4.423 | 4.054 | 5.976 | 4.189 | 4.380 | 4.860 | 4.632 | 4.783 | 4.799 | 5.464 | 5.906 | 6.489 | 6.548 |
| 7.424 | 7.250 | 7.305 | 6.885 | 6.369 | 5.952 | 7.629 | 7.077 | 7.280 | 6.740 | 6.966 | 6.757 | 7.392 | 6.360 | 5.923 | 6.857 |
| 5.448 | 5.920 | 6.186 | 5.954 | 5.251 | 7.441 | 4.634 | 5.321 | 5.845 | 6.146 | 5.859 | 6.108 | 6.517 | 7.956 | 8.141 | 7.455 |
| 5.543 | 5.361 | 5.423 | 5.372 | 5.386 | 5.520 | 5.304 | 5.351 | 5.483 | 5.564 | 5.462 | 5.506 | 5.614 | 5.833 | 5.734 | 5.792 |
| 6.019 | 6.258 | 6.489 | 6.091 | 6.124 | 6.776 | 5.876 | 5.847 | 6.671 | 6.481 | 6.436 | 6.344 | 6.759 | 7.504 | 7.776 | 7.390 |
| 5.643 | 5.815 | 5.703 | 5.778 | 5.550 | 6.120 | 5.518 | 5.705 | 5.850 | 5.975 | 5.967 | 5.827 | 6.165 | 6.532 | 6.683 | 6.509 |
| 5.861 | 6.097 | 6.158 | 5.932 | 6.237 | 7.024 | 5.732 | 5.889 | 6.149 | 6.461 | 6.098 | 6.402 | 6.365 | 7.735 | 7.791 | 6.968 |
| 6.074 | 6.115 | 6.453 | 6.078 | 5.632 | 7.148 | 5.712 | 5.859 | 6.455 | 6.590 | 6.246 | 6.318 | 6.823 | 7.304 | 7.592 | 7.270 |
| 8.257 | 8.259 | 8.308 | 8.137 | 8.083 | 8.226 | 8.080 | 8.130 | 8.503 | 8.347 | 8.396 | 8.364 | 8.713 | 8.553 | 8.760 | 8.699 |
| 6.594 | 6.655 | 6.401 | 6.756 | 6.089 | 6.947 | 6.777 | 6.779 | 7.252 | 6.667 | 7.042 | 6.483 | 7.480 | 7.348 | 7.611 | 7.676 |
| 7.531 | 7.040 | 7.082 | 7.174 | 6.162 | 7.981 | 7.442 | 7.107 | 7.753 | 7.598 | 7.531 | 7.197 | 8.250 | 7.166 | 8.102 | 8.589 |
| 6.420 | 6.265 | 6.198 | 6.092 | 6.396 | 6.265 | 6.224 | 6.409 | 6.352 | 6.415 | 6.461 | 6.582 | 6.676 | 6.539 | 7.018 | 6.779 |
| 6.894 | 6.913 | 7.010 | 6.785 | 6.094 | 7.446 | 6.594 | 6.717 | 7.016 | 6.987 | 6.961 | 6.809 | 7.567 | 7.461 | 7.816 | 8.069 |
| 4.485 | 4.627 | 4.561 | 4.684 | 4.471 | 6.140 | 4.482 | 4.596 | 5.118 | 4.927 | 4.921 | 4.973 | 5.526 | 6.024 | 6.580 | 6.237 |
| 6.056 | 5.567 | 5.600 | 5.357 | 5.388 | 5.321 | 6.064 | 5.960 | 5.771 | 5.841 | 5.980 | 6.005 | 6.364 | 5.535 | 6.102 | 6.366 |
| 4.950 | 5.000 | 5.084 | 4.910 | 4.658 | 5.956 | 4.513 | 4.783 | 5.100 | 5.260 | 5.141 | 5.165 | 5.834 | 6.171 | 6.644 | 6.373 |
| 5.823 | 5.335 | 5.522 | 5.108 | 4.173 | 4.526 | 5.701 | 5.090 | 5.497 | 4.968 | 4.778 | 4.445 | 5.768 | 4.583 | 4.207 | 5.308 |
| 6.626 | 6.026 | 5.738 | 5.960 | 5.136 | 5.989 | 6.367 | 6.045 | 6.784 | 6.121 | 6.347 | 5.904 | 7.318 | 5.671 | 6.612 | 7.516 |
| 5.859 | 5.695 | 5.683 | 5.604 | 5.148 | 6.262 | 5.789 | 5.697 | 6.220 | 6.213 | 5.968 | 5.779 | 6.531 | 6.148 | 6.518 | 6.539 |
| 7.015 | 6.727 | 6.155 | 6.568 | 5.557 | 6.470 | 6.405 | 6.272 | 7.271 | 6.883 | 6.672 | 6.265 | 7.841 | 6.281 | 6.957 | 7.511 |
| 5.631 | 5.395 | 5.347 | 5.299 | 4.986 | 5.662 | 5.214 | 5.278 | 5.790 | 5.605 | 5.540 | 5.351 | 5.999 | 5.761 | 5.909 | 6.079 |
| 6.209 | 6.434 | 6.683 | 6.341 | 5.486 | 6.965 | 5.712 | 5.994 | 6.559 | 6.444 | 6.216 | 6.167 | 7.057 | 7.046 | 6.970 | 7.529 |
| 6.252 | 6.241 | 6.050 | 6.131 | 5.400 | 6.891 | 6.183 | 6.115 | 7.193 | 6.776 | 6.369 | 6.195 | 7.607 | 6.875 | 7.563 | 7.388 |

Supplemental Table 3A. Disease free survival in all peptides showing significant change from DCIS with no later event to IBC later event independent of laterality. HR- Cox Hazard Ratio. * Note: all calculations given are age-adjusted

| Event | Peak (m/z) | HR (Low/High) | 95% CI [lower-upper] | P, surv(high) | P, surv(low) | p-value |
|---|---|---|---|---|---|---|
| IBC | 716.336 | 2.538 | [1.159-5.556] | 0.8543 | 0.6791 | 0.0198 |
| IBC | 741.355 | 2.088 | [0.968-4.505] | 0.8834 | 0.6822 | 0.0605 |
| IBC | 770.415 | 2.267 | [1.038-4.950] | 0.8843 | 0.6817 | 0.0399 |
| IBC | 795.411 | 2.475 | [1.126-5.435] | 0.8946 | 0.6702 | 0.0242 |
| IBC | 971.454 | 2.475 | [1.125-5.435] | 0.8943 | 0.6675 | 0.0243 |
| IBC | 986.471 | 2.381 | [1.086-5.236] | 0.8988 | 0.6714 | 0.0303 |
| IBC | 1034.572 | 1.980 | [0.923-4.255] | 0.8811 | 0.6841 | 0.0794 |
| IBC | 1055.556 | 2.695 | [1.229-5.917] | 0.8545 | 0.6749 | 0.0133 |
| IBC | 1068.508 | 2.000 | [0.933-4.292] | 0.8815 | 0.6808 | 0.075 |
| IBC | 1084.493 | 2.247 | [1.045-4.831] | 0.8195 | 0.7266 | 0.0382 |
| IBC | 1090.491 | 2.415 | [1.091-5.348] | 0.8954 | 0.6771 | 0.0297 |
| IBC | 1350.638 | 2.387 | [1.089-5.208] | 0.8996 | 0.6552 | 0.0296 |
| IBC | 1416.604 | 2.985 | [1.326-6.711] | 0.8621 | 0.6707 | 0.0083 |
| IBC | 1635.783 | 2.604 | [1.186-5.714] | 0.8498 | 0.6845 | 0.017 |
| IBC | 1921.895 | 2.409 | [1.094-5.319] | 0.8991 | 0.6657 | 0.0291 |

Supplemental Table 3B. Disease free survival in all peptides showing significant change from DCIS with no later event to IBC later event ipsilateral. HR- Cox Hazard Ratio. * Note: all calculations given are age-adjusted

| Event | Peak (m/z) | HR (Low/High) | 95% CI [lower-upper] | P, surv(high) | P, surv(low) | P-value |
|---|---|---|---|---|---|---|
| Ipsilateral IBC | 716.336 | 2.833 | [1.006-7.937] | 0.8543 | 0.6791 | 0.0488 |
| Ipsilateral IBC | 741.355 | 2.114 | [0.785-5.682] | 0.8834 | 0.6822 | 0.1388 |
| Ipsilateral IBC | 770.415 | 1.538 | [0.596-3.968] | 0.8843 | 0.6817 | 0.3737 |
| Ipsilateral IBC | 795.411 | 2.786 | [0.986-7.874] | 0.9429 | 0.7729 | 0.0532 |
| Ipsilateral IBC | 971.454 | 2.801 | [0.989-7.937] | 0.9433 | 0.7698 | 0.0525 |
| Ipsilateral IBC | 986.471 | 2.674 | [0.948-7.576] | 0.8988 | 0.6714 | 0.063 |
| Ipsilateral IBC | 1034.572 | 1.580 | [0.610-4.098] | 0.8811 | 0.6841 | 0.3463 |
| Ipsilateral IBC | 1055.556 | 2.320 | [0.867-6.211] | 0.8545 | 0.6749 | 0.0939 |
| Ipsilateral IBC | 1068.508 | 2.024 | [0.757-5.405] | 0.8815 | 0.6808 | 0.1602 |
| Ipsilateral IBC | 1084.493 | 1.761 | [0.678-4.587] | 0.8195 | 0.7266 | 0.2455 |
| Ipsilateral IBC | 1090.491 | 2.710 | [0.950-7.752] | 0.8954 | 0.6771 | 0.0625 |
| Ipsilateral IBC | 1350.638 | 1.618 | [0.626-4.184] | 0.8996 | 0.6552 | 0.3205 |
| Ipsilateral IBC | 1416.604 | 2.841 | [1.010-8.000] | 0.9442 | 0.7598 | 0.0478 |
| Ipsilateral IBC | 1635.783 | 3.906 | [1.282-11.905] | 0.8498 | 0.6845 | 0.0166 |
| Ipsilateral IBC | 1921.895 | 1.631 | [0.625-4.255] | 0.8991 | 0.6657 | 0.3176 |

Supplemental Table 4A. Disease free survival in all peptides showing significant change from DCIS with no later event to later event of DCIS independent of laterality. HR- Cox Hazard Ratio. * Note: all calculations given are age-adjusted

| Event | Peak (m/z) | HR (Low/High) | 95% CI [lower-upper] | P, surv(high) | P, surv(low) | p-value |
|---|---|---|---|---|---|---|
| DCIS | 716.336 | 1.217 | [0.439-1.539] | 0.7337 | 0.6087 | 0.5393 |
| DCIS | 741.355 | 1.730 | [0.304-1.098] | 0.7659 | 0.5956 | 0.0941 |
| DCIS | 770.415 | 0.798 | [0.668-2.350] | 0.6623 | 0.6901 | 0.4827 |
| DCIS | 795.411 | 1.252 | [0.425-1.503] | 0.6939 | 0.6413 | 0.4865 |
| DCIS | 971.454 | 1.227 | [0.432-1.535] | 0.7329 | 0.6224 | 0.5259 |
| DCIS | 986.471 | 1.258 | [0.425-1.486] | 0.69 | 0.6546 | 0.4717 |
| DCIS | 1034.572 | 1.198 | [0.466-1.563] | 0.7285 | 0.6224 | 0.5727 |
| DCIS | 1055.556 | 1.582 | [0.337-1.185] | 0.724 | 0.6162 | 0.1528 |
| DCIS | 1068.508 | 1.099 | [0.487-1.702] | 0.6926 | 0.6503 | 0.7682 |
| DCIS | 1084.493 | 1.346 | [0.397-1.390] | 0.7373 | 0.606 | 0.3525 |
| DCIS | 1090.491 | 1.115 | [0.472-1.704] | 0.6698 | 0.6841 | 0.7395 |
| DCIS | 1350.638 | 1.529 | [0.349-1.224] | 0.7259 | 0.623 | 0.1838 |
| DCIS | 1416.604 | 1.325 | [0.404-1.411] | 0.7368 | 0.6102 | 0.3783 |
| DCIS | 1635.783 | 1.550 | [0.344-1.210] | 0.7015 | 0.6384 | 0.1719 |
| DCIS | 1921.895 | 1.34 | [0.397-1.402] | 0.7025 | 0.6482 | 0.3632 |

Supplemental Table 4B. Disease free survival in all peptides showing significant change from DCIS with no later event to later event of ipsilateral DCIS. HR- Cox Hazard Ratio. * Note: all calculations given are age-adjusted

| Event | Peak (m/z) | HR (Low/High) | 95% CI [lower-upper] | P, surv(high) | P, surv(low) | p-value |
|---|---|---|---|---|---|---|
| Ipsilateral DCIS | 716.336 | 1.119 | [0.367-2.176] | 0.8733 | 0.8518 | 0.8049 |
| Ipsilateral DCIS | 741.355 | 0.927 | [0.432-2.699] | 0.8579 | 0.8672 | 0.8705 |
| Ipsilateral DCIS | 770.415 | 0.596 | [0.669-4.209] | 0.8262 | 0.9034 | 0.2701 |
| Ipsilateral DCIS | 795.411 | 1.149 | [0.355-2.130] | 0.8713 | 0.8524 | 0.7606 |
| Ipsilateral DCIS | 971.454 | 0.923 | [0.435-2.696] | 0.857 | 0.8696 | 0.8636 |
| Ipsilateral DCIS | 986.471 | 1.302 | [0.318-1.858] | 0.8806 | 0.8438 | 0.5585 |
| Ipsilateral DCIS | 1034.572 | 1.030 | [0.398-2.367] | 0.8661 | 0.863 | 0.948 |
| Ipsilateral DCIS | 1055.556 | 1.398 | [0.295-1.734] | 0.8848 | 0.8326 | 0.4583 |
| Ipsilateral DCIS | 1068.508 | 0.821 | [0.495-2.996] | 0.8575 | 0.8716 | 0.6687 |
| Ipsilateral DCIS | 1084.493 | 0.907 | [0.446-2.722] | 0.8503 | 0.8784 | 0.8339 |
| Ipsilateral DCIS | 1090.491 | 0.924 | [0.433-2.705] | 0.8576 | 0.8714 | 0.8662 |
| Ipsilateral DCIS | 1350.638 | 1.272 | [0.325-1.896] | 0.8805 | 0.8435 | 0.5914 |
| Ipsilateral DCIS | 1416.604 | 1.115 | [0.369-2.182] | 0.8632 | 0.8618 | 0.8111 |
| Ipsilateral DCIS | 1635.783 | 1.425 | [0.289-1.703] | 0.8741 | 0.849 | 0.4335 |
| Ipsilateral DCIS | 1921.895 | 1.113 | [0.359-2.153] | 0.8687 | 0.8594 | 0.7777 |

Supplemental Figure 1A. Disease free survival in all peptides showing significant change from primary DCIS to IBC recurrence independent of laterality.

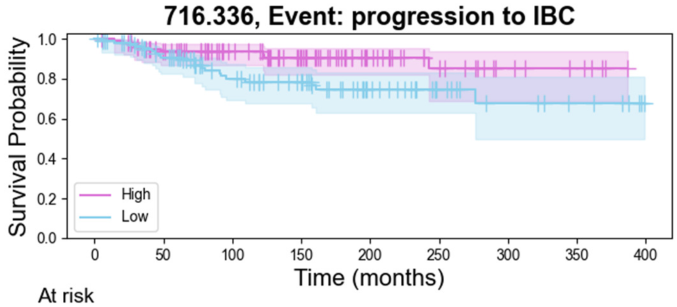


| At risk | | | | | | | | | |
|---|---|---|---|---|---|---|---|---|---|
| High | 103 | 85 | 65 | 49 | 27 | 16 | 8 | 5 | 0 |
| Low | 103 | 82 | 58 | 46 | 29 | 16 | 9 | 6 | 0 |

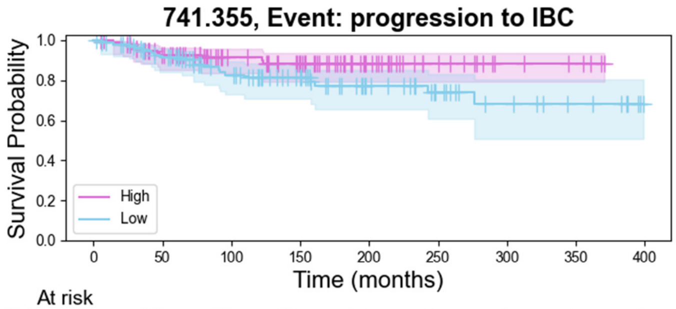


| At risk | | | | | | | | | |
|---|---|---|---|---|---|---|---|---|---|
| High | 103 | 83 | 61 | 50 | 27 | 14 | 6 | 4 | 0 |
| Low | 103 | 84 | 62 | 45 | 29 | 18 | 11 | 7 | 0 |

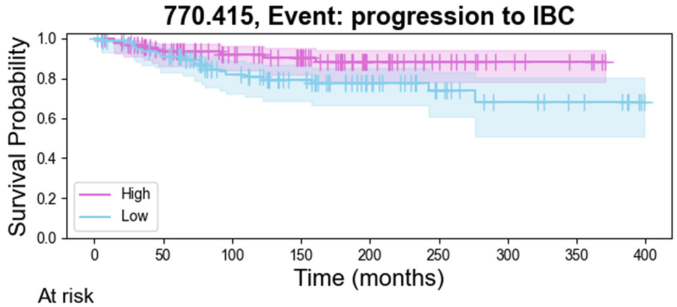


| At risk | | | | | | | | | |
|---|---|---|---|---|---|---|---|---|---|
| High | 103 | 80 | 62 | 48 | 26 | 15 | 7 | 4 | 0 |
| Low | 103 | 87 | 61 | 47 | 30 | 17 | 10 | 7 | 0 |

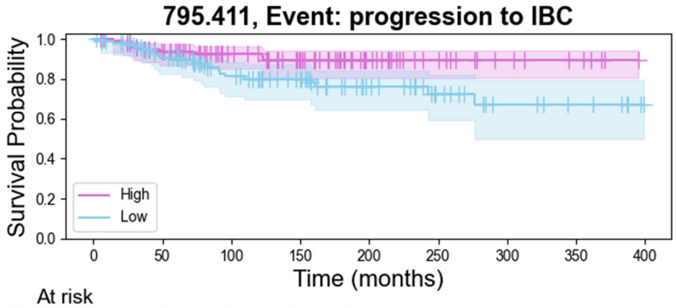


| At risk | | | | | | | | | |
|---|---|---|---|---|---|---|---|---|---|
| High | 103 | 85 | 63 | 49 | 27 | 15 | 8 | 5 | 0 |
| Low | 103 | 82 | 60 | 46 | 29 | 17 | 9 | 6 | 0 |

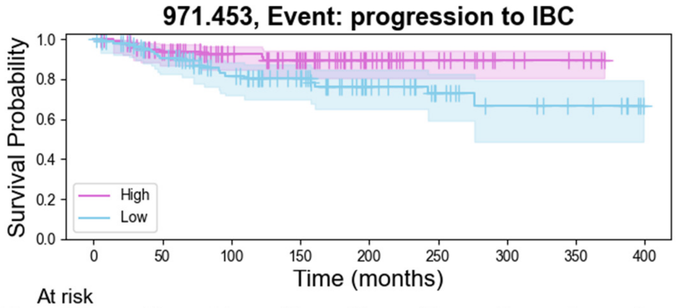


| At risk | | | | | | | | | |
|---|---|---|---|---|---|---|---|---|---|
| High | 103 | 84 | 61 | 49 | 28 | 16 | 7 | 4 | 0 |
| Low | 103 | 83 | 62 | 46 | 28 | 16 | 10 | 7 | 0 |

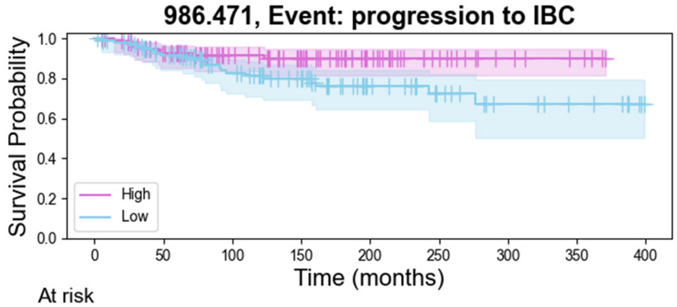


| At risk | | | | | | | | | |
|---|---|---|---|---|---|---|---|---|---|
| High | 103 | 84 | 61 | 48 | 28 | 15 | 7 | 4 | 0 |
| Low | 103 | 83 | 62 | 47 | 28 | 17 | 10 | 7 | 0 |

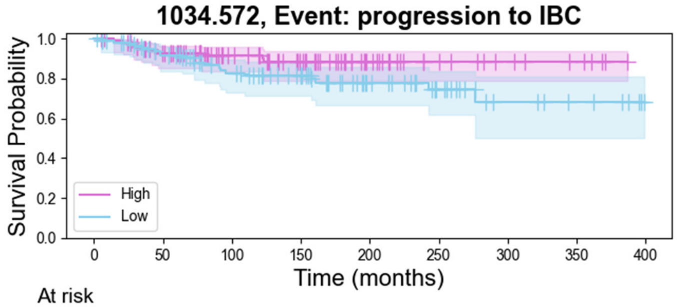


| At risk | | | | | | | | | |
|---|---|---|---|---|---|---|---|---|---|
| High | 103 | 84 | 59 | 44 | 24 | 13 | 8 | 5 | 0 |
| Low | 103 | 83 | 64 | 51 | 32 | 19 | 9 | 6 | 0 |

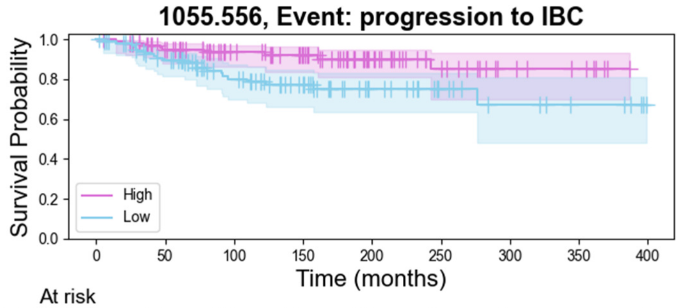


| At risk | | | | | | | | | |
|---|---|---|---|---|---|---|---|---|---|
| High | 103 | 85 | 66 | 55 | 32 | 18 | 9 | 6 | 0 |
| Low | 103 | 82 | 57 | 40 | 24 | 14 | 8 | 5 | 0 |

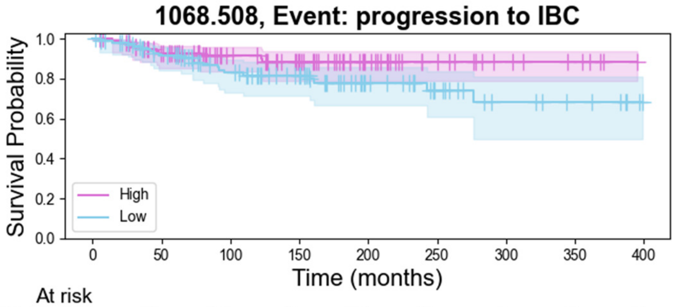


| At risk | | | | | | | | | |
|---|---|---|---|---|---|---|---|---|---|
| High | 103 | 83 | 59 | 46 | 27 | 15 | 8 | 5 | 0 |
| Low | 103 | 84 | 64 | 49 | 29 | 17 | 9 | 6 | 0 |

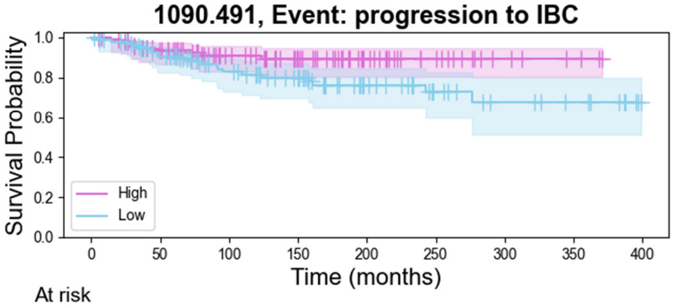


| At risk | | | | | | | | | |
|---|---|---|---|---|---|---|---|---|---|
| High | 103 | 84 | 60 | 46 | 26 | 15 | 6 | 3 | 0 |

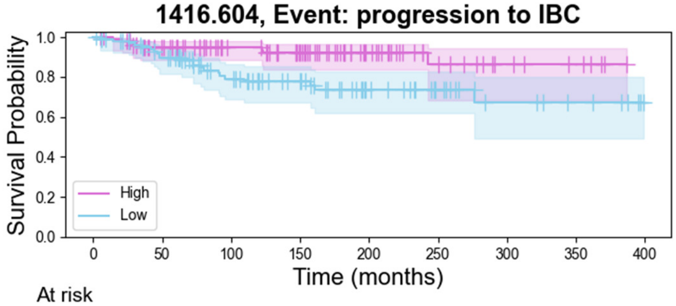


| At risk | | | | | | | | | |
|---|---|---|---|---|---|---|---|---|---|
| High | 103 | 84 | 65 | 52 | 29 | 15 | 8 | 5 | 0 |
| Low | 103 | 83 | 58 | 43 | 27 | 17 | 9 | 6 | 0 |

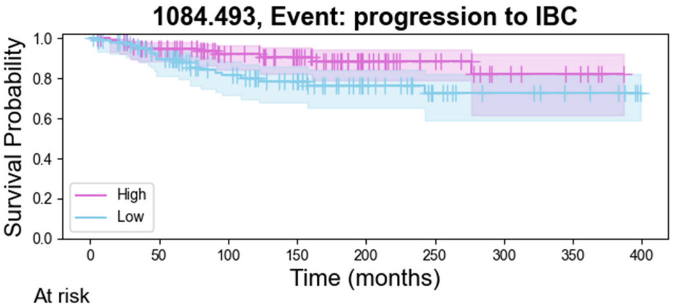


| At risk | | | | | | | | | |
|---|---|---|---|---|---|---|---|---|---|
| High | 103 | 86 | 64 | 51 | 28 | 17 | 8 | 5 | 0 |
| Low | 103 | 81 | 59 | 44 | 28 | 15 | 9 | 6 | 0 |

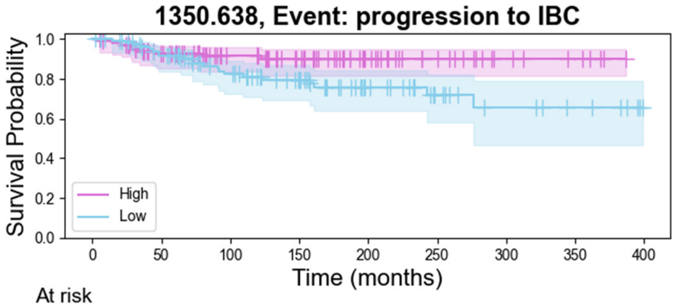


| At risk | | | | | | | | | |
|---|---|---|---|---|---|---|---|---|---|
| High | 103 | 83 | 60 | 48 | 28 | 17 | 8 | 5 | 0 |
| Low | 103 | 84 | 63 | 47 | 28 | 15 | 9 | 6 | 0 |

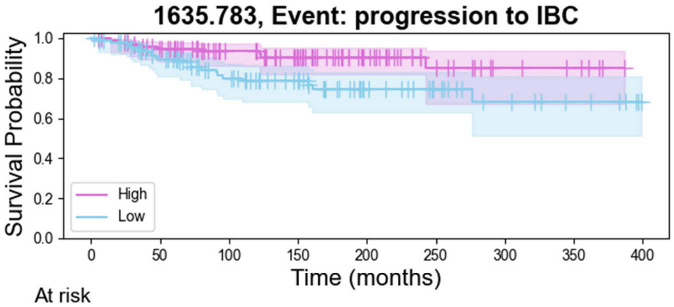


| At risk | | | | | | | | | |
|---|---|---|---|---|---|---|---|---|---|
| High | 103 | 87 | 65 | 52 | 30 | 15 | 7 | 5 | 0 |
| Low | 103 | 80 | 58 | 43 | 26 | 17 | 10 | 6 | 0 |

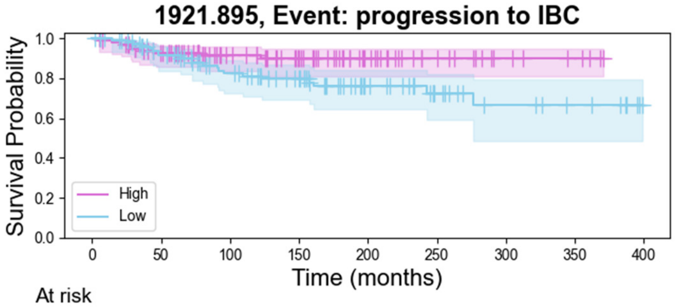


| At risk | | | | | | | | | |
|---|---|---|---|---|---|---|---|---|---|
| High | 103 | 85 | 60 | 46 | 27 | 15 | 7 | 4 | 0 |
| Low | 103 | 82 | 63 | 49 | 29 | 17 | 10 | 7 | 0 |

Supplemental Figure 1B. Disease free survival in all peptides showing significant change from primary DCIS to IBC recurrence independent of laterality.

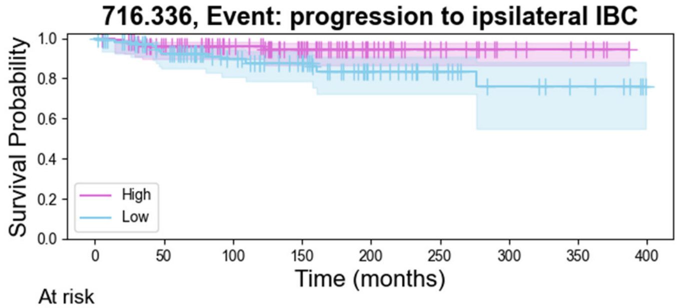


| At risk | | | | | | | | | |
|---|---|---|---|---|---|---|---|---|---|
| High | 103 | 85 | 65 | 49 | 27 | 16 | 8 | 5 | 0 |
| Low | 103 | 82 | 58 | 46 | 29 | 16 | 9 | 6 | 0 |

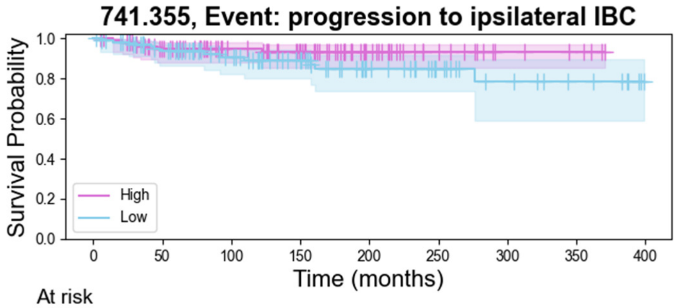


| At risk | | | | | | | | | |
|---|---|---|---|---|---|---|---|---|---|
| High | 103 | 83 | 61 | 50 | 27 | 14 | 6 | 4 | 0 |
| Low | 103 | 84 | 62 | 45 | 29 | 18 | 11 | 7 | 0 |

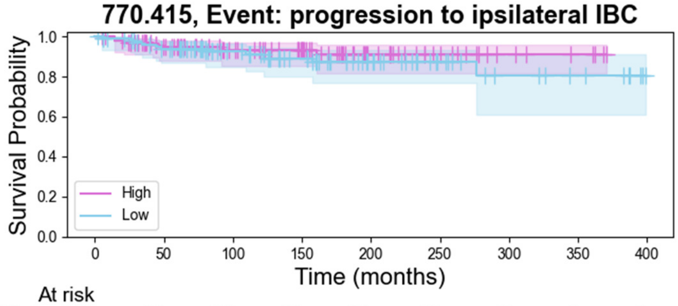


| At risk | | | | | | | | | |
|---|---|---|---|---|---|---|---|---|---|
| High | 103 | 80 | 62 | 48 | 26 | 15 | 7 | 4 | 0 |
| Low | 103 | 87 | 61 | 47 | 30 | 17 | 10 | 7 | 0 |

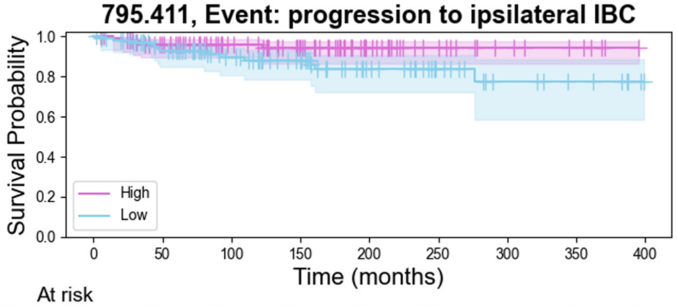


| At risk | | | | | | | | | |
|---|---|---|---|---|---|---|---|---|---|
| High | 103 | 85 | 63 | 49 | 27 | 15 | 8 | 5 | 0 |
| Low | 103 | 82 | 60 | 46 | 29 | 17 | 9 | 6 | 0 |

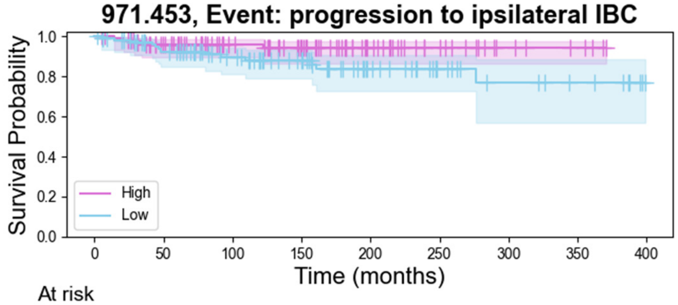


| At risk | | | | | | | | | |
|---|---|---|---|---|---|---|---|---|---|
| High | 103 | 84 | 61 | 49 | 28 | 16 | 7 | 4 | 0 |
| Low | 103 | 83 | 62 | 46 | 28 | 16 | 10 | 7 | 0 |

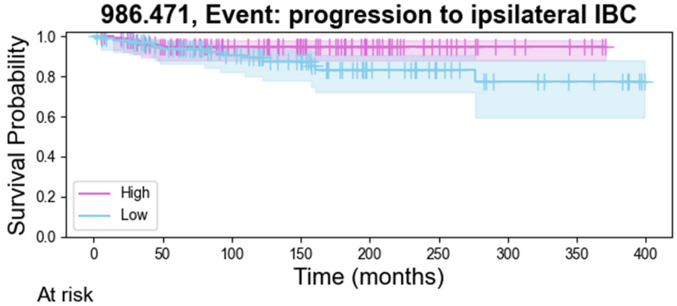


| At risk | | | | | | | | | |
|---|---|---|---|---|---|---|---|---|---|
| High | 103 | 84 | 61 | 48 | 28 | 15 | 7 | 4 | 0 |
| Low | 103 | 83 | 62 | 47 | 28 | 17 | 10 | 7 | 0 |

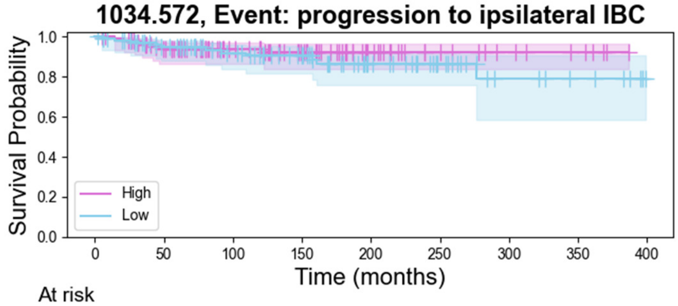


| At risk | | | | | | | | | |
|---|---|---|---|---|---|---|---|---|---|
| High | 103 | 84 | 59 | 44 | 24 | 13 | 8 | 5 | 0 |
| Low | 103 | 83 | 64 | 51 | 32 | 19 | 9 | 6 | 0 |

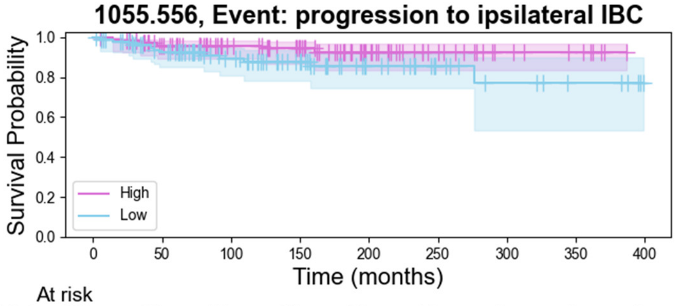


| At risk | | | | | | | | | |
|---|---|---|---|---|---|---|---|---|---|
| High | 103 | 85 | 66 | 55 | 32 | 18 | 9 | 6 | 0 |
| Low | 103 | 82 | 57 | 40 | 24 | 14 | 8 | 5 | 0 |

Supplemental Figure 1C. Disease free survival in all peptides showing significant change from primary DCIS to IBC recurrence independent of ipsalateral

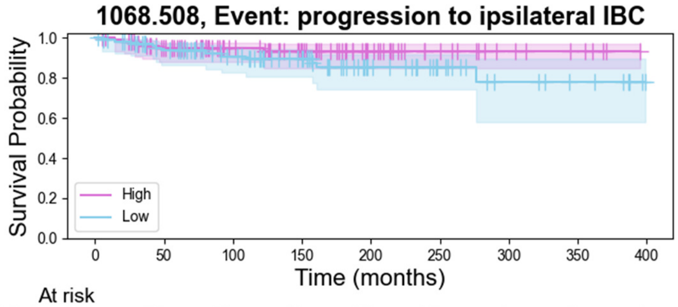


| At risk | | | | | | | | | |
|---|---|---|---|---|---|---|---|---|---|
| High | 103 | 83 | 59 | 46 | 27 | 15 | 8 | 5 | 0 |
| Low | 103 | 84 | 64 | 49 | 29 | 17 | 9 | 6 | 0 |

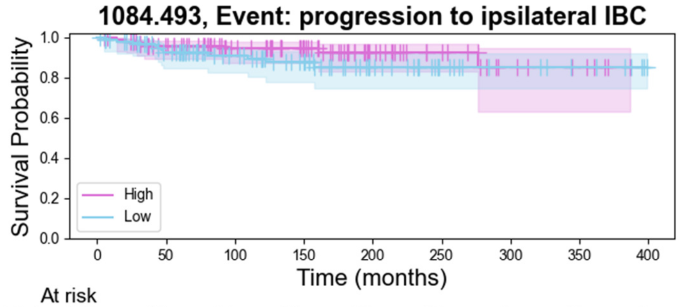


| At risk | | | | | | | | | |
|---|---|---|---|---|---|---|---|---|---|
| High | 103 | 86 | 64 | 51 | 28 | 17 | 8 | 5 | 0 |
| Low | 103 | 81 | 59 | 44 | 28 | 15 | 9 | 6 | 0 |

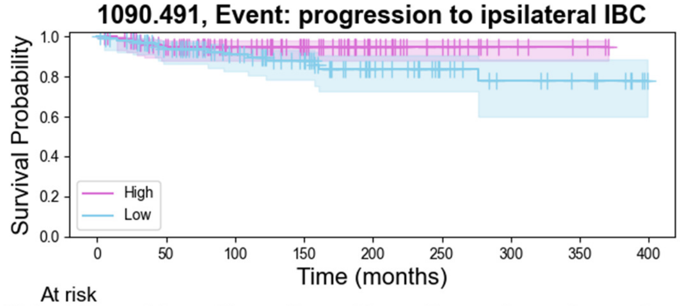


| At risk | | | | | | | | | |
|---|---|---|---|---|---|---|---|---|---|
| High | 103 | 84 | 60 | 46 | 26 | 15 | 6 | 3 | 0 |
| Low | 103 | 83 | 63 | 49 | 30 | 17 | 11 | 8 | 0 |

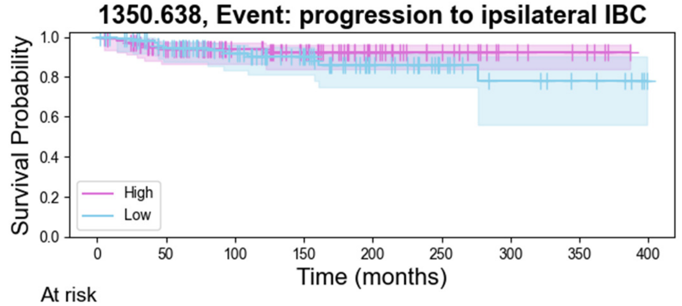


| At risk | | | | | | | | | |
|---|---|---|---|---|---|---|---|---|---|
| High | 103 | 83 | 60 | 48 | 28 | 17 | 8 | 5 | 0 |
| Low | 103 | 84 | 63 | 47 | 28 | 15 | 9 | 6 | 0 |

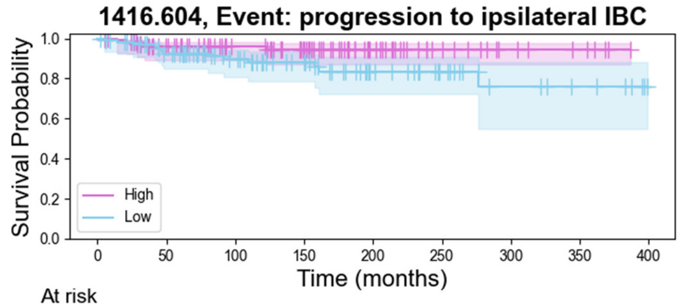


| At risk | | | | | | | | | |
|---|---|---|---|---|---|---|---|---|---|
| High | 103 | 84 | 65 | 52 | 29 | 15 | 8 | 5 | 0 |
| Low | 103 | 83 | 58 | 43 | 27 | 17 | 9 | 6 | 0 |

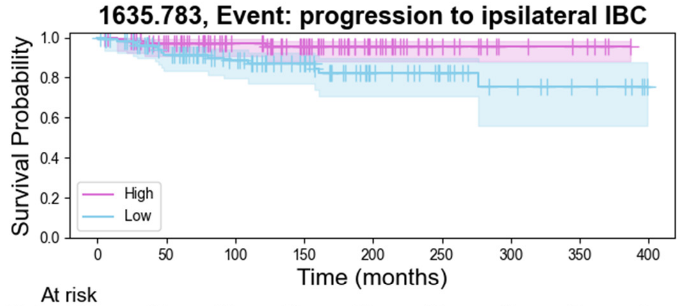


| At risk | | | | | | | | | |
|---|---|---|---|---|---|---|---|---|---|
| High | 103 | 87 | 65 | 52 | 30 | 15 | 7 | 5 | 0 |
| Low | 103 | 80 | 58 | 43 | 26 | 17 | 10 | 6 | 0 |

Supplemental Figure 1D. Disease free survival in all peptides showing significant change from primary DCIS to IBC recurrence independent of ipsalateral

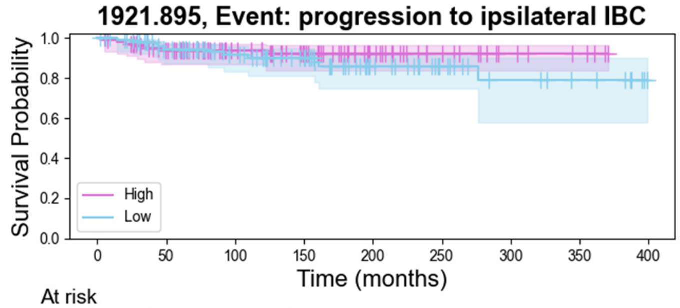


| At risk | | | | | | | | | |
|---|---|---|---|---|---|---|---|---|---|
| High | 103 | 85 | 60 | 46 | 27 | 15 | 7 | 4 | 0 |
| Low | 103 | 82 | 63 | 49 | 29 | 17 | 10 | 7 | 0 |

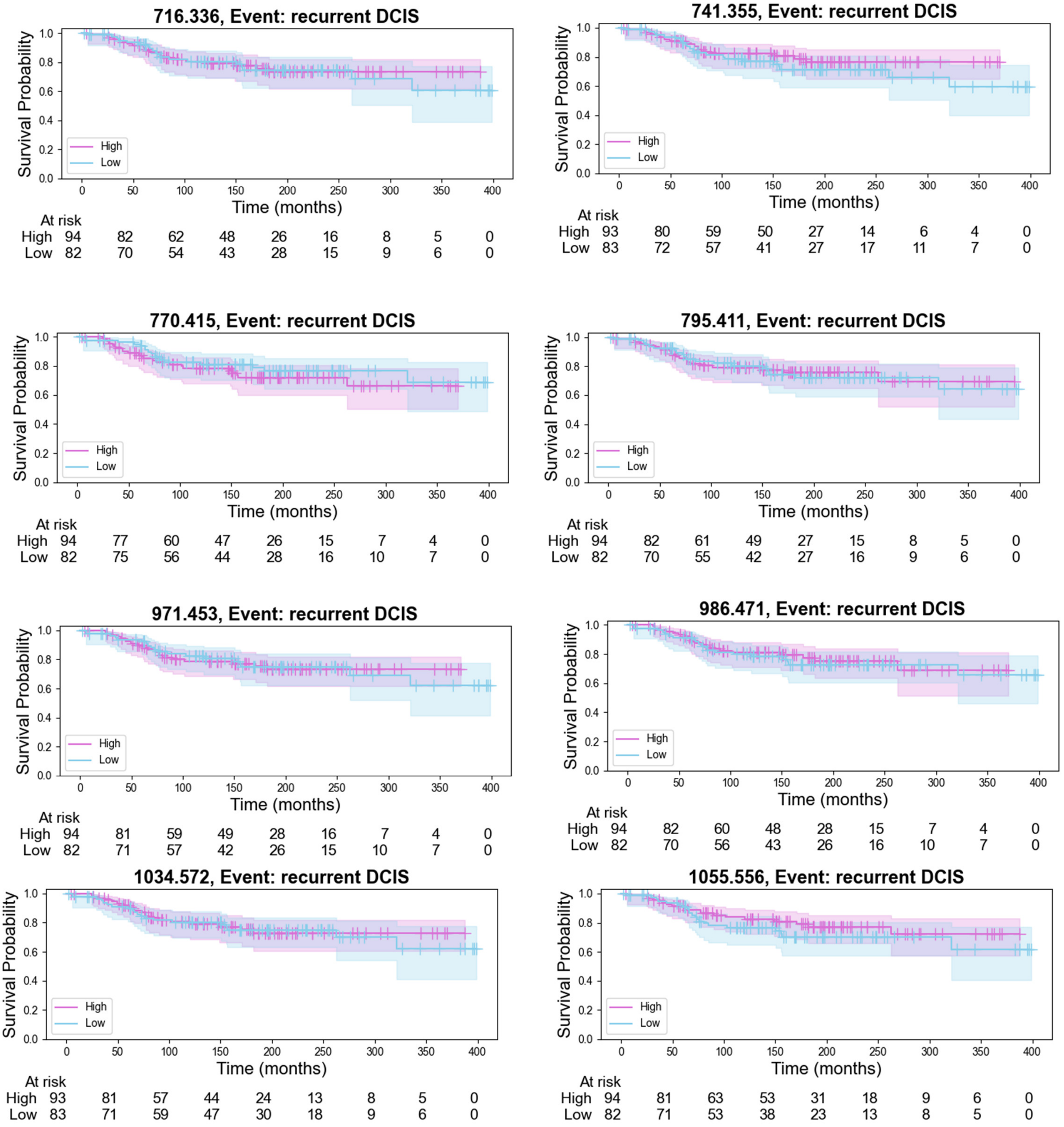


Supplemental Figure 1E. Disease free survival in all peptides showing significant change from primary DCIS to DCIS recurrence independent of laterality.

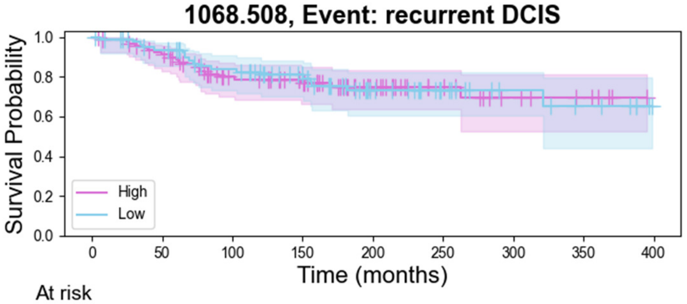


| At risk | | | | | | | | | |
|---|---|---|---|---|---|---|---|---|---|
| High | 93 | 80 | 57 | 46 | 27 | 15 | 8 | 5 | 0 |
| Low | 83 | 72 | 59 | 45 | 27 | 16 | 9 | 6 | 0 |

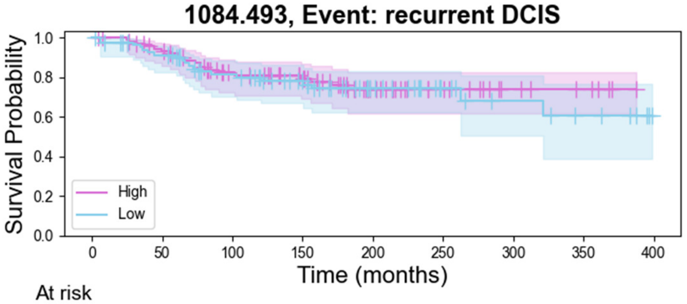


| At risk | | | | | | | | | |
|---|---|---|---|---|---|---|---|---|---|
| High | 93 | 81 | 61 | 49 | 27 | 16 | 8 | 5 | 0 |
| Low | 83 | 71 | 55 | 42 | 27 | 15 | 9 | 6 | 0 |

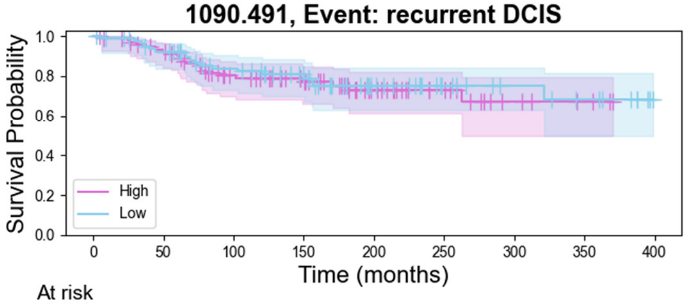


| At risk | | | | | | | | | |
|---|---|---|---|---|---|---|---|---|---|
| High | 94 | 81 | 59 | 46 | 26 | 15 | 6 | 3 | 0 |
| Low | 82 | 71 | 57 | 45 | 28 | 16 | 11 | 8 | 0 |

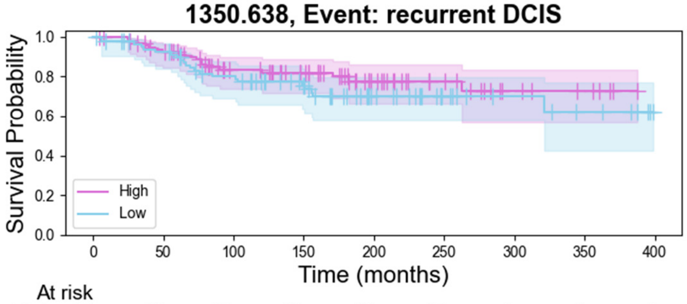


| At risk | | | | | | | | | |
|---|---|---|---|---|---|---|---|---|---|
| High | 94 | 81 | 59 | 48 | 28 | 17 | 8 | 5 | 0 |
| Low | 82 | 71 | 57 | 43 | 26 | 14 | 9 | 6 | 0 |

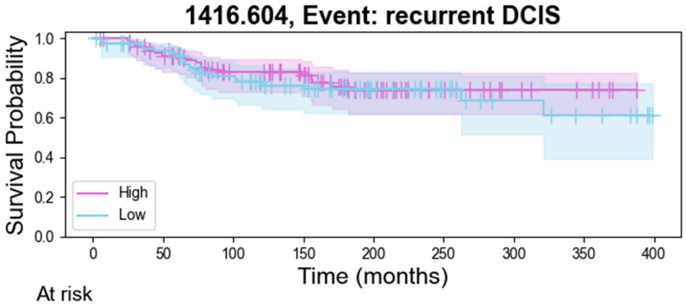


| At risk | | | | | | | | | |
|---|---|---|---|---|---|---|---|---|---|
| High | 95 | 81 | 62 | 51 | 28 | 15 | 8 | 5 | 0 |
| Low | 81 | 71 | 54 | 40 | 26 | 16 | 9 | 6 | 0 |

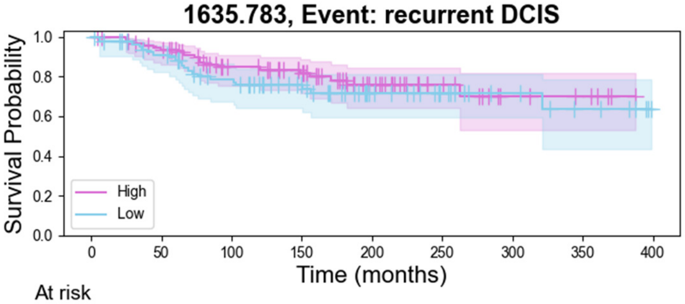


| At risk | | | | | | | | | |
|---|---|---|---|---|---|---|---|---|---|
| High | 94 | 83 | 62 | 51 | 29 | 15 | 7 | 5 | 0 |
| Low | 82 | 69 | 54 | 40 | 25 | 16 | 10 | 6 | 0 |

Supplemental Figure 1F. Disease free survival in all peptides showing significant change from primary DCIS to DCIS recurrence independent of laterality.

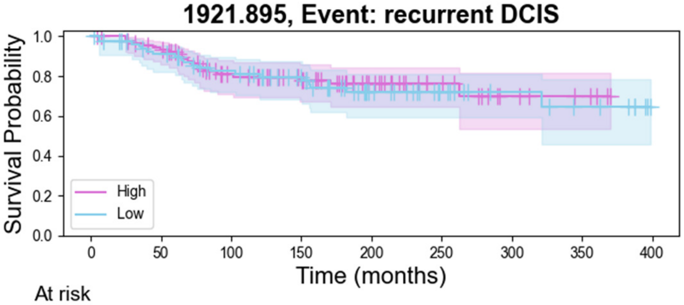


| At risk | | | | | | | | | |
|---|---|---|---|---|---|---|---|---|---|
| High | 94 | 83 | 59 | 46 | 27 | 15 | 7 | 4 | 0 |
| Low | 82 | 69 | 57 | 45 | 27 | 16 | 10 | 7 | 0 |

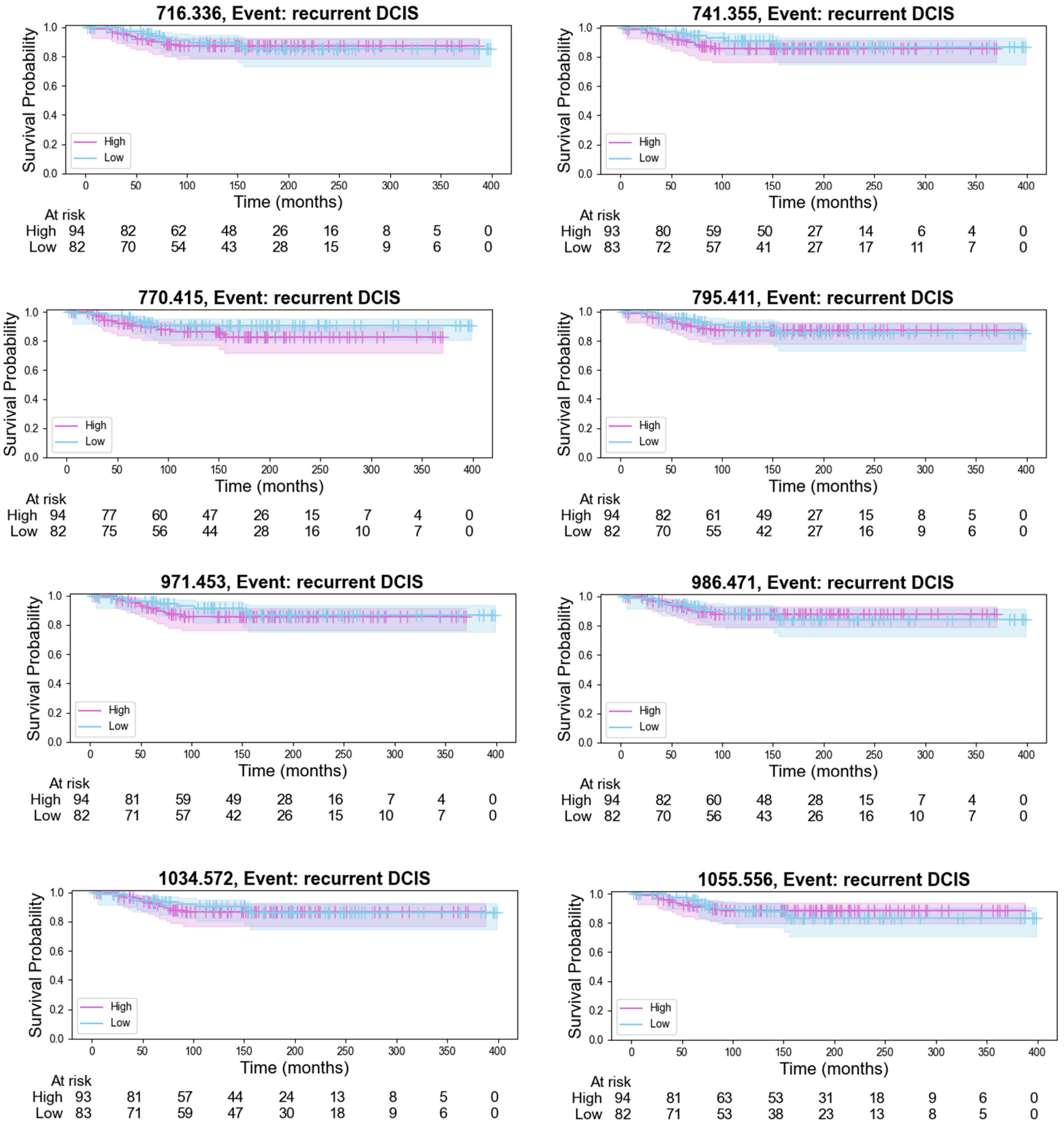


Supplemental Figure 1G. Disease free survival in all peptides showing significant change from primary DCIS to DCIS recurrence ipsilateral.

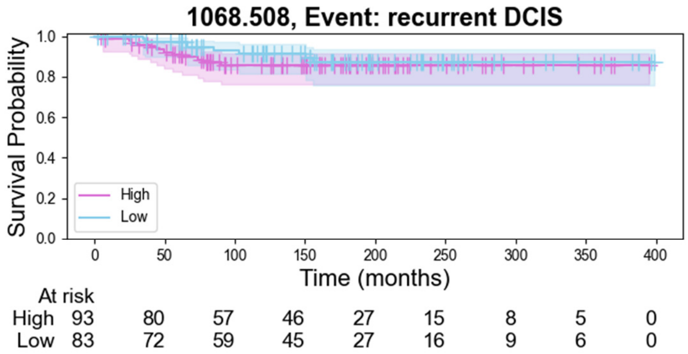


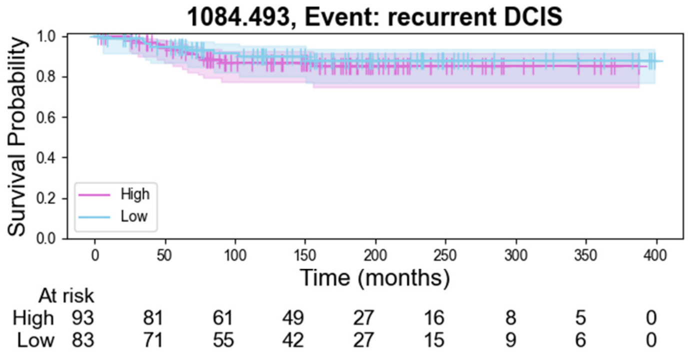


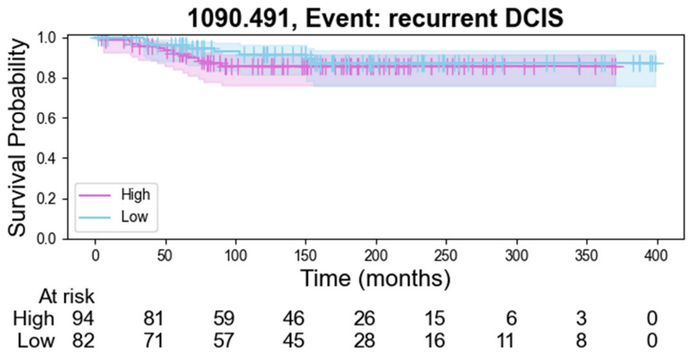


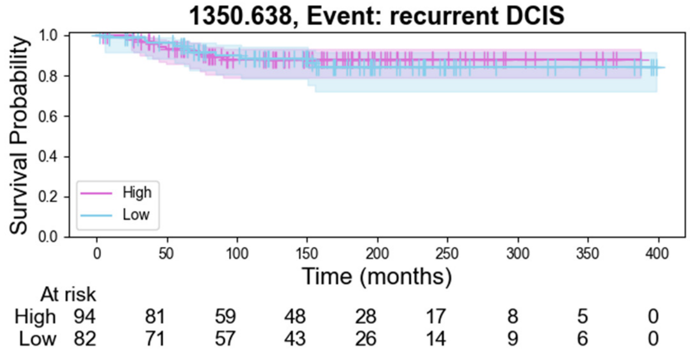


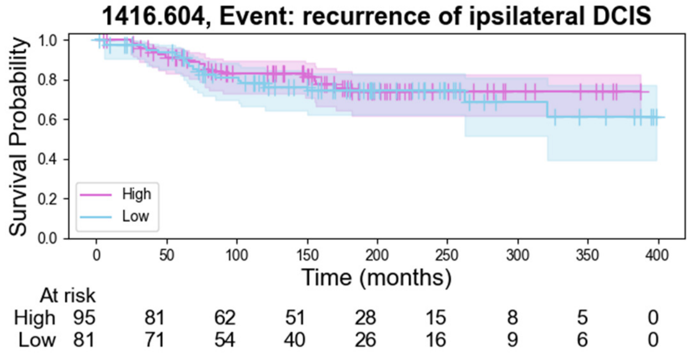


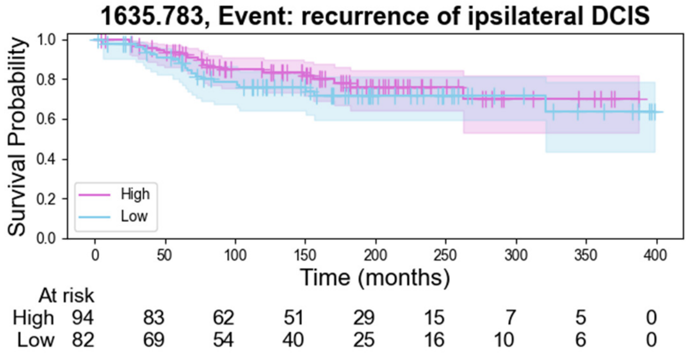


Supplemental Figure 1H. Disease free survival in all peptides showing significant change from primary DCIS to DCIS recurrence ipsilateral.

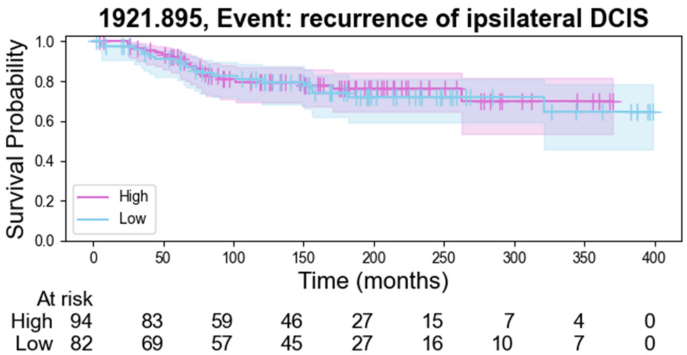

**A** **Model A, example six peptides**
P784.384, P903.502, P953.346,
P1084.493, P1154.616, P1364.581

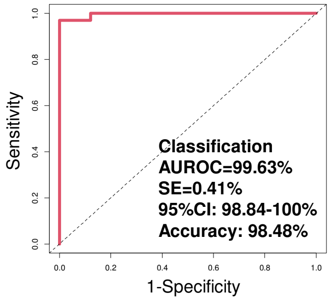


**B** **Model B, example ten peptides**
P743.370, P884.497, P903.502
P1006.414, P1032.469, P1102.398,
P1154.616, P1234.440, P1298.633,
P1364.581

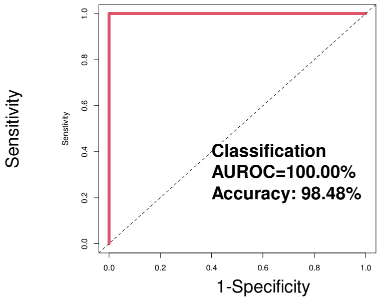


**C** **Model C, example eleven peptides**
P784.384, P840.325, P875.385,
P884.497, P903.502, P913.336,
P1084.493, P1199.449, P1234.440,
P1298.633, P1405.597

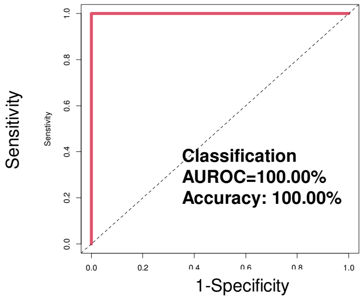


**D** **Model D, example fourteen peptides**
P743.370, P784.384, P875.385,
P884.497, P889.486, P903.502,
P913.336, P953.346, P1084.493,
P1154.512, P1199.449, P1234.440,
P1298.633, P1405.597

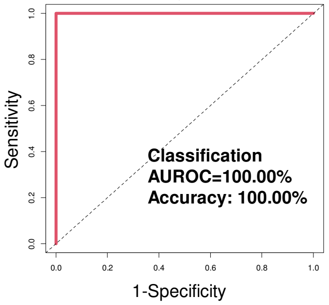


**E** **Model E, example twenty-one peptides**
P611.329, P726.341, P733.277, P741.355,
P743.370, P884.497, P903.502, P913.336,
P943.507, P953.346, P1032.469,
P1084.493, P1102.398, P1133.441,
P1234.440, P1241.484, P1273.536,
P1298.633, P1364.581,
P1405.597, P1508.600

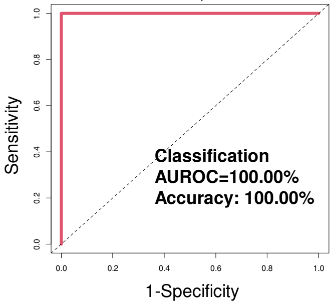


**F** **Exploratory Machine Learning Summary of LOOCV Statistics**

| | Model A | Model B | Model C | Model D | Model E |
|---|---|---|---|---|---|
| AUROC | 98.81% | 99.72% | 99.91% | 100.0% | 100.0% |
| SE | 1.22% | 0.32% | 0.13% | - | - |
| 95%CI | 96.41-100% | 99.11-100.0% | 99.65-100.0% | - | - |
| Accuracy | 98.48% | 96.97% | 96.97% | 98.48% | 93.38% |
| Sensitivity | 100.0% | 100.0% | 96.97% | 100.0% | 100.0% |
| Specificity | 96.97% | 93.94% | 96.97% | 96.97% | 96.97% |
| PPV | 97.06% | 94.29% | 96.97% | 97.06% | 97.06% |
| NPV | 100.0% | 100.0% | 96.97% | 100.0% | 100.0% |

**G** **Exploratory Machine Learning Summary of 3-Fold CV Statistics (N=600 AUROCs after 200 iterations)**

| | Model A | Model B | Model C | Model D | Model E |
|---|---|---|---|---|---|
| Mean AUROC | 99.25% | 99.62% | 99.69% | 99.85% | 99.77% |
| SE | 1.28% | 0.76% | 0.77% | 0.55% | 0.69% |
| Max AUROC | 100% | 100.0% | 100.0% | 100% | 100% |
| Min AUROC | 91.96% | 95.53% | 95.00% | 95.83% | 94.02% |

**Supplemental Figure 2.** Reports of all exploratory machine learning analysis identifies a peptide classification for recurrence on patient-matched samples of DCIS, or recurrent DCIS or IBC. **(A-E)** Support vector machine algorithms used to identify five different peptide classification models. **(F)** Summary of Leave-One-Out Cross Validation (LOOCV) statistics shown in table. **(G)** Summary of 3-Fold Cross Validation (CV) statistics shown in table. AUROC stands for area under the receiver-operator curve. SE stands for standard error. 95%CI stands for 95% confidence interval. PPV stands for positive predictive value and NPV for negative predictive value.
